# Rapid CAR screening and circRNA-driven CAR-NK cells for persistent shed-resistant immunotherapy

**DOI:** 10.1101/2025.08.19.671003

**Authors:** Joo-Yoon Chung, Jisu Hong, Su-Min Yee, Hye Won Lim, Dahye Lee, Sangwon Yoon, Seung-Hwan Lee, Yujin Jung, Seung Hun Shin, Song Cheol Kim, Chang-Han Lee, Mihue Jang

**Affiliations:** Medicinal Materials Research Center, Biomedical Research Division, Korea Institute of Science and Technology (KIST), Seoul 02792, Republic of Korea; KHU-KIST Department of Converging Science and Technology, Kyung Hee University, Seoul 02447, Republic of Korea; Department of Biomedical Sciences, Seoul National University College of Medicine, Seoul 03080, Republic of Korea; Cancer Research Institute, Seoul National University College of Medicine, Seoul 03080, Republic of Korea; Department of Pharmacology, Seoul National University College of Medicine, Seoul 03080, Republic of Korea; Corporate Research & Development Center, UCI Therapeutics, Seoul 04784, Republic of Korea; Department of Biochemistry, Microbiology and Immunology, Faculty of Medicine, University of Ottawa, Ottawa, Ontario K1N 6N5, Canada; Ottawa Institute of Systems Biology, Faculty of Medicine and Centre for Infection, Immunity, and Inflammation, Faculty of Medicine, University of Ottawa, Ontario K1N 6N5, Canada; Department of Biotechnology, College of Life Sciences and Biotechnology, Korea University, Seoul 02841, Republic of Korea; Division of Hepato-Biliary and Pancreatic Surgery, Department of Surgery, University of Ulsan College of Medicine, Asan Medical Center 05505, Seoul, Republic of Korea; BK21 FOUR Biomedical Science Project, Seoul National University College of Medicine, Seoul 03080, Republic of Korea; Wide River Institute of Immunology, Seoul National University, Hongcheon 25159, Republic of Korea; Division of Bio-Medical Science and Technology, KIST School, University of Science and Technology (UST), Seoul 02792, Republic of Korea

**Keywords:** Pancreatic cancer, Tumor microenvironment, Shed mesothelin, mRNA-CAR-NK, Circular RNA, IL-21

## Abstract

Chimeric antigen receptor (CAR)-based immunotherapies against solid tumors face two major hurdles, the “decoy effect” of shedding antigens that sequester CARs, and the limited persistence of immune effectors within the immunosuppressive tumor microenvironment. Here, we present a mechanistic approach to overcome these barriers by integrating a physiologically relevant screening platform with circular RNA (circRNA) engineering. Unlike conventional screens using immortalized cell lines, we performed rapid functional screening directly in human primary natural killer (NK) cells to identify a novel scFv, CLMS10. Structural modeling revealed that CLMS10 targets a membrane-proximal epitope that overlaps the proteolytic cleavage site, thereby evading inhibition by soluble mesothelin (solMSLN). Furthermore, we demonstrated that circRNA-mediated CAR expression, when codelivered with interleukin-21 (IL-21), confers sufficient stability to withstand the continuous antigen shedding induced by cancer-associated fibroblasts (CAFs), resulting in reduced CAR downregulation. In an *in vivo* metastatic pancreatic cancer model, IL-21-augmented circCAR-MS10-NK cells exhibited potency comparable to that of lentivirally engineered CAR-NK cells while offering superior manufacturability. Overall, this study establishes a paradigm for generating shed-resistant CAR therapeutics through the strategic integration of epitope-specific functional selection and enhanced RNA stability.

## Introduction

Natural killer (NK) cells are innate immune effectors that exert rapid and potent cytotoxicity against cancer and virus-infected cells without prior sensitization, making them promising platforms for chimeric antigen receptor (CAR)-based therapies.^1^ Unlike T cells, NK cells utilize a diverse array of germline-encoded receptors to recognize transformed cells, providing a layer of redundancy that enables target elimination even when antigen presentation mechanisms are impaired.^1–3^ This intrinsic versatility suggests that CAR-NK cells may be less vulnerable to immune escape mechanisms, such as MHC downregulation, that limit conventional T-cell therapies.^1–4^ CAR-NK cells integrate the antigen specificity of CAR constructs with the intrinsic cytotoxic mechanisms of NK cells, enhancing overall antitumor efficacy.^2^ Furthermore, activated NK cells can remodel the tumor microenvironment (TME) by secreting proinflammatory cytokines like IFN-γ, potentially promoting broader immune engagement in otherwise immunologically cold tumors.^3, 4^

While CAR-NK cells offer significant advantages over CAR-T cells, including a lower risk of graft-versus-host disease (GVHD) and cytokine release syndrome (CRS), their clinical translation is hindered by manufacturing challenges.^5–8^ Although CAR-NK cells are amenable to off-the-shelf allogeneic applications, they remain difficult to genetically modify and exhibit substantial donor-to-donor variability.^2, 9, 10^ Traditional viral transduction methods are limited by their low efficiency in NK cells and high cost.^9, 10^ More importantly, the conventional viral approach is ill-suited for the iterative, rapid functional optimization required to identify the best CAR candidates among a vast pool of antibodies. Viral workflows are slow and resource-intensive, limiting combinatorial screening of CAR design parameters (scFv epitope, hinge length and signaling domains) in relevant primary cells, and causing costly late-stage attrition when binders fail after CAR reformatting due to synapse-specific constraints.^9, 11, 12^ Consequently, there is a growing demand for a rapid, nonviral gene delivery platform that enables physiologically relevant functional screening directly in primary NK cells, ensuring that selected constructs retain potency in a clinical setting. A platform that can support fast design–build–test cycles in donor-derived NK cells would enable a more engineering-driven approach to CAR discovery, where functional output becomes the primary criterion rather than binding metrics alone.^13–15^

Mesothelin (MSLN) is a promising target for cancer immunotherapy because of its high expression in various malignancies, including mesothelioma, ovarian cancer, pancreatic adenocarcinoma, gastric adenocarcinoma, and triple-negative breast cancer, while its expression in normal tissues is limited.^16–18^ MSLN has been extensively pursued for various therapeutic modalities; however, elevated MSLN levels often correlate with aggressive clinical features and poor prognosis, partly due to its role in immune evasion.^19–21^ Despite their promise, MSLN-targeted strategies are severely compromised by “antigen shedding,” which releases soluble MSLN (solMSLN) that acts as a circulating sink, sequestering therapeutic agents and limiting productive engagement of tumor-associated MSLN.^22–25^ In the context of CAR therapies, this decoy phenomenon is particularly detrimental because soluble antigen binding can reduce effector activation without triggering productive cytotoxicity.^25, 26^ Specifically, first-generation therapeutic agents targeting membrane-distal epitopes (Region I), such as SS1P, have demonstrated limited efficacy due to susceptibility to the decoy effect.^23, 27^ To overcome this limitation, recent strategies have shifted toward membrane-proximal epitopes (Region III), as exemplified by hYP218 and 15B6,^25, 28^ which are retained on the cell surface after cleavage. Conceptually, targeting the membrane-proximal region should reduce susceptibility to soluble decoys.^25, 26, 28, 29^ Although theoretically superior, targeting this region is structurally challenging because antibodies must bind close to the cell membrane without steric hindrance while maintaining high affinity.^30, 31^ The steric constraints imposed by the glycocalyx and the specific geometry required for an optimal immune synapse mean that many proximal binders may fail to induce signaling.^12, 30–32^ Therefore, identifying a “shed-resistant” binder that functions optimally within the structural constraints of a CAR synapse requires rigorous validation beyond simple binding assays. The ideal binder must discriminate membrane-bound antigen from soluble forms while supporting stable synapse formation.^12, 25^

The performance of CAR-T and CAR-NK cell-based therapies is closely linked to the affinity, epitope specificity, and membrane proximity of the selected antibody or single-chain variable fragment (scFv).^11, 30, 33^ While high affinity is generally desired, excessive affinity can impair serial killing or exacerbate the antigen sink effect.^13, 34^ Antibodies reformatted into CARs often lose efficacy or fail to trigger proper immune synapses, necessitating iterative testing.^12, 35^ This “reformatting gap” is particularly acute in NK cells, where optimal signaling domains differ from those in T cells.^2, 35^ However, standard screening methods utilizing immortalized cell lines (e.g., NK-92) often fail to predict efficacy in donor-derived cells.^12, 35, 36^ Immortalized lines often differ in metabolic fitness and activation thresholds, leading to false-positive predictions for candidates that underperform in primary cells.^2, 12, 35^ To successfully exploit membrane-proximal targeting, an efficient screening method is essential to rapidly identify binders that are functionally optimized for the primary CAR-NK context under physiologically relevant conditions, including the presence of soluble antigen.^12, 13, 26, 29, 35^

Beyond antigen engagement, durability remains a critical hurdle. The TME imposes metabolic and suppressive barriers that limit NK cell persistence.^2, 37–39^ Furthermore, stromal components such as cancer-associated fibroblasts (CAFs) actively remodel the TME and modulate protease activity, thereby accelerating antigen shedding.^21, 40, 41^ A robust therapeutic strategy must therefore address not only the direct interaction with the tumor antigen but also the CAF-driven intensification of the decoy effect and the metabolic restrictions of the _TME._21, 37, 41

In this study, we address a central bottleneck in MSLN-directed CAR-NK therapy by identifying binders that remain functional under soluble antigen decoy pressure while sustaining activity in solid-tumor microenvironments. We establish a rapid, nonviral workflow for iterative functional screening of membrane-proximal CAR binders directly in primary NK cells and pair this with circRNA-based CAR expression. Together, this integrated strategy provides a scalable route to engineer CAR-NK candidates aligned with the biological constraints of MSLN shedding and the suppressive solid-tumor milieu.

## Results

### Selective scFv screening against mature MSLN (matMSLN) via the yeast display library system

To generate high-affinity antibodies against matMSLN (residues 296–606), 6-week-old BALB/c mice were immunized intraperitoneally with recombinant matMSLN protein **(Figs. 1 and 2a, and Supplementary Fig. 1).** Upon reaching a serum antibody titer of 1:100,000 (**Supplementary Fig. 2**), variable heavy (VH) and light (VL) chain genes were amplified from the spleen and bone marrow of sacrificed immunized mice, fused to their respective constant domains, and transformed into *Saccharomyces cerevisiae* strains JAR200 (MATa) and YVH10 (MATα) to construct heavy-chain (HC) and light-chain (LC) Fab libraries. Mating the HC and LC yeast libraries^42, 43^ yielded a combinatorial Fab repertoire (> 4[×[10^7^) **(Supplementary Fig. 3a, and Supplementary Table 1).** Flow cytometric selection of clones with high antigen-binding activity and subsequent cell-based panning of MSLN^+^ and MSLN^−^ AsPC-1 cells narrowed the candidates to 96 unique Fab clones **(Supplementary Figs. 3 and 4)**. Most antibody candidates exhibited strong matMSLN binding but negligible affinity for solMSLN (residues 296–592), indicating a favorable selectivity that may reduce interference from shed MSLN **(Fig. 2a, b).**^25, 44, 45^ To identify the most promising clones, we applied sequential selection criteria: retaining the top 90% of expressers to maximize recovery, confirming robust Fab assembly on the basis of diagonal distribution, and isolating the top 5% of matMSLN binders exhibiting preferential specificity for matMSLN over solMSLN. This systematic approach enabled the identification of antibody candidates with optimal expression and selectivity (**Fig. 2b, and Supplementary Fig. 4**).

**Fig. 1.**
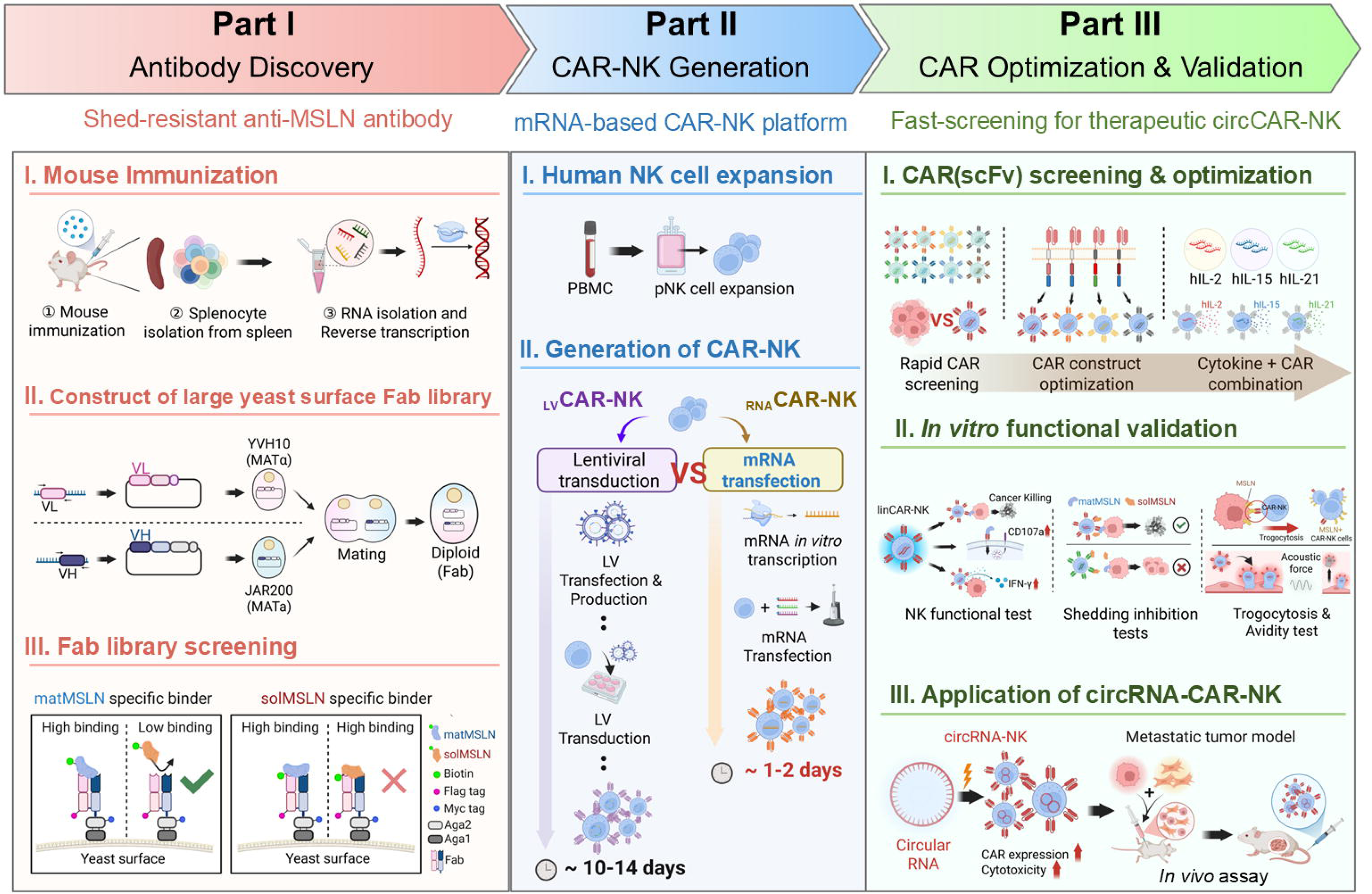
Streamlined chimeric antigen receptor (CAR) screening and circRNA-based CAR (circCAR)-natural killer (NK) cell development to overcome antigen shedding. Schematic workflow illustrating (1) antibody discovery via a yeast surface-display platform to isolate high-affinity clones targeting membrane-bound mesothelin (MSLN); (2) rapid CAR optimization via mRNA-engineering of NK cells for functional screening; and (3) development of persistent shed-resistant CAR-NK cells via circRNAs.

**Fig. 2.**
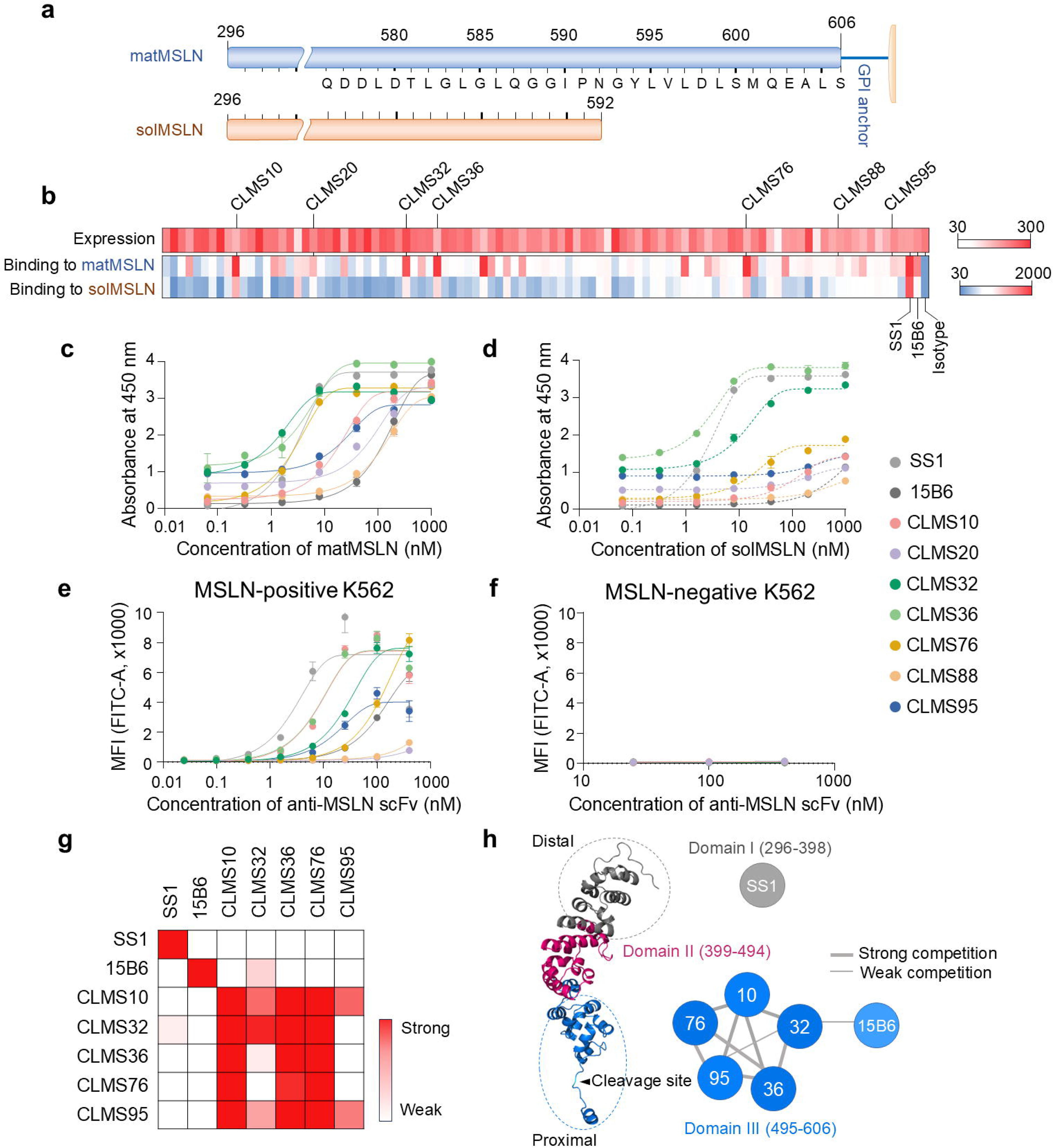
Screening and characterization of mature mesothelin (matMSLN)-specific scFvs. **(a)** Schematic representation of matMSLN (residues 296–606) and its soluble counterpart (solMSLN, residues 296–592). The top bar depicts the full-length matMSLN ending in a GPI anchor at residue 606; the bottom bar represents solMSLN, which lacks a C-terminal GPI-anchoring region. The corresponding amino acid sequence is aligned with the annotated residue positions. **(b)** Heatmap showing the levels and antigen-binding affinities of individual clones. Yeast-displayed clones were analyzed by flow cytometry using an anti-FLAG antibody (expression) and 10 nM biotinylated matMSLN or solMSLN (binding). SS1 and 15B6 were included as positive controls, and an isotype antibody served as a negative control. Clones are ranked by mean fluorescence intensity (MFI). **(c, d)** ELISA-based binding of recombinant anti-MSLN scFvs to matMSLN **(c)** and solMSLN **(d)**. Binding in immobilized scFvs incubated with serial dilutions of biotinylated antigen and detected via streptavidin-HRP. Note that the yeast-display binding values in **(b)** represent semiquantitative MFI-based screening at a fixed antigen concentration, whereas the ELISA results in **(c, d)** reflect the quantitative binding of the purified scFvs. Apparent differences between these assays arise from their distinct experimental formats, with ELISA providing a more accurate measure of solMSLN binding. **(e, f)** Flow cytometry of cell surface binding by scFvs to MSLN^+^ **(e)** and MSLN^−^ K562 cells **(f)**. Serially diluted antibodies were incubated with the cells, and the bound scFvs were detected via an anti-His-FITC secondary antibody. **(g)** Epitope binning of anti-MSLN scFvs (CLMS10, CLMS32, CLMS36, CLMS76, CLMS95, SS1, and 15B6) by competitive ELISA. The immobilized scFvs were challenged with biotinylated matMSLN preincubated with competing antibodies. The heatmap intensity reflects the degree of competitive binding, indicating epitope overlap. **(h)** Predicted epitope mapping of anti-MSLN scFvs on the basis of binning data. According to UniProt annotations, the extracellular domain of MSLN is divided into Region I (residues 296–398), Region II (399–494), and Region III (495–606). Three independent experiments were performed; the data are presented as the means ± standard deviations (s.d.).

On the basis of these selection criteria, seven antibody clones—CLMS10, CLMS20, CLMS32, CLMS36, CLMS76, CLMS88, and CLMS95—were selected for further analysis **(Fig. 2b)**. After the production of recombinant scFvs^46, 47^, their binding activities to both matMSLN and solMSLN were evaluated via ELISA **(Fig. 2c, d)**. Except for CLMS32 and CLMS36, all clones selectively bound to matMSLN (EC_50_ = 1.7 ± 0.1–151.1 ±[33.3 nM; **Supplementary Table 2**) over solMSLN. Notably, CLMS10 demonstrated high affinity for matMSLN (EC_50_ = 18.6 ± 1.1 nM), but minimal binding to solMSLN (EC_50_ > 1 μM). Surface plasmon resonance (SPR) confirmed that the affinity of CLMS10 for matMSLN was 8.22 ± 0.40 nM **(Supplementary Fig. 5)**. All scFvs, except CLMS20 and CLMS88, specifically bound to MSLN-positive K562 cells, with EC_50_ values ranging from a few nM to several hundred nM, while negligible binding to MSLN-negative K562 cells was detected **(Fig. 2e, f, and Supplementary Table 3)**. These results indicate that our anti-MSLN antibodies selectively recognize membrane-bound MSLN on the cell surface.

To further define the epitopes of the antibody candidates, we performed competitive ELISA using reference antibodies with well-characterized binding sites: SS1, which targets domain I (residues E296–L398)^28^, and 15B6, which binds the C-terminal region (residues Y594–L605).^25^ CLMS10, CLMS32, CLMS36, CLMS76, and CLMS95 competed for antigen binding, suggesting that they either share overlapping epitopes or exhibit steric hindrance that prevents simultaneous binding **(Fig. 2g)**. Notably, CLMS10 and CLMS76 exhibited weak binding to solMSLN, implying that their epitopes are localized near the C-terminus of domain III, which is known to be susceptible to proteolytic shedding **(Fig. 2d, h)**. Collectively, these findings suggest that the epitopes of CLMS10, CLMS32, CLMS36, CLMS76, and CLMS95 are clustered near the C-terminal region of domain III, supporting their therapeutic potential as shed-resistant antibodies **(Fig. 2h)**.

### Rapid and efficient live-cell screening of functional scFvs via a linRNA-based CAR-NK (linCAR-NK) cell platform

Standard screening methods often rely on immortalized cell lines (e.g., NK-92), which fail to recapitulate the complex biology of donor-derived cells. To ensure translational relevance, we implemented a “direct-to-primary” screening strategy. This streamlined approach enables rapid live functional screening directly in clinically relevant human primary NK cells, yielding high-fidelity phenotypic data and mitigating the risk of candidate failure during translational scale-up (**Figs. 1 and 3a**). Each scFv-containing CAR construct was synthesized as a linRNA via *in vitro* transcription (IVT). The quality of the synthesized linRNAs was validated via agarose gel electrophoresis and purity assessment (**Supplementary Fig. 6a**). Human primary NK (pNK) cells were electroporated with each linRNA, and CAR expression and cell viability were assessed in a time- and dose-dependent manner. For example, CAR-SS1-NK cells presented over 87% CAR expression at 1 µg linRNA per 10^6^ cells, with detectable expression persisting for up to 5 days, albeit at reduced levels **(Supplementary Fig. 6b, c)**. Following linRNA electroporation, flow cytometry confirmed CAR and GFP expression across all clones, with GFP coexpressed via a P2A linker. CAR expression levels varied among constructs and were quantified by staining with PE- or APC-conjugated streptavidin in combination with biotinylated MSLN extracellular domain proteins **(Supplementary Figs. 6d and 7)**. The cell viability was largely consistent across the clones (**Supplementary Fig. 6e)**.

**Fig. 3.**
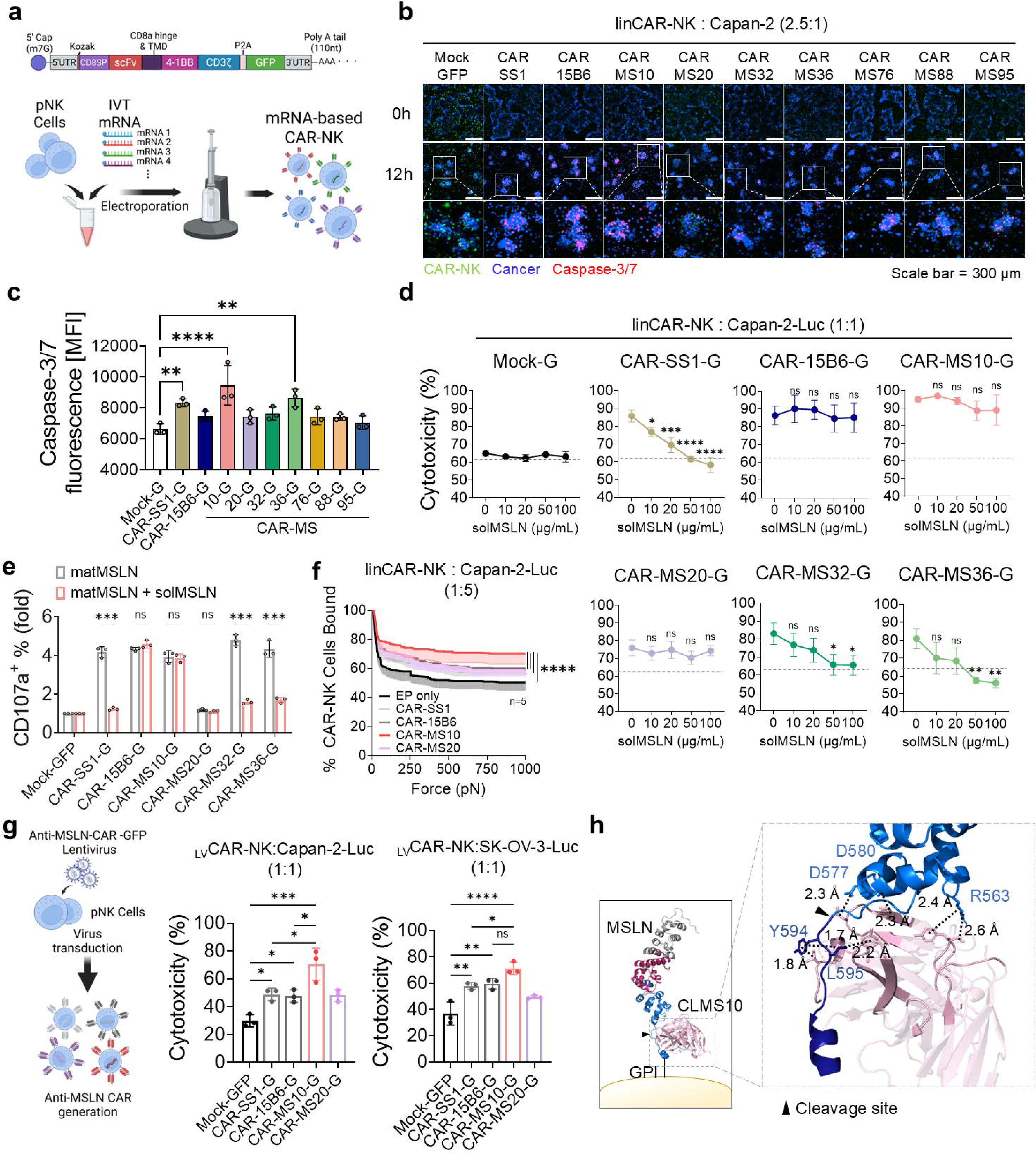
mRNA-based screening identifies CLMS10 as a potent shedding-resistant CAR construct in NK cells. **(a**) Generation of linear mRNA-based CAR (linCAR)-NK cells via electroporation (EP) of human primary NK cells. **(b, c)** Establishment of a rapid and cost-effective screening platform for evaluating CAR-NK cytotoxicity against Capan-2 cells. **(b)** Visualization of CAR-NK-mediated cancer cell killing. Capan-2 cells were stained with CellTrace Violet Cell (blue) and live-cell Caspase-3/7 dye (red), and cocultured with CAR-NK cells (green). **(c)** Quantification of apoptotic cancer cells based on Caspase-3/7 activation (*n* = 3). **(d)** Shedding inhibition assay for determining CAR-NK cytotoxicity in the presence of increasing concentrations of solMSLN (residues 296–592); dashed lines indicate the baseline cytotoxicity of allogeneic NK cells. **(e)** CD107a activation in CAR-NK cells stimulated with matMSLN-coated plates (20 µg/mL) in the presence or absence of solMSLN (50 µg/mL) was analyzed by flow cytometry (*n* = 3). **(f)** Cell–cell avidity between linCAR-NK and Capan-2 cells was quantified via acoustic force-based microfluidic microscopy (*n* = 5). **(g)** Luciferase-based cytotoxicity of lentivirus (LV)-generated CAR(_LV_CAR)-NK cells against Capan-2-Luc and SK-OV-3-Luc cells (*n* = 3). **(h)** Predicted docking model of CLMS10 scFv in complex with matMSLN using Schrödinger software. Residues with side chains within 2.5 Å of the binding interface are highlighted, revealing key antigen-antibody interactions. The soluble MSLN regions are indicated in marine blue, and the membrane-bound matMSLN regions are indicated in dark blue. The data are presented as the means ± s.d. Statistical analyses: one-way ANOVA in **(c)** and **(g)** and two-way ANOVA in **(f)**; two-tailed Student’s *t* tests in **(d)** and **(e)**; *\*p* < 0.05, *\*\*p* < 0.01, *\*\*\*p* < 0.001, *\*\*\*\*p* < 0.0001.

**Fig. 4.**
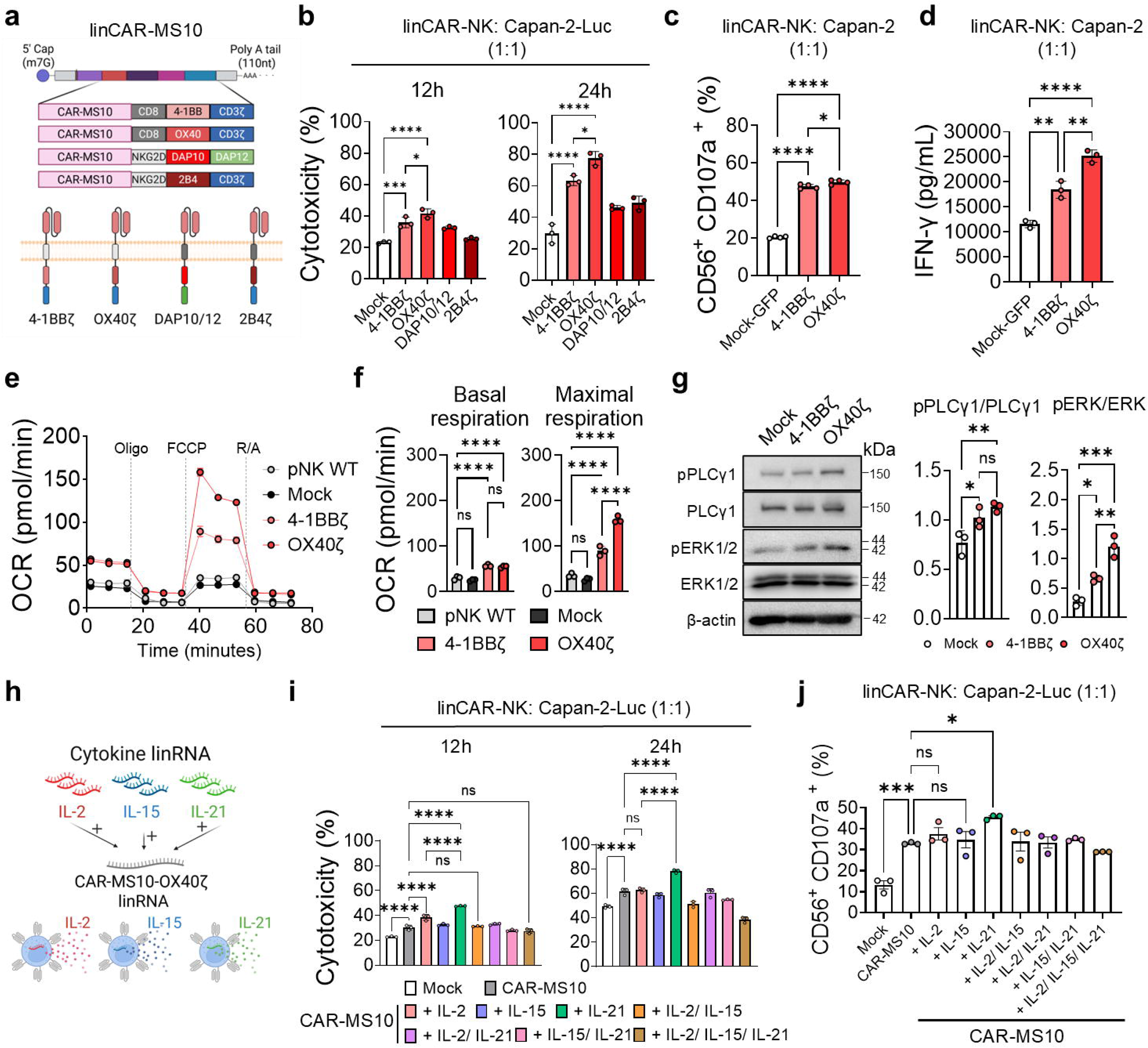
CAR-MS10 structural optimization and cytokine coexpression enhance CAR-NK cytotoxicity and functionality. **(a)** Four CAR constructs were designed to evaluate combinations of transmembrane (CD8, NKG2D), costimulatory (4-1BB, OX40, DAP10, and 2B4), and signaling (CD3ζ, DAP12) domains. **(b)** Cytotoxicity of linCAR-NK cells at 12 h and 24 h post coculture with Capan-2 cells determined via a luciferase-based assay (*n* = 3). **(c)** CD107a degranulation in CD56^+^ linCAR-NK cells after coculture with Capan-2 cells was analyzed by flow cytometry (*n* = 4). **(d)** IFN-γ secretion after 24 h of coculture (*n* = 3). **(e, f)** Metabolic activity [oxygen consumption rate (OCR)] after 2 h of stimulation on matMSLN-coated plates (20 µg/mL) (*n* = 3). **(g)** Intracellular signaling pathway activation by the lead construct (linCAR-MS10-OX40ζ), quantified by normalizing the levels of phosphorylated PLCγ and ERK to their respective total forms (*n* = 3). **(h–j)** Functional assessment of primary NK cells cotransfected with linCAR-MS10-OX40ζ and cytokine-encoding linRNAs (IL-2, IL-15, or IL-21). Luciferase-based cytotoxicity **(i)** and CD107a degranulation **(j)** were measured in NK cells cotransfected with single or combined cytokines (*n* = 3). The data are shown as the means ± s.d. One-way ANOVA was used in panels **(b–d)**, **(f)**, **(i)**, and **(j)**; *\*p* < 0.05, *\*\*p* < 0.01, *\*\*\*p* < 0.001, *\*\*\*\*p* < 0.0001.

We established a rapid live-cell screening platform to assess the cytotoxic function of linCAR-NK cells against an MSLN-overexpressing pancreatic cancer cell line (Capan-2). Real-time cancer cell death by linCAR-NK cells was monitored by tracking Caspase-3/7 activation during live coculture, with CAR-SS1-NK and CAR-15B6-NK cells used as controls **(Fig. 3b, c, and Supplementary Fig. 8a)**. Notably, CAR-MS10-NK cells caused a significant increase in cancer cell death, as evidenced by a 1.5-fold increase in Caspase-3/7-positive Capan-2 cells compared with mock-NK cells. A luciferase-based cytotoxicity assay revealed that CAR-MS10-NK cells achieved the strongest killing efficacy against both Capan-2-Luc and SK-OV-3-Luc cells, which express different MSLN levels **(Supplementary Fig. 8b, c)**. We assessed the sensitivity of each CAR-NK construct to solMSLN-mediated inhibition. In coculture with solMSLN, CAR-SS1-NK, CAR-MS32-NK, and CAR-MS36-NK cells exhibited a dose-dependent reduction in killing activity. In contrast, CAR-MS10-NK, CAR-15B6-NK, and CAR-MS20-NK cells maintained their cytotoxicity, indicating resistance to solMSLN-mediated inhibition **(Fig. 3d)**. The expression of CD107a, a marker of degranulation was unaffected by solMSLN treatment in CAR-MS10-NK, CAR-MS20-NK, and CAR-15B6-NK cells activated with matMSLN, further supporting their shedding resistance profiles **(Fig. 3e)**. We examined the extent of trogocytosis associated with impaired CAR function and NK-mediated fratricide in linCAR-NK cells.^48, 49^ Compared with CAR-SS1-NK cells, CAR-MS10-NK cells exhibited significantly reduced trogocytosis **(Supplementary Fig. 8d)**. Furthermore, CAR-MS10-NK cells presented the greatest degree of cell avidity among all the tested CAR constructs, suggesting the formation of robust immunological synapses **(Fig. 3f)**.^50^ To validate these findings in a more clinically relevant system, we introduced the scFvs into lentivirus-based CAR constructs, which were transduced into pNK cells. The results of functional assays revealed similar cytotoxicity patterns as those observed with linCAR-NK cells, with CAR-MS10-NK cells showing an approximately 22% higher killing rate against Capan-2 cells than CAR-SS1- and CAR-15B6-transduced cells **(Fig. 3g and Supplementary Fig. 9a)**. We evaluated whether this functional trend persisted across immune cell types such as linCAR-T cells. In cytotoxicity assays against Capan-2-Luc cells, linCAR-15B6-T cells showed comparable efficacy to linCAR-MS10-T and linCAR-SS1-T cells **(Supplementary Fig. 9b–f).** CAR-MS20-T cells showed limited activity, which is consistent with the NK cell results, suggesting that the functional impact of scFvs may vary depending on the immune effector cell type.^13, 14, 32, 51^

Finally, *in silico* docking analysis revealed a critical structure–function relationship. Unlike SS1, which binds to distal Region I, CLMS10 interacts with a conformational epitope that overlaps with or lies immediately adjacent to the proteolytic cleavage site (**Fig. 3h**). This unique binding geometry likely imposes steric hindrance that prevents access by sheddases or stabilizes the membrane-bound conformation of MSLN.

Consequently, CLMS10 selectively engages membrane-anchored MSLN while exhibiting drastically reduced affinity for the shed soluble form, providing a structural basis for its superior performance in shedding inhibition assays. Additionally, our linRNA-based screening platform offers an efficient functional approach for discovering and optimizing clinically relevant MSLN-targeting CARs.

### linRNA-driven screening for optimized CAR architecture and cytokine coexpression

We selected CAR-MS10 for further analysis. We employed a linRNA-based system to screen various combinations of transmembrane and costimulatory domains. Specifically, CD8 and NKG2D were chosen as the transmembrane domains (TMD_CD8_ and TMD_NKG2D_), and four CAR variants were designed: TMD_CD8_–4-1BB–CD3ζ (4-1BBζ), TMD_CD8_–OX40–CD3ζ (OX40ζ), TMD_NKG2D_–DAP10–DAP12 (DAP10/12), and TMD_NKG2D_–2B4–CD3ζ (2B4ζ) **(Fig. 4a).**

The cytotoxic potential of the CAR variants against Capan-2-Luc cells was evaluated via a luciferase-based killing assay. Among the variants, the CAR-MS10-OX40ζ construct exhibited the strongest cytotoxicity, followed by CAR-MS10-4-1BBζ **(Fig. 4b and Supplementary Fig. 10a)**. A detailed analysis of the functional activities across the 4-1BBζ and OX40ζ constructs revealed that, consistent with the cytotoxicity data, CAR-MS10-OX40ζ resulted in a significant increase in CD107a activation compared with the prototype CAR-MS10-4-1BBζ, accompanied by increased secretion of IFN-γ **(Fig. 4c, d)**. In addition to direct cytotoxicity, metabolic fitness is a key determinant of CAR-NK cell survival in the nutrient-deprived TME.^37^ The incorporation of the OX40 costimulatory domain significantly reprogrammed NK cell metabolism, enhancing both basal and maximal mitochondrial respiration upon antigen stimulation (**Fig. 4e, f**). This elevated spare respiratory capacity suggests that CAR-MS10-OX40ζ-NK cells are better equipped to sustain energy production under metabolic stress, contributing to prolonged antitumor activity.

To further investigate the underlying signaling pathways involved in OX40ζ-mediated CAR activation, CAR-MS10-OX40ζ-NK cells were stimulated with an MSLN recombinant protein and analyzed via western blotting **(Fig. 4g).** Compared with those in the 4-1BBζ group, the phosphorylation of phospholipase C gamma and extracellular signal-regulated kinase, key downstream molecules of CD3ζ signaling, was greater in the OX40ζ group.^52,53^ Collectively, these results identify OX40ζ as the optimal intracellular domain configuration for CAR-MS10, enhancing both effector function and signaling capacity.

We cotransfected cytokine-encoding linRNAs into CAR-MS10-OX40ζ cells to increase their cytotoxicity and functionality. Cytokines play pivotal roles in modulating NK cell activity, proliferation, and persistence.^54–56^ On the basis of their well-characterized functions, we selected interleukin-2 (IL-2), IL-15, and IL-21 to reinforce NK cell effector function and longevity. The synergistic effects of the selected cytokines were evaluated in CAR-MS10-NK cells, generated by coelectroporating CAR-MS10-OX40ζ linRNAs with linRNAs encoding each cytokine **(Fig. 4h and Supplementary Fig. 10b, c).** Among the various cytokine combinations, IL-21–CAR-MS10-OX40ζ-coexpressing NK cells presented the strongest antitumor activity against Capan-2-Luc cells, along with significantly increased CD107a expression **(Fig. 4i, j)**. Taken together, the results of the linRNA-based approach demonstrated that the IL-21–CAR-MS10-OX40ζ configuration endows NK cells with superior cytotoxicity and functional activity.

### CircRNA-based CAR-NK cells exhibited superior cytotoxicity with prolonged CAR expression

To address the transient nature of linRNA expression in mRNA-based cell therapies, we explored the use of circRNAs for prolonging CAR expression. Compared with linRNAs, circRNAs have previously been used to prolong transgene expression and offer improved structural stability and translational efficiency.^57–59^ We monitored CAR expression and cell viability over time in CAR-NK cells generated from either linRNAs or circRNAs **(Fig. 5a)**. Notably, circCAR-NK cells presented significantly prolonged and sustained CAR expression, as demonstrated by longitudinal flow-cytometry analyses of both the percentage of CAR[ cells and the CAR mean fluorescence intensity (MFI) across multiple timepoints (Days 1, 3, and 5) **(Fig. 5b, c)**. In contrast, there was no notable difference in viability between linRNA-and circRNA-transfected cells over the same time period **(Fig. 5d)**. We evaluated the persistence of cytotoxicity using serial killing assays, in which CAR-NK cells were rechallenged with fresh cancer cells for five consecutive rounds **(Fig. 5e)**. Consistent with prolonged CAR expression, circCAR-NK cells maintained superior activity, although a gradual decline was observed over time **(Fig. 5f)**. circCAR-NK cells exhibited robust IFN-γ secretion, supporting a direct association between prolonged CAR expression and sustained effector function **(Fig. 5g)**. circCAR-MS10-NK cells retained strong cytotoxic activity even under solMSLN-shedding conditions, indicating resistance to solMSLN-decoy effects **(Fig. 5h)**. In contrast, circCAR-SS1-NK cells showed limited cytotoxicity and a marked reduction in IFN-γ secretion in the presence of solMSLN, highlighting the importance of target epitope selection and construct design **(Fig. 5i)**. Importantly, circRNA-driven CAR-NK cells consistently demonstrated stronger cytotoxicity and NK-cell activation than linCAR- or _LV_CAR-NK cells did, further supporting the superior functional durability of the circRNA platform **(Supplementary Fig. 11).** Together, these findings underscore the superior performance of circCAR-NK cells over their linRNA counterparts, demonstrating enhanced CAR expression, prolonged cytotoxicity, and greater functional resilience under repeated tumor challenge and antigen-shedding stress.

**Fig. 5.**
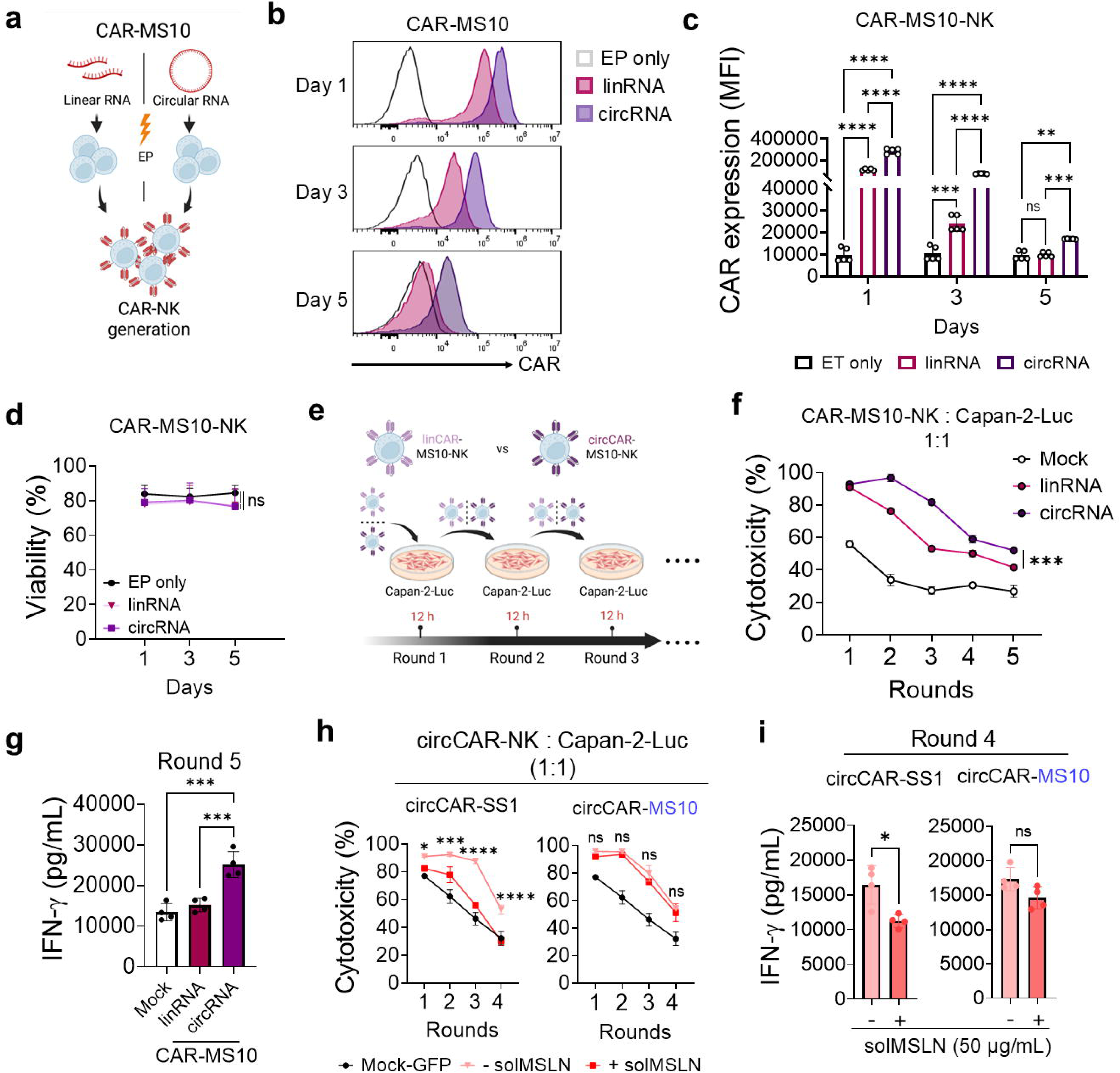
circCAR-MS10-CAR-NK cells exhibit prolonged CAR expression, sustained cytotoxicity, and resistance to antigen shedding. **(a)** Generation of circCAR-NK cells via EP. **(b–d)** CAR expression and viability of linCAR-and circCAR-NK cells over time. **(c)** CAR levels were measured and compared to those of the EP-negative control (*n* = 5). **(d)** NK viability was assessed by live/dead staining following EP (*n* = 3). **(e)** Serial killing assay of CAR-NK cells sequentially exposed to fresh Capan-2-Luc cancer cells at 12 h-intervals. Serial cytotoxicity **(f)** and IFN-γ secretion **(g)** were evaluated to determine the sustained antitumor activity of circCAR-NK cells (*n* = 3). **(h, i)** Impact of solMSLN (50 µg/mL) on CAR-NK cell function during repeated tumor challenges. Serial cancer killing **(h)** and IFN-γ secretion **(i)** were assessed following solMSLN treatment (50 µg/mL) at each round (*n* = 3). The data are presented as the means ± s.d. Statistical analyses: one-way ANOVA for **(c), (d), (f),** and **(g)**, and two-tailed Student’s t tests for **(h)** and **(i)**; *\*p* < 0.05, *\*\*p* < 0.01, *\*\*\*p* < 0.001, *\*\*\*\*p* < 0.0001.

### CircCAR-MS10-NK cells show potent antitumor activity in solMSLN-shedding models in vitro *and* in vivo

The pancreatic TME is highly immunosuppressive and is characterized by a dense extracellular matrix and abundant TGF-β signaling, which primarily originates from CAFs. This milieu impairs NK cell function and greatly challenges the efficacy of immunotherapies.^38–40^ Recent studies have demonstrated that anti-mesothelin antibodies not only target tumor cells but also disrupt antigen-presenting CAFs, reshaping the TME and enhancing immune responses.^60^ However, the therapeutic efficacy of MSLN-targeted CAR-T-therapies is undermined by the decoy effect of solMSLN-mediated antigen shedding, impeding CAR-Tcell activity.^25^ To address the suppression of CAR-Tcell function induced by solMSLN under physiologically relevant conditions, we established an *in vitro* cancer–CAF coculture model to mimic the TME. Interestingly, we found that the presence of human primary pulmonary fibroblasts or CAFs derived from patients with pancreatic cancer significantly accelerated the secretion of solMSLN compared with that of cancer cells alone **(Fig. 6a, b, and Supplementary Fig. 12a, b).**

**Fig. 6.**
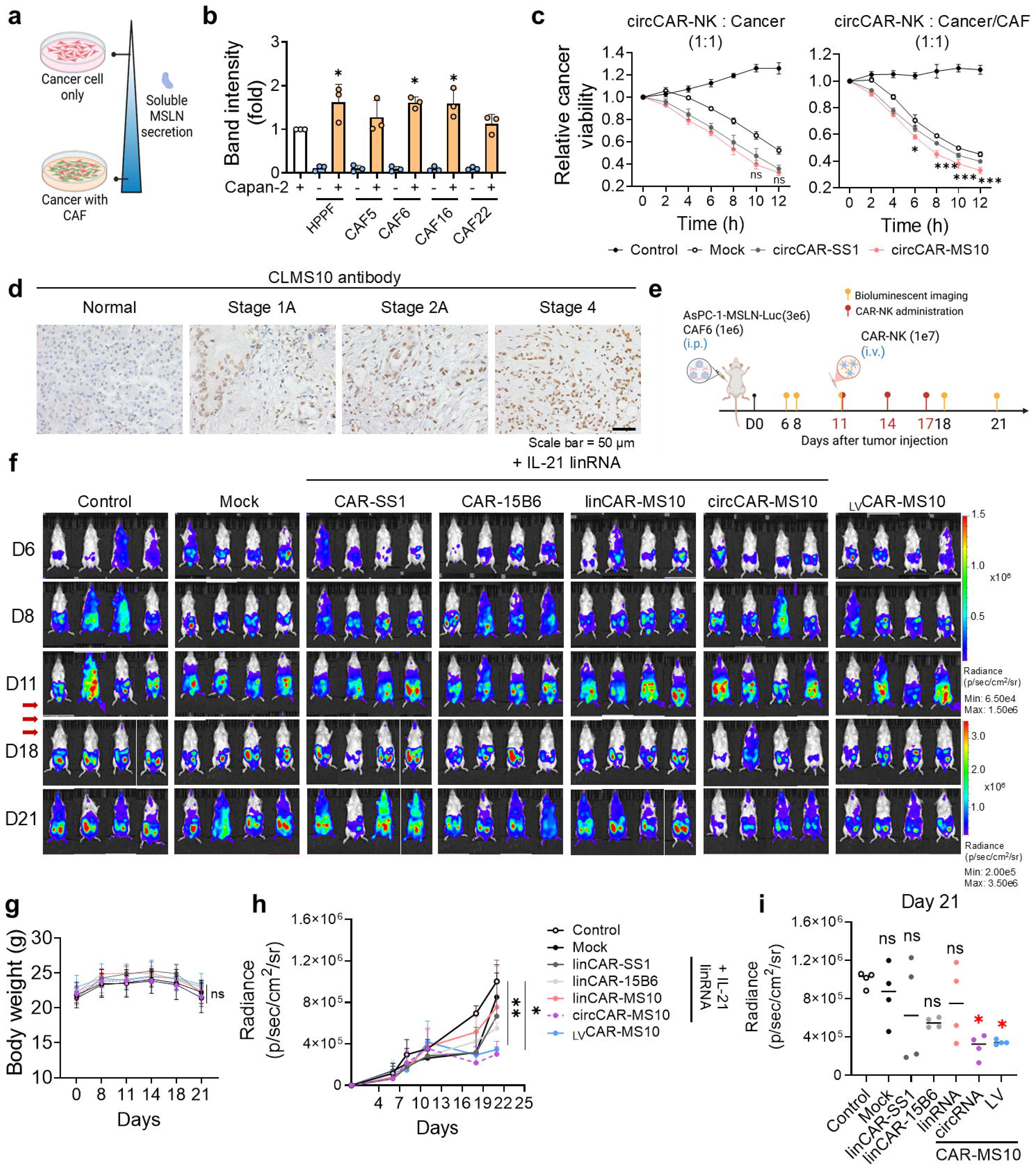
circCAR-MS10-NK cells maintain superior antitumor activity under CAF-induced shedding conditions *in vitro* and *in vivo* (a, **b)** Soluble MSLN (solMSLN) levels were elevated in Capan-2 cells cocultured with human pancreatic primary fibroblasts (HPPFs) or CAFs (CAF5, CAF6, CAF16, and CAF22), as shown by western blotting and quantified relative to those in cancer-only controls (*n* = 3). **(c)** *In vitro* real-time cytotoxicity of circCAR-NK cells cocultured with mCherry-expressing Capan-2 cancer cells, in the absence or presence of CAFs. CAFs were coincubated to mimic the tumor microenvironment characterized by shed MSLN secretion. Cancer cell survival was monitored by quantifying mCherry/RFP fluorescence, presented as relative cancer viability, which was defined as the mCherry signal normalized to the 0 h value (set to 1.0) (*n* = 3). Statistical comparisons were performed between the circCAR-SS1 and circCAR-MS10 groups at each indicated time point. **(d)** IHC of MSLN expression in human pancreatic tumor tissues using a CLMS10-derived antibody across stages 1A, 2A, and 4. **(e, f)** *In vivo* therapeutic effects of CAR-NK cells were evaluated in a metastatic mouse model with intraperitoneal AsPC-1-MSLN-Luc tumors containing CAF6 cells. **(f)** Representative IVIS images of mice treated with mock, linCAR-NK (SS1, 15B6, MS10), circCAR-MS10, or _LV_CAR-MS10 cells (*n* = 4 per group); all mRNA-CAR groups were coelectroporated with IL-21 mRNA. **(g)** Body weight monitoring during treatment. **(h, i)** Tumor burden quantified via IVIS imaging over time **(h)** and on day 21 **(i)** (*n* = 4). The data are presented as the means ± s.d. Statistical analyses: one-way ANOVA for **(b)** and **(h)** and two-way ANOVA for **(c), (g),** and **(i)**; \**p* < 0.05, \*\**p* < 0.01, \*\*\**p* < 0.001, \*\*\*\**p* < 0.0001.

Based on the observed increase in solMSLN secretion in the presence of CAFs, we further assessed the therapeutic efficacy of CAR-NK cells under CAF-mediated shedding inhibition via real-time live-cell imaging **(Fig. 6c, and Supplementary Fig. 12c)**. The CAF-induced increase in solMSLN levels significantly reduced cancer cell death in the circCAR-SS1-NK cell-treated group. In contrast, the cytotoxic efficacy of circCAR-MS10-NK cells was maintained despite MSLN shedding. Notably, strong MSLN expression was associated with advanced cancer progression, as evidenced by CLMS10-IgG staining in pancreatic tumor tissues, suggesting that MSLN is a therapeutically actionable target **(Fig. 6d)**. We established an *in vivo* intraperitoneal metastatic tumor model that mimics the TME^5, 25^, including CAF-mediated MSLN shedding, by coinjecting AsPC-1-MSLN-Luc tumor cells and CAFs into NOD-*Prkdc^em1Baek^Il2rg^em1Baek^*(NSG) mice and treating them with various CAR-NK platforms: linCAR-NK cells expressing CAR-SS1, CAR-15B6, or CAR-MS10; circCAR-MS10-NK cells; and _LV_CAR-MS10-NK cells **(Fig. 6e)**. To enhance NK cell function and persistence, IL-21 linRNA was coelectroporated into all the mRNA-based CAR-NK groups, but not into the lentivirus-based groups. Compared with the mock group, both the circRNA-based and lentivirus-based CAR-MS10-NK groups demonstrated significant tumor regression **(Fig. 6f-i)**. circCAR-MS10-NK cells showed anticancer activity comparable to that of _LV_CAR-MS10-NK cells. In contrast, compared with mock cells, linCAR-NK cells induced a modest reduction in tumor size, although the difference was not significant. These findings demonstrate that optimized circCAR-MS10-NK cells retain antitumor efficacy under TME-mimicking conditions and elevated solMSLN levels, supporting the potential applicability of the nonviral CAR-NK platform for treating solid tumors.

Given that IL-21 was codelivered to increase NK activity, we also assessed potential systemic IL-21-associated toxicity. Following three repeated infusions of CAR-NK cells, we observed no significant changes in body weight, gross liver and kidney morphology, or hepatic and renal function markers, indicating the absence of acute organ toxicity (**Supplementary Fig. 13**). These findings suggest that cytokine-augmented circCAR-NK cells do not elicit measurable systemic toxicity, further supporting their translational potential as safe and effective nonviral CAR-NK therapies for solid tumors. To further characterize the behavior of the different CAR-NK modalities *in vivo*, we next examined the distribution and persistence of the infused NK cells **(Supplementary Fig. 14)**. Quantification of CAR[ NK cells in the peripheral blood at 72 h confirmed successful engraftment in all the CAR-bearing groups, whereas the mock-NK group presented only background-level events. By day 7, distinct persistence patterns had emerged among the platforms. As expected from the transient nature of linear mRNAs, linCAR-NK cells were largely undetectable in the circulation and secondary lymphoid tissues, highlighting the short-lived *in vivo* expression driven by linRNAs. In contrast, IL-21–armored circCAR-NK cells demonstrated the most robust persistence among all groups, with clear detection in the spleen 7 days after intravenous injection, which aligns with the known survival- and proliferation-supporting functions of IL-21. Conversely, even _LV_CAR-NK cells showed limited persistence on day 7. To complement these *in vivo* persistence data and to directly assess the stability of CAR surface expression under repeated antigen exposure, we next evaluated CAR stability *in vitro* by repeatedly challenging CAR-NK cells with Capan-2-Luc tumor cells for up to four consecutive rounds **(Supplementary Fig. 15)**. Across all platforms, serial stimulation ultimately resulted in progressive CAR downregulation. Notably, the surface CAR MFI of linCAR-, _LV_CAR-, and nonarmored circCAR-NK cells rapidly decreased. However, IL-21–incorporated circCAR-NK cells presented a modest transient increase in CAR expression during the first stimulation round, followed by a gradual reduction over subsequent rounds. Overall, compared with nonarmored circCAR controls, IL-21 armoring substantially improved the maintenance of CAR surface expression across stimulation rounds within the CAR[ NK-cell population **(Supplementary Fig. 15)**. These findings indicate that transient IL-21 delivery helps to preserve CAR surface expression during repeated antigen encounters—an essential feature for sustaining NK-cell activity in solid-tumor contexts.

## Discussion

In this study, we identified and addressed a critical failure mode of solid tumor immunotherapies, specifically the suppression of CAR-NK activity by CAF-mediated antigen shedding.^61^ A major finding is that successful targeting of shed-prone antigens such as MSLN requires a dual mechanism involving (1) structural evasion of the decoy effect via membrane-proximal binders (CLMS10) and (2) sustained effector function via IL-21–armored, circRNA-driven expression. Our approach integrated rational antibody engineering, rapid and efficient mRNA-based functional screening, and circRNA technology to overcome the limitations of current solid tumor therapies. Our data revealed that in the presence of CAFs, which actively accelerate shedding, standard CARs (SS1) and transient expression systems fail. In contrast, our optimized circCAR-MS10-NK cells maintained robust cytotoxicity, providing a unique contribution to the field by defining the design principles required for next-generation solid tumor therapies.

A major obstacle in targeting MSLN is the shedding of the extracellular domain, which acts as a decoy receptor, sequestering CARs and diminishing cytotoxic potential.^26, 62^ To address this issue, we generated a panel of MSLN-specific antibodies and demonstrated the performance of CLMS10 for binding membrane-proximal epitopes while displaying minimal affinity for solMSLN (**Fig. 2**). Unlike standard screens, our two-step yeast surface display workflow ensures the identification of binders that retain structural integrity in the CAR format. A two-step Fab-to-scFv workflow further minimizes candidate dropout by first recovering correctly paired, stable VH/VL candidates in Fab format and then validating their compatibility after scFv conversion. This staged process ensured that only robust shed-resistant binders advanced into CAR screening. In contrast to SS1, the CLMS10-based CAR maintained strong cytotoxic activity even at high solMSLN concentrations, emphasizing the importance of epitope selection (**Figs. 2c–h and 3d–g**). Notably, these shed-resistant CARs demonstrated efficacy across both NK and T cells, indicating broad applicability across effector platforms. Mechanistically, the membrane-proximal binding of CLMS10 not only evades the “sink effect” but also significantly reduces trogocytosis compared with that of the distal-binding SS1 CAR (**Supplementary Fig. 8d**). Trogocytosis is known to drive CAR internalization, fratricide, and effector exhaustion.^34, 49^ The lower trogocytic activity of CLMS10 is likely attributable to its membrane-proximal, shed-resistant binding mode, which minimizes soluble antigen–mediated CAR clustering and contributes to more controlled synapse engagement. This restrained interaction may underlie the enhanced functional durability observed with CLMS10-based CAR-NK cells.

A key advantage of our workflow is the transition from cell-line-based screening to rapid mRNA-based functional assessment directly in primary immune cells. Conventional screens using immortalized lines (e.g., NK-92) often fail to predict translational efficacy due to divergent signaling and metabolic profiles. By employing electroporation-based delivery in primary NK cells, we rapidly phenotyped multiple CAR constructs, identifying the OX40ζ signaling domain and IL-21 codelivery as the optimal configuration. Notably, this combination enhanced cytotoxicity and significantly improved metabolic fitness, as evidenced by increased mitochondrial respiration (**Fig. 4**). As IL-21 is known to increase PI3K–Akt activation^55, 63^ and trigger the expression of T-BET, EOMES, and CEBPD^55, 64^, our findings suggest that the synergy between OX40 signaling and IL-21-mediated metabolic reprogramming enables NK cells to persist and function within the nutrient-deprived TME.

Another major advance is the application of circRNAs for CAR expression in NK cells. Compared with linRNAs, circRNAs exhibit prolonged transcript stability and sustained CAR expression, owing to their resistance to exonuclease degradation.^65, 66^ circCAR-NK cells remained functionally active during serial tumor challenges, with cytotoxicity maintained despite repeated antigen exposure **(Fig. 5)**. This highlights the unique advantage of circRNAs in bridging transient and durable CAR expression systems, combining the safety of RNA-based approaches with the persistence of integrated vectors. Despite its advantages, circRNA-based CAR expression remains transient, indicating that further RNA engineering strategies could enhance durability. The incorporation of stabilizing elements or translation-enhancing motifs around the IRES may prolong protein production from circular transcripts.^67, 68^ Additional approaches that increase RNA stability or translational persistence could likewise be applied to future CAR designs. Our data also show that IL-21 codelivery improves functional persistence, suggesting that cytokine armoring can complement RNA engineering by supporting NK-cell survival and activity. Together, these strategies represent promising avenues to strengthen the durability and therapeutic performance of nonviral CAR-NK platforms.

In terms of translational relevance, we tested our platform in a CAF-rich tumor model characterized by dense stroma, high solMSLN levels, and immunosuppressive factors. While CAFs compromised the cytotoxicity of circCAR-SS1-NK cells, circCAR-MS10-NK cells retained potent cytotoxicity (**Fig. 6a–c**). Human pancreatic tumor tissues presented high MSLN expression (**Fig. 6d**). In AsPC-1-MSLN-Luc/CAF xenografts, circCAR-MS10-NK cells achieved tumor regression, exhibiting activity comparable to that of lentiviral CAR-NK cells (**Fig. 6e–i**). These findings underscore the platform’s shed-resistance and persistence under clinically relevant TME conditions.

Our RNA engineering strategy supports the development of off-the-shelf, allogeneic CAR-NK therapies. Compared with T cells, NK cells are inherently less prone to GVHD and CRS, providing a potential universal donor base.^3, 4, 51^ Previous trials using HLA-mismatched cord blood-derived CAR-NK cells expressing IL-15 have demonstrated safe and durable responses in patients.^69^ By avoiding viral vectors, our mRNA and circRNA systems reduce manufacturing complexity, shorten production timelines, and enable rapid adaptation to new targets.^15^ The transient expression profile of RNA further permits repeat dosing and modular CAR redesign, offering unmatched flexibility. Furthermore, recent advances in circRNA synthesis, particularly self-splicing intron-based circularization, have improved yield and scalability, facilitating broader application of this approach in cell therapy manufacturing.^68,70,71^ While our shed-resistant circCAR-MS10-NK cells demonstrate strong potential in addressing antigen shedding and mitigating elements of the immunosuppressive TME, solid tumors such as pancreatic cancer remain highly refractory due to additional resistance mechanisms. These include antigen downregulation and the secretion of immunosuppressive factors such as TGF-β within the dense stromal matrix.^72–74^ Consequently, further studies are warranted to assess the long-term persistence, tumor trafficking, and safety of circRNA-engineered CAR-NK cells *in vivo*. Future therapeutic strategies may benefit from combining these cells with agents that target complementary immune escape pathways, such as immune checkpoint inhibitors or TGF-β blockade, to increase antitumor efficacy within the complex TME.

In conclusion, we propose a comprehensive framework for generating shed-resistant, long-lasting CAR-NK therapies with enhanced cell–cell avidity against solid tumors. By strategically integrating antibody engineering, rapid mRNA-based CAR discovery, and circRNA-mediated expression combined with IL-21, we address key limitations of CAR-based therapies for solid tumors. The IL-21–armored circCAR-MS10-NK system offers a potent, scalable, and clinically translatable solution for treating malignancies such as pancreatic cancer, effectively overcoming antigen shedding and immunosuppressive TMEs that have previously impeded therapeutic success.

## Materials and methods

### Isolation and *ex vivo* expansion of human pNK cells

PBMCs were purchased from Lonza (Walkersville, MD, USA). NK cells were cultured in medium from Miltenyi Biotec (Bergisch Gladbach, Germany). pNK cells were cultured and maintained in the presence of IL-2 (200 IU/mL; PeproTech, Rocky Hill, NJ, USA) and IL-15 (10 ng/mL; PeproTech), along with K562 feeder cells obtained from the American Type Culture Collection (Manassas, VA, USA). Following a 2-week expansion period, pNK cells were isolated via an NK cell isolation kit (Miltenyi Biotec) according to the manufacturer’s guidelines.

### Animals

NSG mice and BALB/c nude mice were purchased from JABIO (Gyeonggi-do, Korea). BALB/c mice were purchased from KOATECH (Gyeonggi-do, Korea). Male mice aged 6–8 weeks were randomly assigned to experimental groups for the study.

### Yeast strains and culture conditions

*S. cerevisiae* JAR200 (MATa, GAL1-AGA1::KanM×4ura3D45, ura3–52 trp1 leu2D1 his3D200 pep4::HIS3 prb1D1.6R can1) and YVH10 (MATα, PDI::ADHII-PDI-Leu2 ura3-52 trp1 leu2D1 his3D200 pep4::HIS3 prb1D1.6p can1 GAL) strains were used to construct and screen the yeast surface-display Fab library. Both strains were kindly provided by Prof. Dane Wittrup (Massachusetts Institute of Technology, Cambridge, MA, USA). The haploid yeast library was maintained in SDCAA (6.7 g/L yeast nitrogen base, 5 g/L casamino acids, 14.7 g/L sodium citrate, 4.29 g/L citric acid monohydrate, and 20 g/L dextrose in deionized water) supplemented with either uracil (0.0002 g/L) for JAR200 or tryptophan (0.0004 g/L) for YVH10, while the diploid Fab library was maintained in SDCAA without additional supplements. The Fab surface display library was induced in 2× SGCAA medium (13.4 g/L yeast nitrogen base, 10 g/L casamino acids, 5.4 g/L Na_2_HPO_4_, 8.56 g/L NaH_2_PO_4_·H_2_O, and 20 g/L galactose in deionized water) at a starting OD_600_ of 0.5, with incubation at 20 °C and 160 rpm for 48 h.

### Construction of the immune Fab library

The immune Fab library was constructed via a yeast surface display system to facilitate proper eukaryotic protein folding, overcoming the structural limitations often associated with prokaryotic phage display. A Fab-based format was employed to maximize combinatorial library diversity by mating, thereby achieving a library size and screening throughput significantly superior to those typically attainable in mammalian display systems. Following the final immunization, the mice were sacrificed, and mRNA was isolated from the spleen and bone marrow samples. The VH and VL genes were amplified from cDNA via a primer mixture **(Supplementary Table 1)**. Through overlapping PCR, each amplified VH and VL gene was fused to the CH1 and CL regions, respectively. The resulting VH-CH1 fragment (4 µg) was cotransformed with the linearized pYDS-H vector into *S. cerevisiae* strain JAR200, while the VL-CL fragment was cotransformed with the linearized pYDS-K vector into *S. cerevisiae* strain YVH10. Homologous recombination facilitated by electroporation was used for both transformations. The resulting haploid HC library in JAR200 (MATa) and the LC library in YVH10 (MATα) were cultured for 15 h at 30 °C and 160 rpm in 200 mL of SDCAA medium supplemented with uracil (for JAR200) or tryptophan (for YVH10), reaching an OD[[[ of ∼5 in the late exponential phase. For mating, equal volumes (OD[[[ = 100) of the HC and LC library cells were mixed thoroughly by vortexing and inversion. The mixture was plated onto YPD agar (pH 4.5) and incubated at 30 °C for 8 h. After incubation, the cells were washed twice with sterile distilled water, resuspended in SDCAA medium at an initial OD[[[ of 0.1, and then grown overnight at 30 °C with shaking at 160 rpm. To evaluate mating efficiency, serial dilutions of the mated cultures were plated on agar with either SDCAA or SDCAA supplemented with tryptophan. Mating efficiency was calculated as the ratio of colony counts on SDCAA to those on SDCAA with tryptophan.

### *In vitro* transcription (IVT) for linCAR

To construct the CAR plasmid, the newly generated scFv containing the second-generation CAR construct and GFP was cloned and inserted into the pmREX plasmid vector via the EZ-Fusion HT Cloning Kit (Cat. EZ015TL, Enzynomics, Daejeon, Republic of Korea) along with the SS1 (US Patent: US7081518) and 15B6 (US Patent: US7081518B1) scFv DNA fragments. The sequences of these scFvs were subsequently cloned and inserted into template DNA for mRNA synthesis. IVT of the mRNA was performed via the use of a linearized plasmid vector as the template and a MEGAscript T7 transcription kit (Cat. AMB13345; Invitrogen, Waltham, MA, USA). The synthesized mRNA was capped with CleanCap Reagent AG (Cat. N-7113, Trilink, San Diego, CA, USA). mRNA was precipitated via lithium chloride and purified, and the yield was measured via a NanoDrop 1000 UV visible spectrophotometer (Thermo Fisher Scientific, Waltham, MA, USA).

### Generation of CAR-NK cells via electroporation

The circRNA construct, designed as a second-generation CAR incorporating either the SS1 or CLMS10 scFv, was provided by GenScript (Nanjing, China). Expanded pNK cells were collected by centrifugation at 300 ×*g* for 5 min, and the culture medium was removed. The cells were washed twice with 10 mL of DPBS and resuspended in Resuspension Buffer T from a Neon Transfection System 100 μL Kit (4.0 × 10[ cells/mL; Invitrogen). CAR-NK cells were generated via IVT using 1 µg of mRNA per 10[ cells. The mRNA was mixed thoroughly with 100 µL of cells suspended in Resuspension Buffer T. Electroporation was performed via a Neon Transfection System (Invitrogen) to facilitate intracellular mRNA delivery. For circCAR-NK generation, 0.5 µg of circRNA per 10[ cells was used for electroporation. In the case of multiple transfections, the concentration of each mRNA was adjusted on the basis of its corresponding molecular weight (bp size) to achieve the appropriate molarity, and the final amount per 10[ cells was used for electroporation.

### *In vivo* efficacy of CAR-NK cells

All animal procedures were conducted in accordance with institutional guidelines and approved by the Institutional Animal Care and Use Committees of KIST (KIST–2024–079). A metastatic tumor model reflecting the TME with high solMSLN secretion was established via intraperitoneal injection of either 3 × 10[ AsPC-1-MSLN-Luc cells alone or a mixture of AsPC-1-MSLN-Luc and CAF6 cells at a 3:1 ratio. After 11 days, the mice were randomly divided into seven groups and treated under various conditions. Each tumor-bearing mouse received three intravenous administrations of 1 × 10[ linCAR-NK, circCAR-NK, or _LV_CAR-NK cells. Tumor progression was monitored via *in vivo* luciferase imaging on an IVIS Spectrum imaging system (Revvity, Waltham, MA, USA) following intravenous injection of a luciferin substrate solution (Promega, Madison, WI, USA) under isoflurane anesthesia. The tumor burden was quantified by measuring the average radiance (photons/s/cm²/sr).

Further methodological details are provided in the Supplementary Materials.

## Statistical analysis

All the data are expressed as the means ± s.d. Statistical analyses were performed via GraphPad Prism 10. Two-tailed Student’s *t*-test or two-way ANOVA was performed, depending on the experimental design, with Tukey’s post hoc analysis. *p* < 0.05 was used to identify statistically significant differences.

## Ethics statement

All animal experiments were performed in accordance with guidelines approved by the Institutional Animal Care and Use Committee of Seoul National University (protocol no. SNU-221108-1-1) and KIST (protocol no. KIST–2024–079).

## Acknowledgments

This work was supported by the National Research Foundation (NRF) of Korea, funded by the Korean Government (grant nos. RS-2024-00463774, RS-2024-00400042, RS-2024-00509114, RS-2023-NR076986, RS-2025-02216943, and RS-2023-00278980). Additional support was provided by the KIST Institutional Program (grant no. 2E33761) and the Seoul National University Hospital (SNUH) Research Fund (grant no. 03-2024-3030).

## Author contributions

M.J. and C.L. conceived and designed the study; J.C., J.H., S.Yee., H.L., D.L., S.Yoon., S.L., Y.J., S.S., and S.K. performed the experiments and data analysis; J.C. and J.H. wrote the original draft of the manuscript; M.J. and C.L. revised the manuscript; and M.J. and C.L. supervised the project and validated the final version. All authors have read and approved the final manuscript.

## Data availability statement

The original raw microscopy images underlying the figures, including uncropped images and ROI information, have now been deposited in a public repository (DOI: 10.6084/m9.figshare.31109158).

