## Supplementary info for "Rapid CAR screening and circRNA-driven CAR-NK cells for persistent shed-resistant immunotherapy"

**This PDF file includes:**

Parts of materials and methods

Supplementary Figures. S1 to S15

Supplementary Tables S1 to S3

Uncropped western blots

**Parts of Materials and Methods**

**Cell lines and culture**

MSLN-positive AsPC-1, parental AsPC-1, MSLN-positive K562, and parental K562 cells were cultured in RPMI 1640 medium (Welgene, Gyeongsan, Republic of Korea) supplemented with 10% fetal bovine serum (FBS; Gibco, Waltham, MA, USA) and 1× penicillin‒streptomycin (Thermo Fisher Scientific, Waltham, MA, USA). All the cells were maintained at 37 °C in a humidified atmosphere containing 5% CO_2_. The pancreatic cancer Capan-2 and ovarian cancer SK-OV-3 cell lines were obtained from the Korean Cell Line Bank (Seoul, Republic of Korea). Capan-2 cells were cultured in RPMI 1640 medium (Welgene) supplemented with 10% FBS (Welgene) and 1% antibiotic–antimycotic solution (Welgene). SK-OV-3 cells were cultured in McCoy's 5A medium (Thermo Fisher Scientific) supplemented with 10% FBS (Thermo Fisher Scientific) and 1% antibiotic-antimycotic (AA) solution (Welgene). HEK293T cells were obtained from American Type Culture Collection (ATCC; Manassas, VA, USA) and were maintained in Dulbecco’s modified Eagle’s Medium (DMEM; Welgene) supplemented with 10% FBS (Welgene), 1% AA solution (Welgene) and 10 mM HEPES. Primary pancreatic CAFs and HPPFs were obtained from Asan Medical Center (Seoul, Republic of Korea) and ScienCell Research Laboratories (Carlsbad, CA, USA), respectively. Both CAFs and HPPFs were cultured in RPMI 1640 medium (Welgene) supplemented with 10% FBS (Welgene), 1% AA solution (Welgene), and 10 ng/mL fibroblast growth factor (FGF; PeproTech, Rocky Hill, NJ, USA). All the cultures were maintained at 37 °C in a humidified incubator with 5% CO_2_.

**Flow cytometry**

The cells were harvested, washed, and incubated with antibodies and detection reagents for the analysis of cellular proteins via flow cytometry. The following antibodies and reagents were used: mouse anti-FLAG antibody (MA1-91878, 1:200, Invitrogen, Waltham, MA, USA), FITC-anti-mouse IgG (F0257, 1:200, Sigma-Aldrich, MO, USA), SA-PE (S866, 1:200, Invitrogen), SA-APC (SA1005, 1:200 Invitrogen), anti-HIS-FITC (ab1206, 1:400, Abcam, Cambridge, UK), eBioscience Fixable Viability Dye eFluor 780 (65-0865-14, 1:200, Thermo Fisher Scientific), APC-conjugated anti-CD107a antibody (328620, 1:100, BioLegend, San Diego, CA, USA), anti-mesothelin antibody (ab196235, 1:1000, Abcam), CD56-APC (17-0566-42, 1:100, Invitrogen), BD Pharmingen PE Streptavidin (554061, 1:1000, BD Pharmingen, CA, USA), BD Pharmingen APC Streptavidin (554067, 1:1000, BD Pharmingen) and BD Pharmingen FITC Mouse Anti-Human CD45 (555482, 1:5, BD Pharmingen). The detailed labeling conditions, including the antibody concentrations and incubation protocols, are described in the corresponding Methods section.

For primary NK cells, the collected cells were washed three times with staining buffer (DPBS supplemented with 2% FBS) and stained with eBioscience Fixable Viability Dye eFluor 780 (Thermo Fisher Scientific) for 20 min at 4 °C. After staining, the cells were washed twice with staining buffer and then incubated with cell surface antibodies for 20 min at 4 °C. Following incubation, the cells were washed again and resuspended in 300 μL of staining buffer prior to analysis. Flow cytometric analysis was performed via a CytoFLEX flow cytometer (Beckman Coulter Life Sciences, Munich, Germany), and the data were analyzed via FlowJo software (BD Biosciences, San Jose, CA, USA).

**Preparation of recombinant proteins**

To express matMSLN (residues 296–606) and solMSLN (residues 296–592), both MSLN gene sequences (synthesized by Integrated DNA Technologies, Coralville, IA, USA) were cloned and inserted into the pcDNA3.4 mammalian expression vector via Gibson assembly. Protein expression was performed in Expi293 cells (Thermo Fisher Scientific), and His-tagged MSLN proteins were purified by affinity chromatography using Ni-NTA agarose resin (QIAGEN, Germantown, MD, USA). Following elution and buffer exchange, protein purity was confirmed by SDS-PAGE. Recombinant MSLN proteins were biotinylated via an EZ-Link Sulfo-NHS Biotinylation Kit (Thermo Fisher Scientific).

**Mouse immunization**

BALB/c mice (6 weeks old) were immunized biweekly via the intraperitoneal injection of 50 μg of MSLN-His protein mixed with aluminum hydroxide adjuvant [1:1 (v/v); Thermo Fisher Scientific]. Blood samples were collected weekly to monitor the immune response. Sera were separated by centrifugation and used to measure anti-MSLN antibody levels. For the ELISA, 100 ng of matMSLN was coated onto the bottom of each well of a 96-well MaxiSorp microplate (Thermo Fisher Scientific) and incubated overnight at 4 °C. After being blocked with phosphate-buffered saline (PBS, pH 7.4) containing 3% bovine serum albumin (BSA; Sigma‒Aldrich, St. Louis, MO, USA) for 1 h at room temperature, the plates were washed three times with PBST (PBS, pH 7.4 containing 0.05% Tween 20). Mouse sera were serially diluted 10-fold and incubated on the experimental plates for 1 h at room temperature. After washing, horseradish peroxidase (HRP)-conjugated goat anti-mouse IgG (H+L) secondary antibody (31430, 1:8,000 Invitrogen) was added, and the mixture was incubated for 1 h. After three additional washes, the plates were developed with 3,3′,5,5′-tetramethylbenzidine (TMB) substrate. The reaction was stopped by the addition of 2 M H_2_SO_4_, and the absorbance was measured at 450 nm via an Infinite 200 PRO NanoQuant microplate reader (Tecan Trading AG, Männedorf, Switzerland).

**Screening of the immune Fab library**

To display the Fab fragments on the yeast surface, the Fab library was inoculated at a starting OD_600_ of 0.5 in 2× SGCAA medium and incubated at 20 °C with shaking at 160 rpm for 48 h. The expression of the surface-displayed Fabs was assessed by labeling the FLAG tag at the C-terminus of the LC with a mouse anti-FLAG antibody (MA1-91878, 1:200, Invitrogen). The yeast was coincubated with a biotinylated antigen for 1 h at room temperature. After incubation, the cells were washed with PBSA (pH 7.2, containing 0.1% BSA). Secondary labeling was performed using FITC-conjugated anti-mouse IgG (F0257, 1:200, Sigma‒Aldrich) and SA-PE (S866, 1:200, Invitrogen) or SA-APC (SA1005, 1:200, Invitrogen) at 4 °C for 15 min in the dark. After a final wash with PBSA, the cells were analyzed and sorted by flow cytometry using an SH800S cell sorter (Sony Biotechnology Inc., San Jose, CA, USA). For negative selection, MSLN-negative AsPC-1 cells were seeded in 6-well plates (6 × 10^5^ cells/well) and incubated at 37 °C and 5% CO_2_ for 2 d. Following incubation, the cells were fixed with 10% neutral-buffered formalin. To remove nonspecific binders, 5 × 10^7^ yeast cells (OD_600_ = 5) were added to wells containing MSLN-negative AsPC-1 cells and incubated at 4 °C with gentle shaking for 2 h. The supernatant was collected, and the wells were washed by adding 1 mL of ice-cold PBSA. For positive selection, the cells collected from the negative selection were incubated with MSLN-positive AsPC-1 cells. After incubation, the supernatant was discarded, and the cells were washed twice by adding 1 mL of ice-cold PBSA buffer to each well. SDCAA medium was then added to each well and the cells were scraped and cultured overnight at 30 °C with shaking at 160 rpm.

**Preparation of recombinant anti-MSLN scFv proteins**

The selected binders were subjected to a Fab-to-scFv conversion workflow to mitigate developability risks, ensuring that only candidates retaining high affinity and stability in the scFv format were selected. Accordingly, the antibody genes were subcloned in-frame into the pMopac12 vector via Gibson Assembly (New England Biolabs, Ipswich, MA, USA). The resulting scFv-pMopac12 plasmid was transformed into *Escherichia coli* BL21 (DE3) cells, which were cultured in 500 mL of terrific broth supplemented with chloramphenicol at 37 °C and 200 rpm until an OD_600nm_ of 0.6–0.8 was reached. Protein expression was induced with 0.5 mM IPTG, and the cultures were incubated for an additional 16 h at 25 °C with shaking at 160 rpm. The cells were harvested via centrifugation at 12,000 ×*g* for 1 h and resuspended in 32 mL of Tris-sucrose solution (pH 7.5; 100 mM Tris and 0.75 M sucrose). Lysozyme (20 mg/mL) was added to the suspension before incubation at 4 °C with shaking. The cells were treated with 1 mM EDTA (pH 8.0) and incubated once more. The periplasmic fraction was collected via centrifugation and filtration. His-tagged recombinant scFv proteins were purified via affinity chromatography.

**ELISA of anti-MSLN scFvs**

The binding activities of the anti-MSLN scFvs were assessed via ELISA. Each well of a 96-well MaxiSorp microplate was coated with 100 ng of scFvs and incubated overnight at 4 °C. After blocking with 3% PBSA for 1 h at room temperature, biotinylated MSLN-His antigens were serially diluted and added to the wells. The plates were incubated at room temperature for 1 h, washed with PBST, treated with streptavidin-HRP (1:40,000, Abcam, Cambridge, MA, USA), and incubated for 1 h at room temperature. After washing, TMB substrate was added, the reaction was stopped with 2 M H_2_SO_4_, and the absorbance at 450 nm was measured via an Infinite 200 PRO NanoQuant microplate reader (Tecan Trading AG).

**Competitive ELISA of anti-MSLN scFvs**

Each well of a 96-well MaxiSorp microplate was coated with 100 ng of scFv and incubated overnight at 4 °C. After blocking with 3% PBSA for 1 h at room temperature, biotinylated MSLN-His antigens (100 nM) or preincubated biotinylated MSLN-His (100 nM) and anti-MSLN scFv (400 nM) complexes were added to the wells. The plates were incubated at room temperature for 1 h, washed with PBST, treated with streptavidin-HRP (1:40,000, Abcam), and incubated for 1 h at room temperature. Following the wash step, TMB substrate was added to initiate the reaction. The reaction was subsequently stopped with 2 M H_2_SO_4_, and the absorbance was measured at 450 nm using an Infinite 200 PRO NanoQuant microplate reader (Tecan Trading AG). The outcomes were calculated by normalizing the antigen-only signal to 100% and expressing the measured signals as a percentage of this value.

**Cell binding assay of anti-MSLN scFvs**

MSLN^+^ K562 and MSLN^-^ K562 (5 × 10^4^) cells were resuspended in PBSF, plated in a 96-well round-bottom plate, and centrifuged at 300 ×*g* for 3 min, after which the cell pellets were washed with fresh PBSF. Recombinant anti-MSLN scFvs were added at various concentrations and incubated at 4 °C for 1 h. After incubation, the cell pellets were washed with fresh PBSF and then incubated with anti-HIS-FITC (ab1206, 1:400, Abcam) at 4 °C for 15 min. The cells were washed with PBSF and analyzed with a BD FACSCanto II flow cytometer (BD Biosciences) to quantify scFv binding.

***In silico* structure analysis**

The three-dimensional structures of the scFvs were determined via AlphaFold3 (Google DeepMind) ^1^. The model with the highest predicted local distance difference test and TM score was selected for further analysis. Structural comparisons were conducted via PyMOL, and protein–protein docking of MSLN to each scFv was performed via the antibody docking mode in Maestro 13.6 (Schrödinger Suite; Schrödinger, New York, NY, USA) while masking non-CDR regions. Among the generated poses, the one with the strongest binding affinity (under default parameters) was selected for subsequent structural evaluation.

**Surface plasmon resonance (SPR)**

The binding kinetics of CLMS10 IgG were assessed via a BIAcore T200 instrument. The mature form of MSLN was immobilized onto CM5 sensor chips via amine coupling, following the manufacturer’s guidelines (Cytiva, Marlborough, MA, USA). HBS EP buffer (BR100669, Cytiva) was used as the running buffer. CLMS10 IgG, prepared in a series of dilutions, was injected at 30 μL/min for 120 s, followed by a 5‑min dissociation phase. The chip was then regenerated by sequential 30‑s injections of 5 mM NaOH and 0.5 M arginine (pH 8.0). Equilibrium dissociation constants (K_D_) were derived by fitting the data to a 1:1 Langmuir model via BIAevaluation 3.2 software (Cytiva). Each K_D_ value represents the mean from three independent experiments (*n* = 3).

**Immunohistochemistry (IHC)**

We evaluated the binding capacity of CLMS10 IgG for human pancreatic cancer tissue via PDAC tumor tissue microarrays (PA482a Tissue Array, Derwood, MD, USA). The tissue sections were heated at 60 °C in a drying oven for 30 min, followed by deparaffinization and rehydration. Antigen retrieval was performed via Epitope Retrieval Solution (Bond Polymer Refine Detection Kit, DS9800; Leica Biosystems, Nußloch, Germany). After blocking, the sections were incubated for 30 min with CLMS10 IgG (1.22 mg/mL, 1:30 dilution) and treated with 3,3′-diaminobenzidine (DAB) substrate (Bond Polymer Refine Detection Kit, DS9800, Leica Biosystems) for signal detection. The slides were mounted and imaged using an Olympus BX53 microscope (Olympus Corporation, Tokyo, Japan). All the IHC staining procedures were conducted following the standard protocols of the Pathology Specimen Production Center at Seoul National University.

**linCAR-T-cell production**

PBMCs were thawed and seeded in 24-well TC-treated plates (#CLS3524, Corning, NY, USA) precoated with 10 µg/mL anti-human CD3ε antibody (#16-0037-85, eBioscience) in DPBS for 3 h at 37 °C. The cells were cultured in AIM-V medium (#12055-091, Gibco) supplemented with IL-2 (200 IU/mL; PeproTech), GlutaMAX (#35050061, 100X, Gibco), and 2 µg/mL anti-human CD28 antibody (#16-0289-85, eBioscience). After 24 h of incubation at 37 °C in 5% CO₂, additional AIM-V medium was added depending on the cell condition. T-cell culture was conducted with RPMI 1640 medium (Welgene) supplemented with IL-2 (200 IU/mL). Primary T cells were expanded and harvested via centrifugation at 500 ×*g* for 5 min, after which the culture medium was removed. The cells were washed twice with 10 mL of DPBS and resuspended in Resuspension Buffer T (Invitrogen). The final cell concentration was adjusted to 4.0 × 10⁷ cells/mL. The IVT of the mRNA encoding the CAR construct was conducted at 1 µg per 10⁶ cells after thorough mixing in a 100 µL cell suspension. Electroporation was performed via the Neon Transfection System (Invitrogen) under the following conditions: 1600 V, 10 ms, 3 pulses.

**Live cell imaging-based Caspase-3/7 assay**
To evaluate linCAR-NK cell candidates, Capan-2 cells were stained with a CellTrace Violet Cell Proliferation Kit (Invitrogen) according to the manufacturer’s instructions. The IVT of linCAR-NK cells involves GFP linked to the CAR construct via a P2A sequence. For the assay, Capan-2 cells were cocultured with CAR-NK cells at a 2.5:1 effector : target (E:T) ratio in 100 µL of coculture medium supplemented with IL-2 (200 IU/mL). To measure apoptosis in real-time, CellEvent Caspase-3/7 detection reagents RED (100X; Invitrogen) was added to the culture wells. The plates were placed into a BioTek Lionheart FX live cell imaging system (Agilent Technologies, Santa Clara, CA, USA) and incubated for 12 h to monitor Caspase-3/7 activation as an indicator of apoptosis. Images were captured at the indicated times via an automated live-cell imager.

**Luminescence-based cancer-killing assay**
We assessed the cytotoxic effects of NK cells on NanoLuc luciferase (Luc)-overexpressing cancer cells.^2,3^ Capan-2-Luc or SK-OV-3-Luc cancer cells were cocultured with pNK cells at the indicated E:T ratios. Following coculture, cancer cell viability was determined in terms of luciferase activity measured via a Nano-Glo luciferase assay system (Promega, Madison, WI, USA) according to the manufacturer’s instructions.

**Shedding inhibition assay**

To assess the impact of solMSLN on CAR-NK cell activity, a luminescence-based cancer-killing assay was performed. Cancer cells were cocultured with CAR-NK cells at the indicated E:T ratios for 12 h in IL-2 (200 IU/mL)-supplemented medium with or without solMSLN at concentrations up to 100 µg/mL.

For the CD107a degranulation assay, matMSLN protein (20 µg/mL)-coated 96-well plates were seeded with CAR-NK cells (1.0 × 10⁵ cells/well) and stimulated for 4 h with or without solMSLN (50 µg/mL). During stimulation, CD107a surface expression was assessed via an APC-conjugated anti-CD107a antibody (1:100; 328620; BioLegend) and BD GolgiStop containing Monensin (554724; BD Biosciences). After 4 h, the cells were stained with a viability dye and analyzed via flow cytometry via FlowJo software (BD Biosciences).

**Cell avidity assay**

Monolayer Capan-2 cells (5.0 × 10⁷/mL) were seeded into precoated z-Movi chips (LUMICKS, Amsterdam, Netherlands) at 20 μL per well to achieve 60–80% confluency. linCAR-NK cells were harvested and labeled with Vybrant DiD Cell-Labeling Solution (5 μL/test; V22887, Invitrogen) following the manufacturer’s protocol. After 15 min of staining, labeled linCAR-NK cells (1.0 × 10⁷/mL) were loaded into the chips (20 μL per test) and coincubated with the adherent monolayer cells for 10 min. An acoustic force ramp (0–1000 pN over 2.5 min) was then applied via the z-Movi Cell Avidity Analyzer (LUMICKS) to quantify the interaction strength between the effector and target cells. Avidity was assessed via image acquisition via both brightfield and fluorescence channels and analyzed via Ocean software (LUMICKS). Monolayer integrity and the detachment of tracked cells were recorded and quantified as measures of cell–cell avidity.

**Trogocytosis assay**

To assess trogocytosis in CAR-NK cells, Capan-2 cells were seeded in a 48-well plate (1.0 × 10^5^ cells/well) and incubated for 12 h; CAR-NK cells were added at a 1:1 ratio and cocultured for 4 h. Cell–cell interactions were stimulated via brief centrifugation (200 ×*g*, 30 s, 4 °C). Following coculture, the cells were harvested in cold DPBS and stained with an anti-mesothelin antibody (ab196235; 1:1,000; Abcam) and a CD56-APC antibody (.17-0566-42; 1:100; Invitrogen) to identify NK cells that had undergone trogocytosis. The degree of trogocytosis was determined as the percentage of MSLN^+^ cells within the CD56^+^ NK cell population via CytoFLEX (Beckman Coulter Life Sciences).

**Metabolic activity assay**

To promote cell adherence, a Seahorse XFe96/XF Pro Cell Culture Microplate (Agilent Technologies) precoated with poly-D-lysine (A-003-E, Sigma-Aldrich) was used. linCAR-NK cells were seeded at 2.5 × 10^5^ cells per well in Seahorse XF DMEM (pH 7.4; 103575-100; Agilent Technologies) supplemented with 10 mM glucose, 1 mM pyruvate, and 2 mM glutamine. The oxygen consumption rate was measured via a Seahorse XF Pro analyzer (Agilent Technologies). To assess mitochondrial function and metabolic activity, oligomycin (1.5 μM), FCCP (1 μM), rotenone (0.5 μM), and antimycin A (0.5 μM) were sequentially added at the indicated time points.

**Detection of secreted solMSLN in coculture media**

Capan-2 cancer cells and CAFs were seeded in 100 mm plates, either separately or together (2 × 10⁶ cells). After 24 h in serum-containing media, the cells were washed and incubated with serum-free RPMI 1640 supplemented with 1% AA and 10 ng/mL FGF for 48 h. The conditioned media were collected and concentrated 40-fold via Amicon Ultra centrifugal filters (10 kDa MWCO; Merck Millipore, Darmstadt, Germany) at 2,600 ×*g* at 4 °C for 45 min. The concentrated media were used for western blot analysis.

**Western blot analysis**

The cells were lysed via RIPA buffer (Thermo Fisher Scientific) containing a protease and/or phosphatase inhibitor cocktail. Proteins from lysates or conditioned media were separated by SDS‒PAGE and transferred onto polyvinylidene difluoride membranes blocked with 5% skim milk in 0.2% Tween 20/TBS for 2 h at room temperature and incubated overnight at 4 °C with an anti-mesothelin antibody (ab196235, 1:1000, Abcam). After washing, the membranes were incubated with HRP-conjugated goat anti-rabbit IgG (32460, 1:10,000, Invitrogen) and visualized via SuperSignal West Pico PLUS substrate (34580, Thermo Fisher Scientific) with a ChemiDoc imaging system (Bio-Rad Laboratories, Hercules, CA, USA).

**IFN-γ detection**

To evaluate cytokine secretion *in vitro*, various NK cell subsets were cocultured with Capan-2 cells. After 48 h of incubation, the culture supernatants were harvested, and the IFN-γ levels were quantified via a human IFN-γ ELISA kit (DIF50C, R&D Systems, Minneapolis, MN, USA) according to the manufacturer’s instructions.

**Lentivirus production and titration**

HEK293T cells were cotransfected with a CAR-expressing plasmid, the psPAX2 packaging plasmid, and a Baboon envelope plasmid at a 3:2:1 ratio via polyethylenimine transfection reagent (Polysciences Inc., Warrington, PA, USA). After 24 and 48 h, the supernatant containing the lentivirus was collected and concentrated via the Lenti-X Concentrator (Takara, Kusatsu, Shiga Japan). The viral titer was then determined via the Lenti-X p24 Rapid Titer Kit (Takara) following the manufacturer’s protocol.^3^

**linRNA and circCAR-NK-mediated serial killing assay**

To assess serial killing capacity, linCAR and circCAR-NK cells were cocultured with Capan-2-Luc cells (4 × 10^4^ Capan-2-Luc cells/well) at a 1:1 ET ratio. After 12 h of coculture, the CAR-NK cells were collected via centrifugation, resuspended in fresh IL-2-supplemented media (200 IU/mL), and transferred to newly seeded Capan-2-Luc cells. This process was repeated multiple times to evaluate the duration of cytotoxicity. The cytotoxicity at each round was quantified via a luminescence-based assay. For solMSLN shedding inhibition, the assay was conducted in media containing 50 µg/mL recombinant solMSLN protein.

**CAR downregulation assay under serial target stimulation**

CAR-NK cells were cocultured with Capan-2-Luc cells at a 1:1 E:T ratio in 48-well plates. For each round of stimulation, 1 × 10^5^ Capan-2-Luc cells were seeded in individual wells, and CAR-NK cells were added and incubated for 12 h. After each coculture period, the NK cells were recovered via gentle centrifugation, washed once with RPMI 1640, resuspended in fresh IL-2–supplemented media (200 IU/mL), and transferred to newly prepared target-cell wells at the same E:T ratio to initiate the next round. This procedure was repeated for up to four sequential rounds (Rounds 1–4). To assess CAR stability, NK cells from each round were collected from separate wells, ensuring parallel sampling across rounds. An aliquot of NK cells from each round was stained for flow cytometric analysis of surface CAR expression.

***In vivo* toxicity assay**

For histological evaluation, liver and kidney tissues were collected on day 6, fixed in 10% neutral-buffered formalin, embedded in paraffin, and sectioned at 4–5 μm. Deparaffinized and rehydrated sections were stained with hematoxylin and eosin using standard procedures. Stained tissues were examined under light microscopy to assess potential organ injury, including inflammatory infiltrates, necrosis, or structural abnormalities.

For hematological and biochemical toxicity measurements, blood was obtained from the abdominal vein (*n* = 4 per group). Approximately 500 μL of whole blood was centrifuged at 500 ×*g* for 15 min at 4 °C, and the plasma was carefully transferred to collection tubes. Serum aspartate aminotransferase, alanine aminotransferase, and blood urea nitrogen levels were quantified using an AU480 Chemistry Analyzer (Beckman Coulter Life Sciences) with Pureauto S series reagents (Sekisui Medical, Japan) following the manufacturer’s guidelines. These parameters were used to evaluate acute hepatic and renal toxicity following repeated administration of CAR-NK cells.

***In vivo* persistence analysis of CAR-NK cells**

Male BALB/c nude mice (6 weeks old) were used for the CAR-NK persistence study. Each mouse received a single intravenous injection of 1 × 10⁷ NK cells from each indicated platform. Peripheral blood was collected 72 h after NK-cell infusion via orbital bleeding, and CAR expression was analyzed by flow cytometry. At 7 days post-injection, the mice were euthanized, and both the peripheral blood and the spleen were harvested for NK-cell persistence analysis. Single-cell suspensions from spleens were prepared by mechanical dissociation and filtration through a 40-µm strainer (SPL Life Sciences, Gyeonggi-do, Korea, 93040). Red blood cells from both blood and spleen samples were lysed using RBC lysis buffer (BioLegend, 420301) prior to staining. To quantify CAR-NK persistence, flow-cytometry gating was performed to identify viable CD45⁺ NK cells, followed by measurement of the CAR⁺ population within this compartment.

**Supplementary Figures**

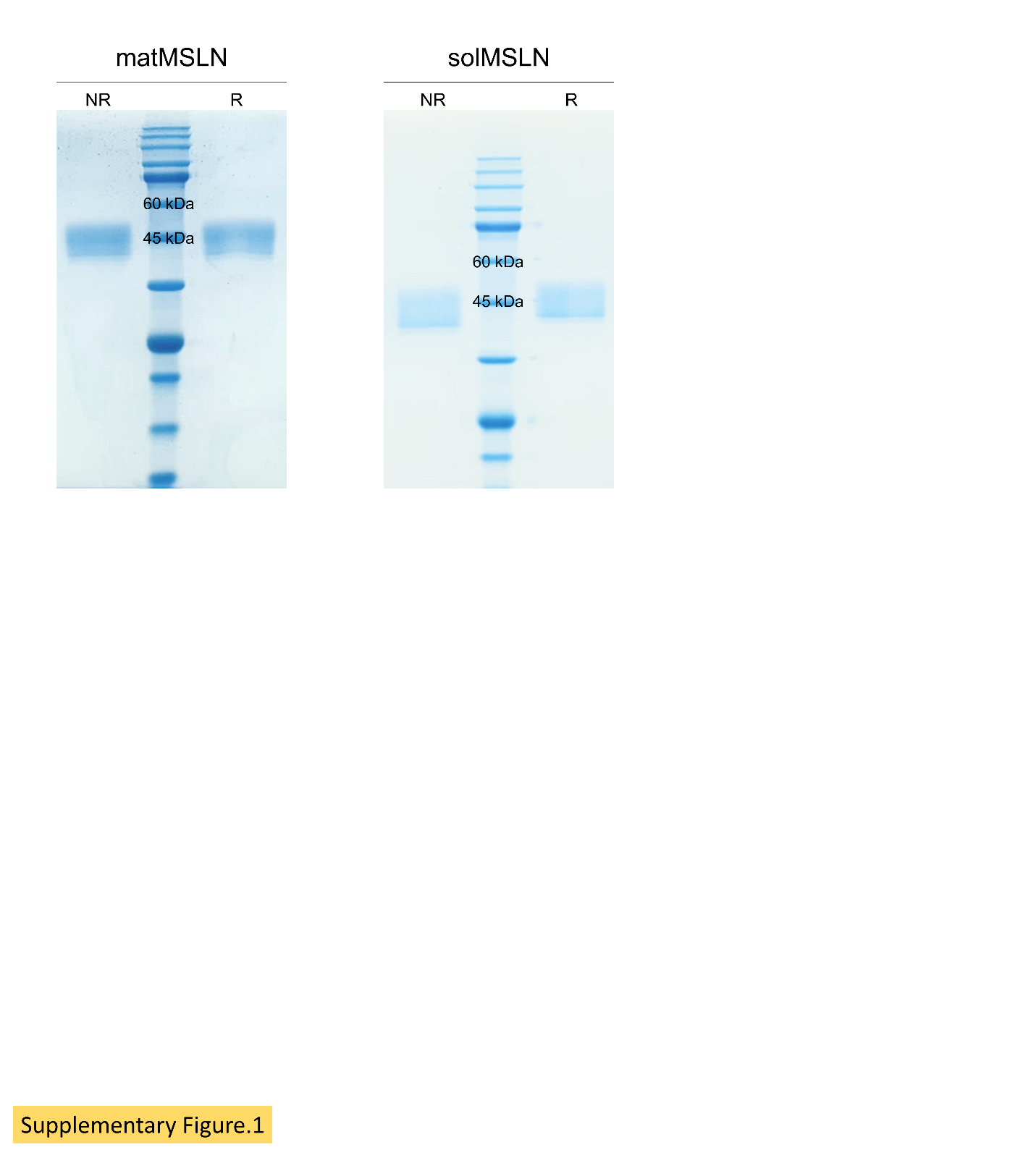

**Supplementary Figure 1. Preparation of recombinant MSLN proteins**

Recombinant mature mesothelin (matMSLN) and soluble mesothelin (solMSLN) were expressed in Expi293 cells and purified via Ni-NTA affinity chromatography. SDS-PAGE analysis was performed under non-reducing (NR) and reducing (R) conditions, with matMSLN in the left panel and solMSLN in the right panel.

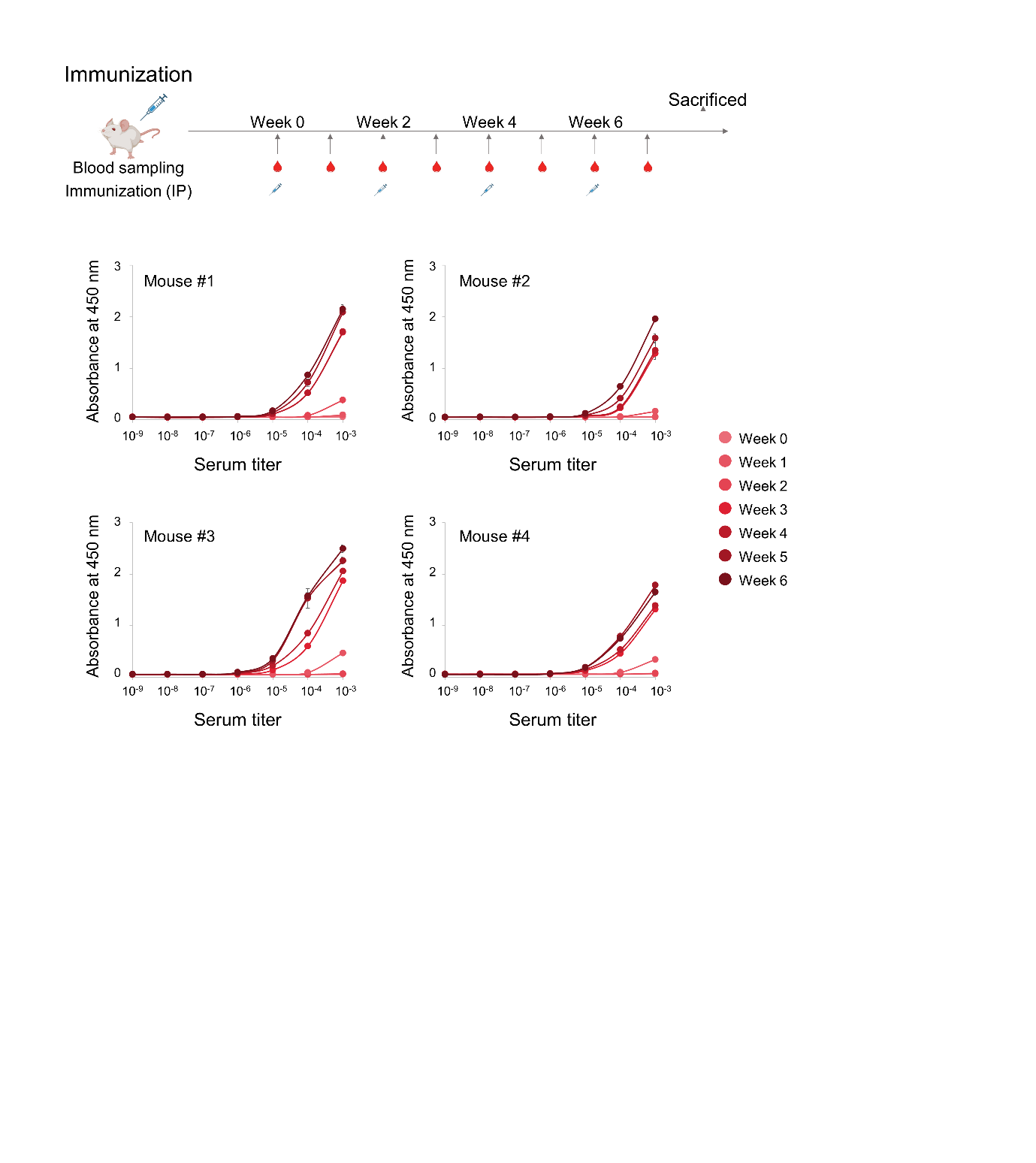

**Supplementary Figure 2. Strategy for mouse immunization**

Mice were intraperitoneally immunized with recombinant MSLN, and serum samples were collected weekly throughout the immunization period. MSLN-specific antibody titers were quantified using an ELISA. Briefly, immunoplates were coated with recombinant MSLN, followed by incubation with serially diluted mouse sera. Bound antibodies were detected with HRP-conjugated goat anti-mouse IgG. All experiments were performed in triplicate; data are presented as mean ± s.d.

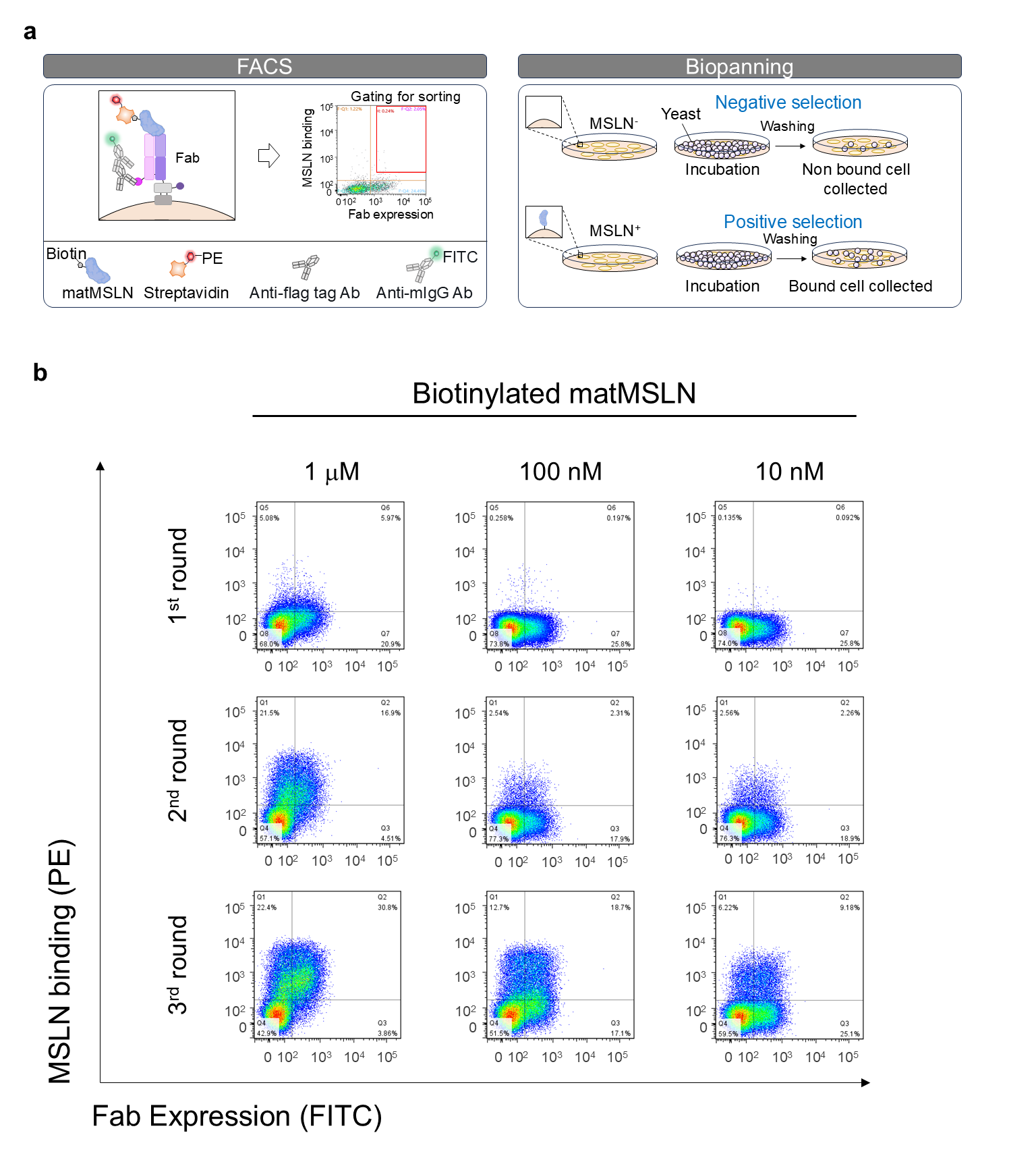

**Supplementary Figure 3. Screening of immune Fab library using yeast surface display**

**(a)** Schematic illustration of FACS-based sorting strategy. The immune Fab library, displayed on the surface of yeast, was screened for clones binding both recombinant MSLN proteins and MSLN-expressing cells. To remove non-specific binders, yeast cells displaying the Fab library were first incubated with MSLN-negative AsPC-1 cells (negative selection), then with MSLN-positive AsPC-1 cells to enrich MSLN-specific binders (positive selection). **(b)** Flow cytometry analysis of Fab surface expression (x-axis) and MSLN binding (y-axis) during sequential rounds of library screening.

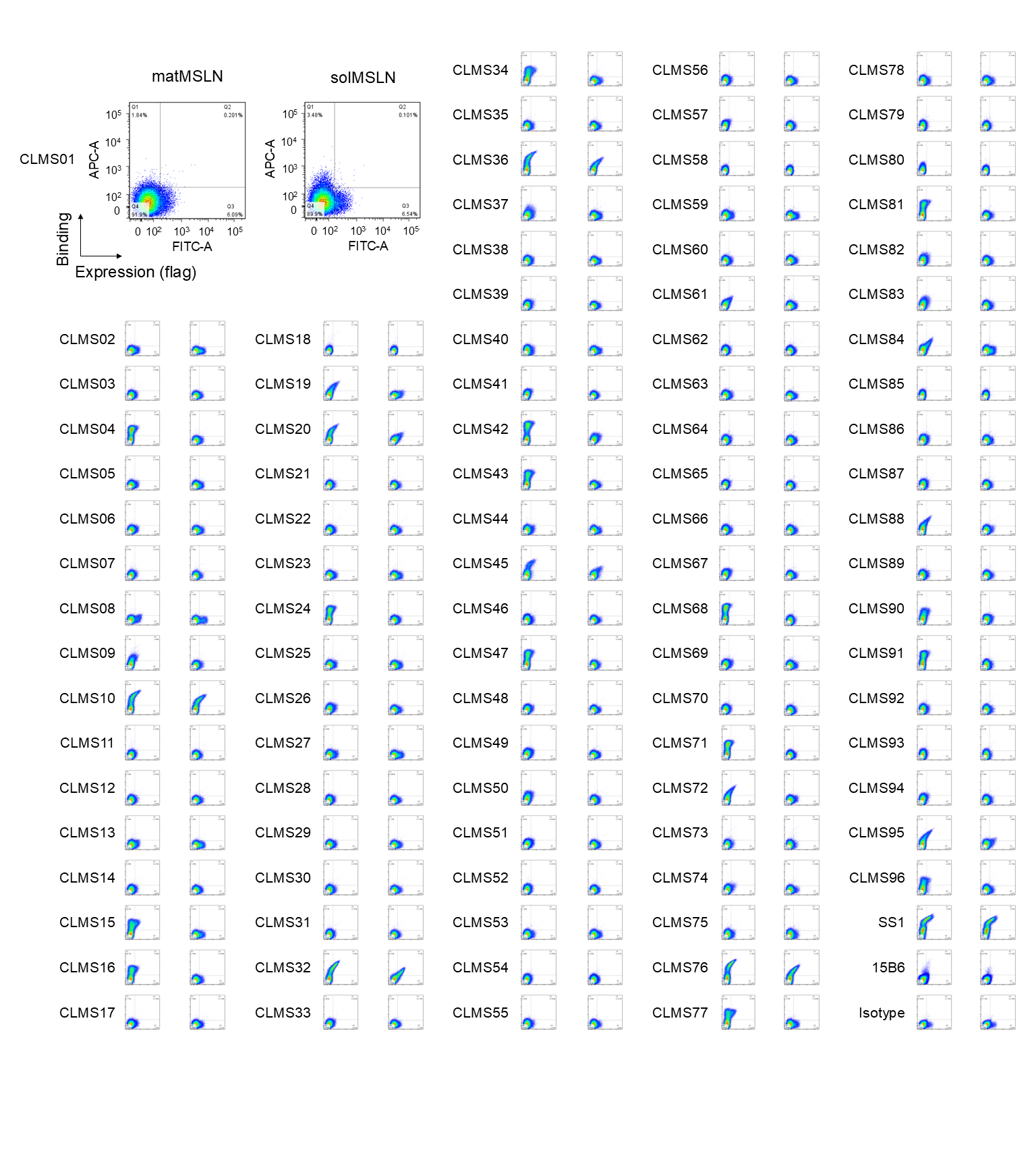

**Supplementary Figure 4. Flow cytometric analysis of individual anti-MSLN Fab clones** Diploid yeast cells expressing individual anti-MSLN Fab clones were incubated with 10 nM biotinylated matMSLN or solMSLN. Antigen binding was detected using streptavidin- allophycocyanin, and Fab surface expression was assessed using an anti-FLAG antibody followed by goat anti-mouse IgG-FITC staining. SS1 and 15B6 were included as positive controls, and an isotype antibody served as a negative control.

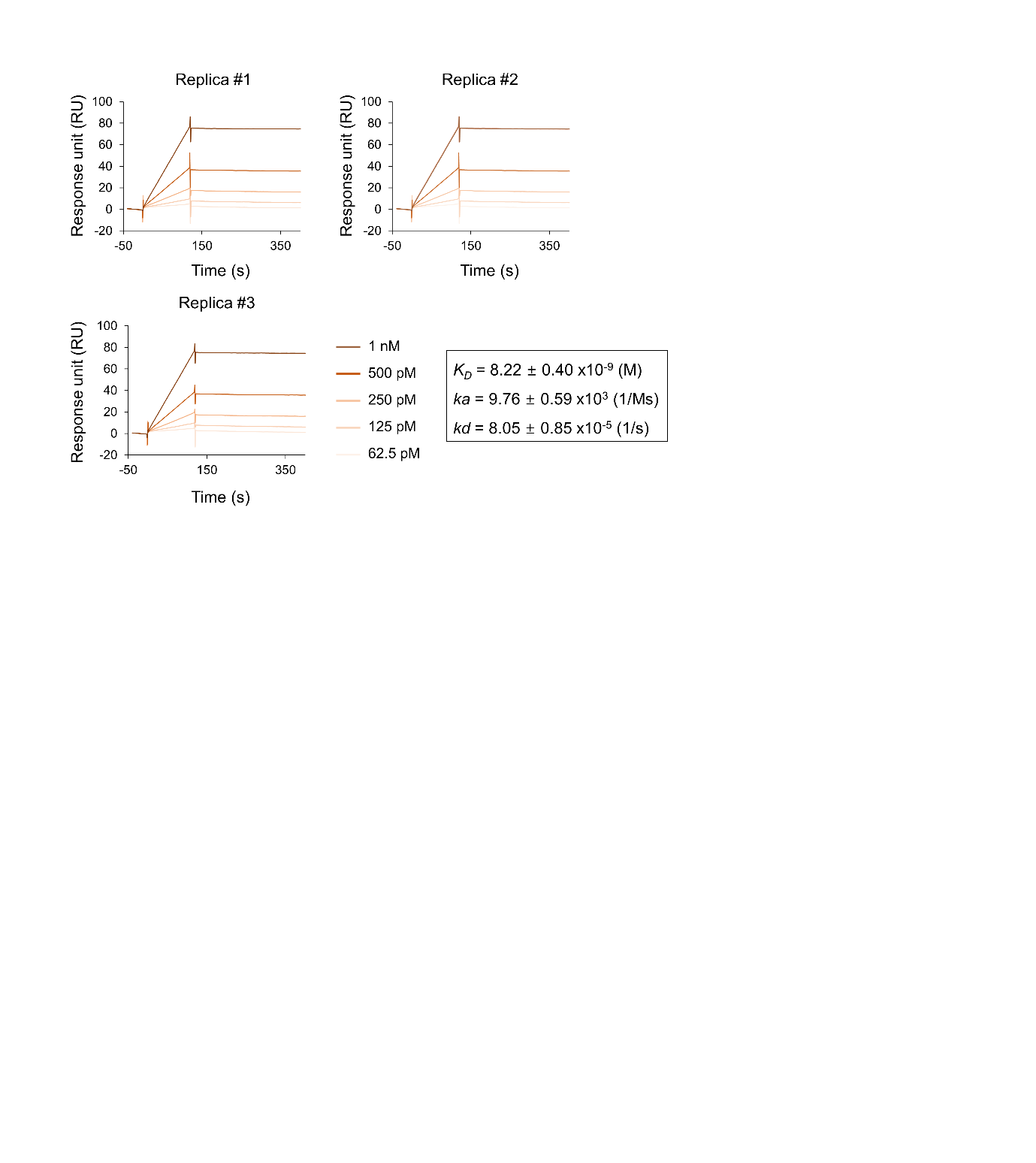

**Supplementary Figure 5. SPR analysis of CLMS10 IgG**

Binding affinity of CLMS10 IgG to recombinant MSLN was evaluated by SPR. Recombinant MSLN was immobilized on CM5 sensor chips, and serial dilutions of CLMS10 IgG were injected across the chip surface. The sensorgrams display real-time binding kinetics. Data are presented as mean ± s.d of three independent experiments.

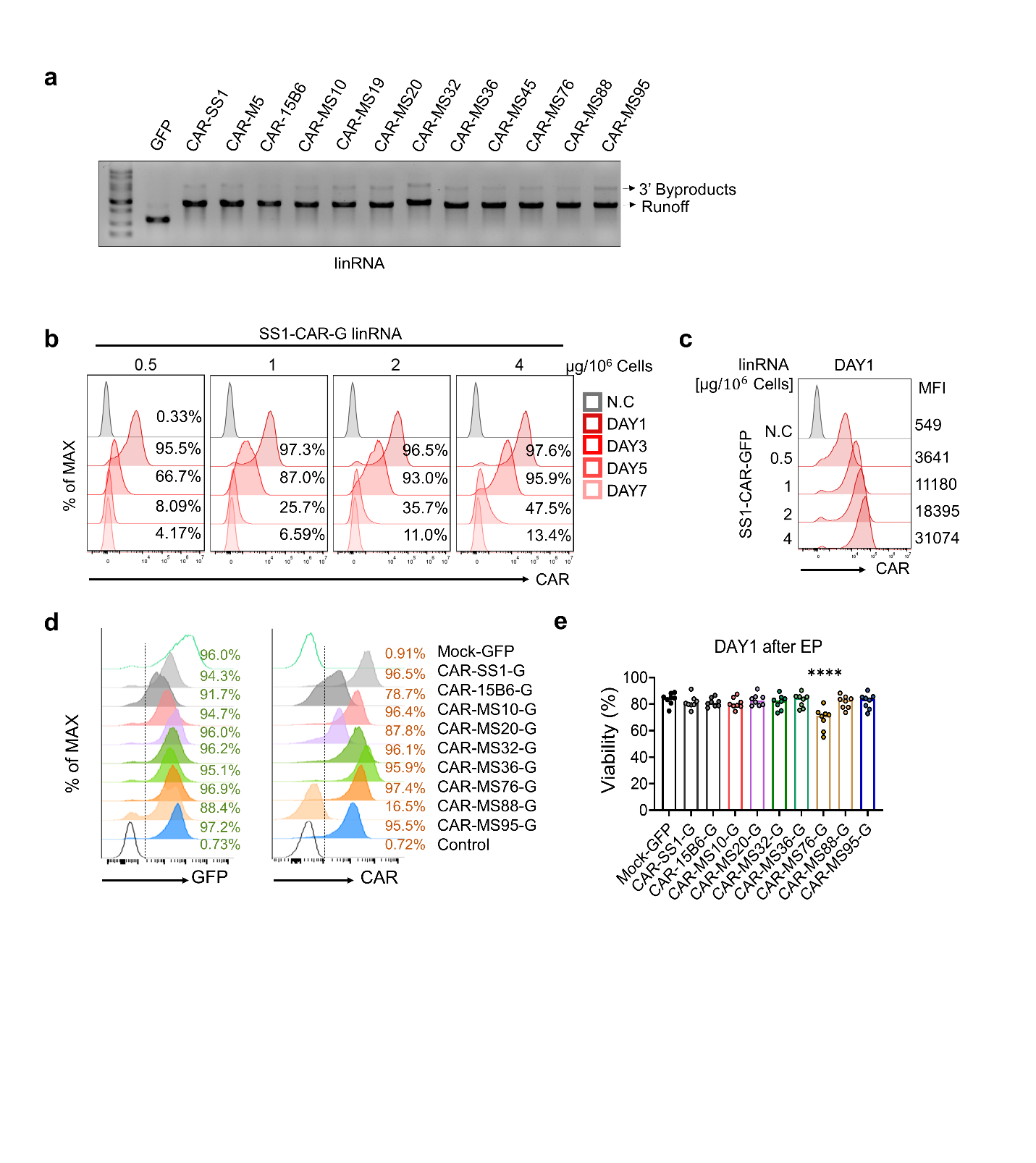

**Supplementary Figure 6.** **Characterization of linCAR-NK cells in terms of expression efficiency and viability**

**(a)** Agarose gel electrophoresis confirming the integrity and purity of *in vitro* transcribed linRNAs encoding various CAR constructs. **(b)** Time-course analysis of CAR expression in primary NK cells electroporated with 0.5–4 µg linRNA per 10⁶ cells, assessed by flow cytometry over 7 days. **(c)** Dose-dependent CAR expression, measured as MFI of CAR-SS1 at 24 h post-EP across linRNA concentrations (0.5–4 µg/10⁶ cells). **(d)** Flow cytometry of CAR and GFP expression across different scFv constructs using biotinylated matMSLN protein and PE-streptavidin, measured 24 h post-electroporation. **(e)** Cell viability of CAR-NK cells at day 1 post-EP, showing consistent viability across CAR variants. Data were analyzed using one-way ANOVA in **(e)**; Data are presented as mean ± s.d.; **p* < 0.05, ***p* < 0.01, ****p* < 0.001, and *****p* < 0.0001.

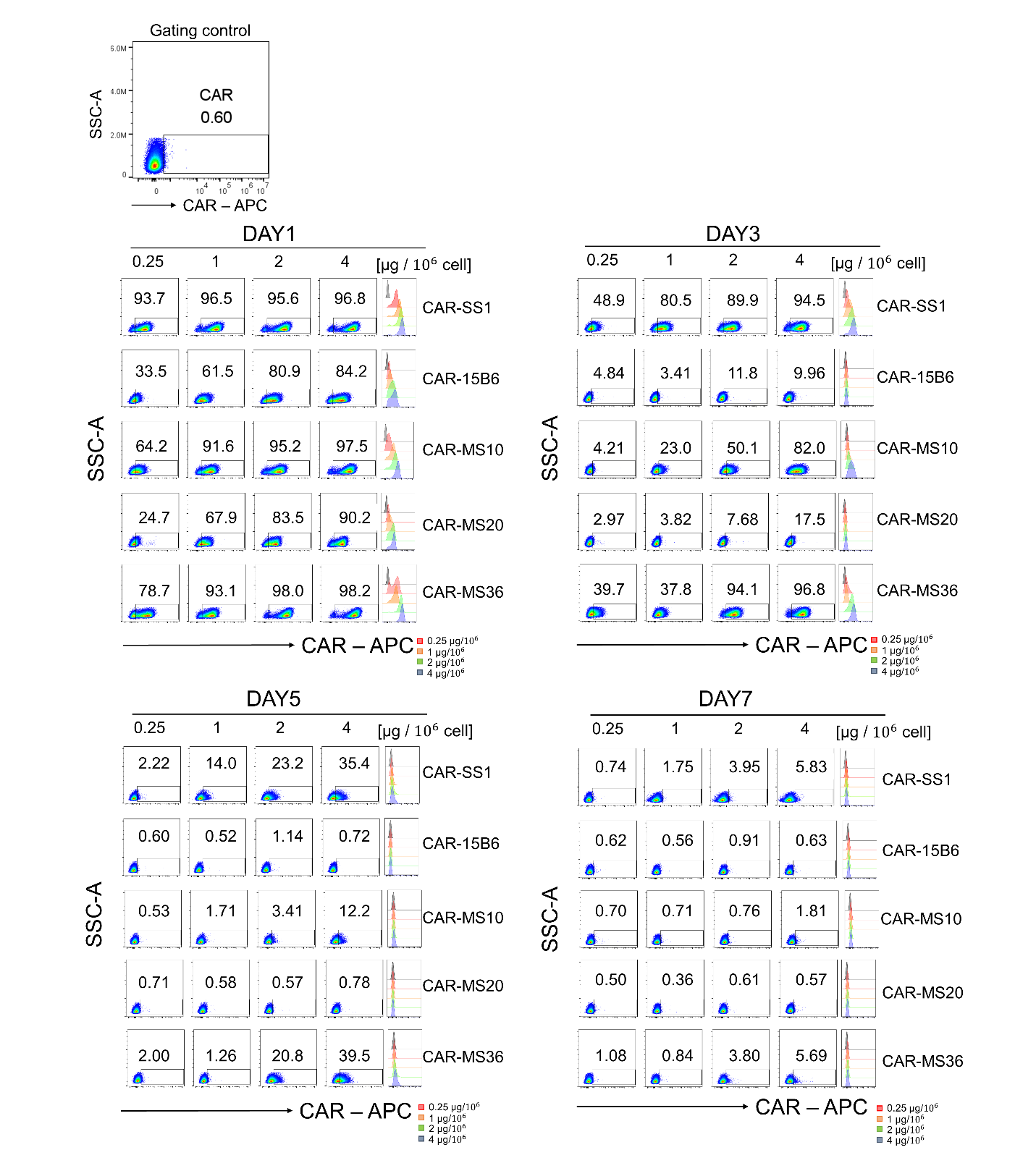

**Supplementary Figure 7. Comprehensive evaluation of CAR expression in various linCAR-NK cells**

CAR surface expression was comprehensively evaluated in various linCAR-NK cells following electroporation in a time- and dose-dependent manner. Overall, linCAR-NK cells exhibited the highest CAR expression on day 1, followed by a steep decline beginning on day 3, with expression becoming barely detectable by days 5 and 7, as measured using biotinylated matMSLN protein and APC-streptavidin at 24 h post-electroporation. CAR expression levels were positively correlated with the amount of electroporated linRNA.

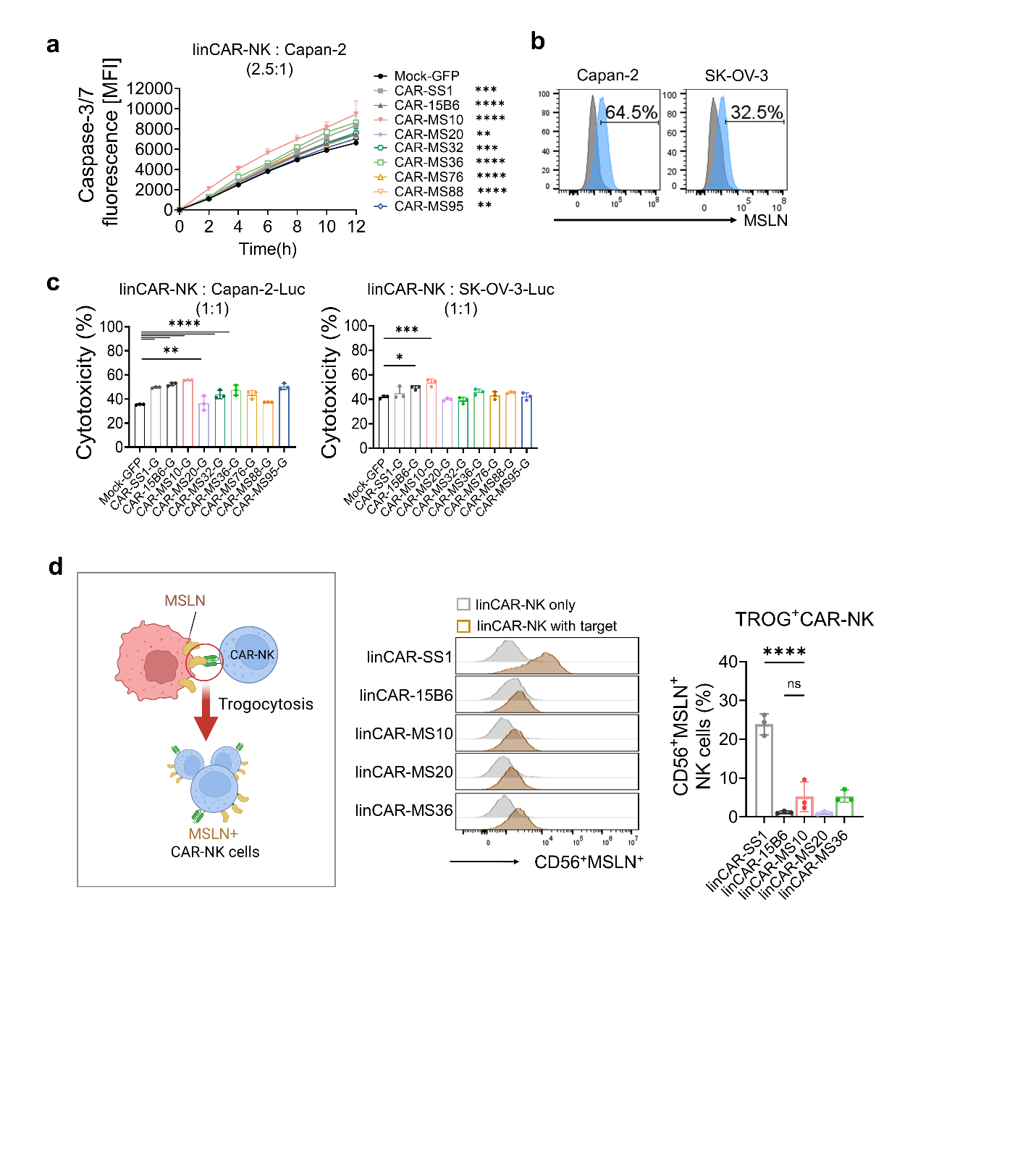

**Supplementary Figure 8.** **Functional cytotoxicity screening and trogocytic activity of linCAR-NK cells**

**(a)** Live-cell imaging of Caspase-3/7 activation in Capan-2 cells cocultured with linCAR-NK cells at a 2.5:1 E:T ratio for 12 h. **(b)** MSLN expression in Capan-2 and SK-OV-3 cells assessed by flow cytometry. **(c)** Luciferase-based cytotoxicity assay of linCAR-NK cells against Capan-2-Luc and SK-OV-3-Luc at a 1:1 E:T ratio after 12 h coculture. **(d)** MSLN antigen expression in trogocytosed linCAR-NK cells after coculture. Data were analyzed using one-way ANOVA in **(b–d)**; Data were analyzed using two-way ANOVA in **(a)** and one-way ANOVA in **(c)** and **(d)**; data are presented as mean ± s.d.; **p* < 0.05, ***p* < 0.01, ****p* < 0.001, and *****p* < 0.0001.

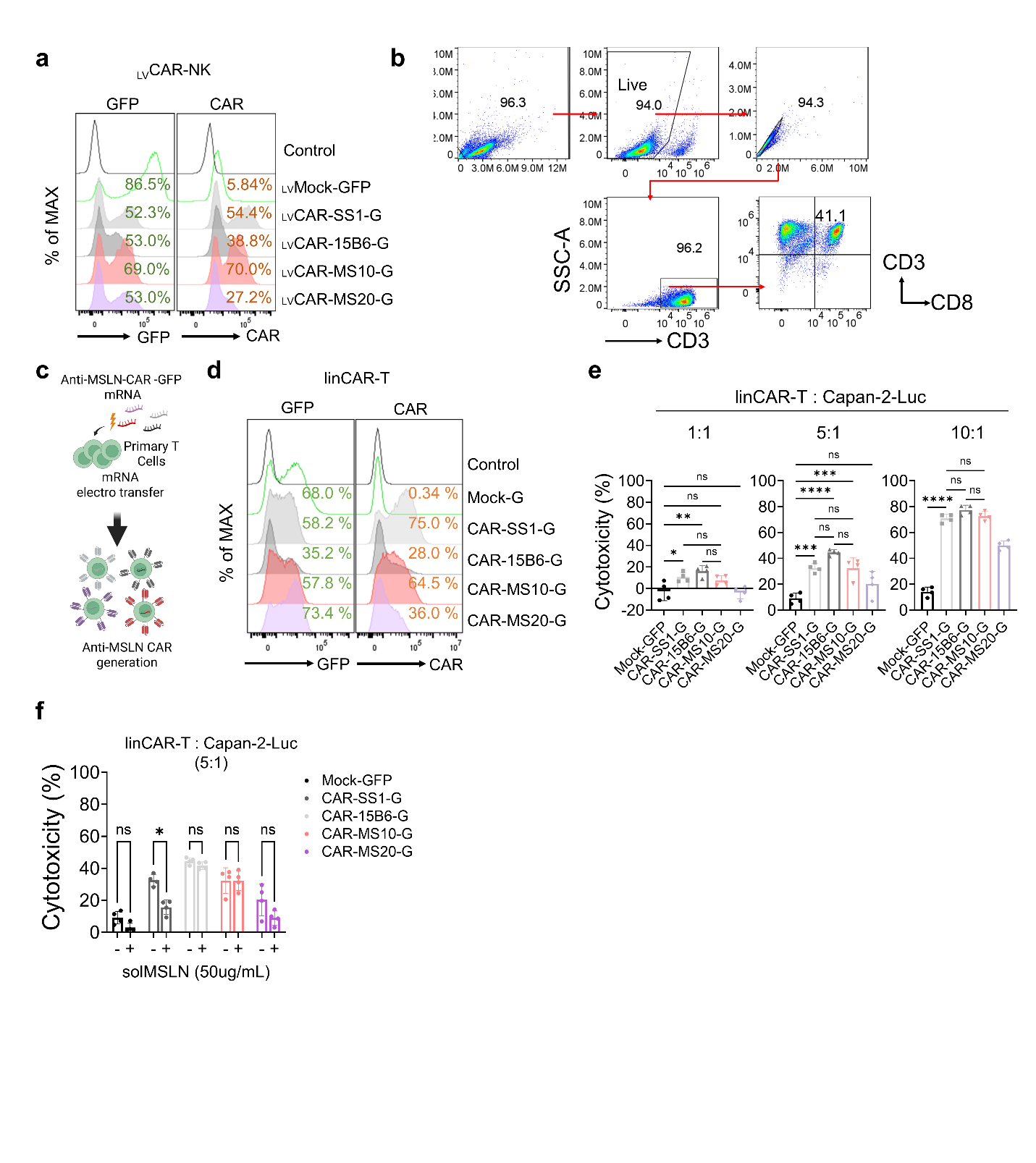

**Supplementary Figure 9.** **Comparative evaluation of _LV_CAR-NK and linCAR-T platform**
**(a)** Flow cytometry analysis of CAR expression in primary NK cells transduced with lentiviral CAR constructs targeting MSLN (CAR-SS1, CAR-15B6, CAR-MS10, and CAR-MS20). **(b)** Flow cytometry of CD3⁺CD8⁺ population after 5 days of *in vitro* expansion from PBMC-derived T cells. **(c)** Schematic of linCAR-T cell generation, involving electroporation of *in vitro* transcribed CAR linRNA into PBMC-derived T cells. **(d)** Flow cytometric analysis of GFP and CAR expression in linCAR-T cells 24 h post-electroporation. **(e)** Luciferase-based cytotoxicity of linCAR-T cells cocultured with Capan-2-Luc for 12 h at E:T ratios of 1:1, 5:1, and 10:1. **(f)** Cytotoxicity assay of linRNA-CAR-T against Capan-2-Luc cells in the presence or absence of solMSLN (50 µg/mL) for 12 h. Data were analyzed using one-way ANOVA in **(e)** and **(f)**; data are presented as mean ± s.d.; **p* < 0.05, ***p* < 0.01, ****p* < 0.001, and *****p* < 0.0001.

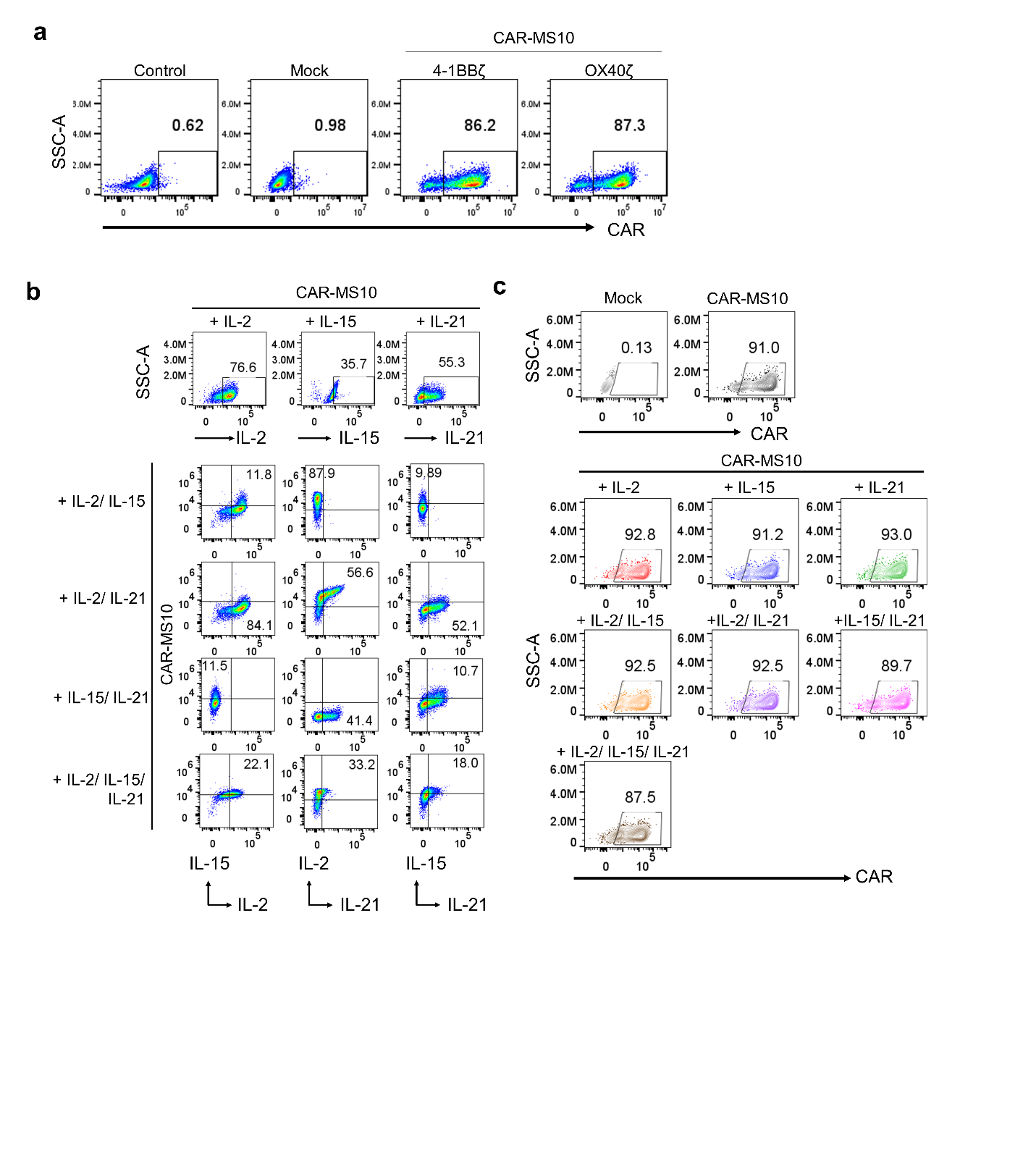

**Supplementary Figure 10.** **FACS analysis of CAR and cytokine expression for costimulatory domain optimization and codelivery strategies**

**(a)** Flow cytometry analysis of CAR expression in NK cells 12 h post-electroporation with linCAR-MS10 constructs containing 4-1BBζ and OX40ζ. **(b)** Intracellular flow cytometry of both CAR and cytokine expression in NK cells 4 h post-electroporation with linRNAs encoding CAR-MS10 and individual or combined cytokines (IL-2, IL-15, and IL-21). **(c)** Flow cytometry of GFP and CAR expression in NK cells from the same experimental groups as in **(b)**, evaluated 12 h post-electroporation.

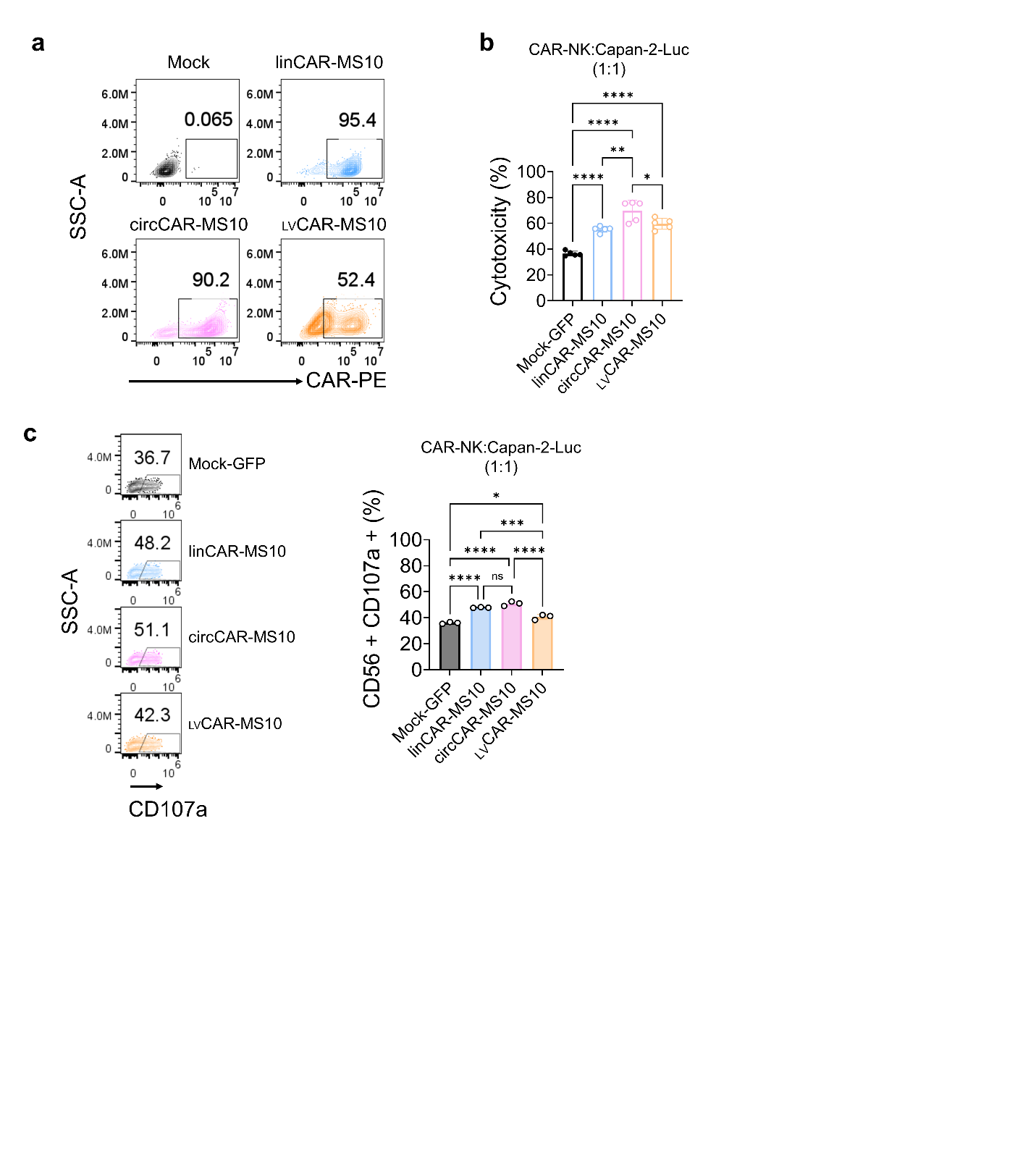

**Supplementary Figure 11. In vitro comparison of CAR platforms, including linCAR, circCAR and _LV_CAR.**

**(a)** Surface CAR expression measured 24 h after electroporation (mRNA-CAR) or 5 d after transduction (_LV_CAR), prior to coculture. **(b)** In vitro cytotoxicity of each CAR-NK group was measured 12 h after coculture with Capan-2-Luc cells at a 1:1 E:T ratio. **(c)** CD107a degranulation was assessed by intracellular staining 4 h after coculture. Data were analyzed using one-way ANOVA in **(b)** and **(c)**; data are presented as mean ± s.d.; **p* < 0.05, ***p* < 0.01, ****p* < 0.001, and *****p* < 0.0001.

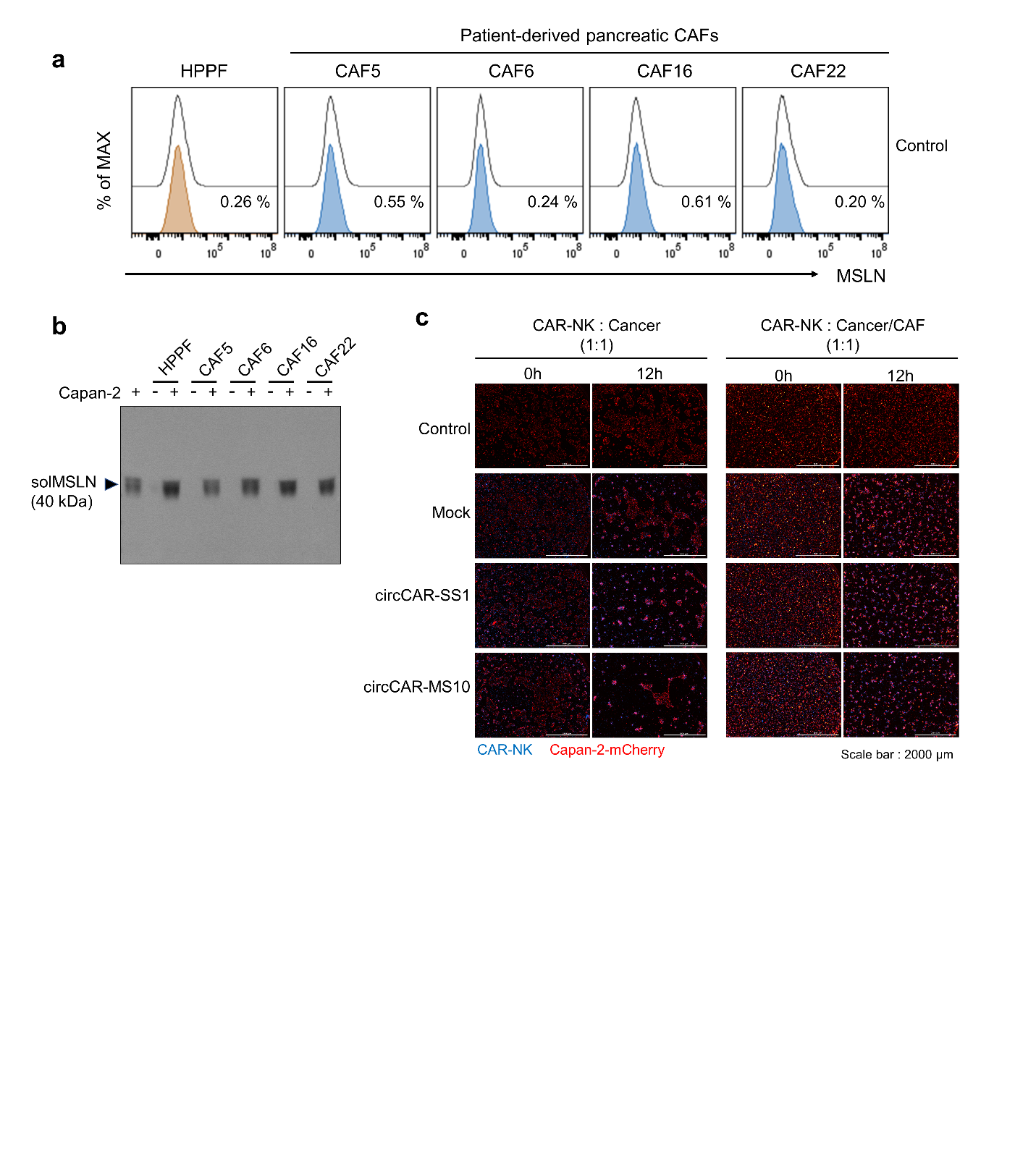

**Supplementary Figure 12.** **Enhanced solMSLN secretion in cancer–CAF coculture model**

**(a)** MSLN expression in pancreatic cancer-associated fibroblasts (CAFs; CAF5, CAF6, CAF16, and CAF22) and human primary pulmonary fibroblasts (HPPFs). **(b)** Western blot analysis of solMSLN levels in conditioned media collected from Capan-2 cells alone, CAFs alone, or cocultures of Capan-2 cells with pancreatic CAFs or HPPFs. **(c)** Representative live cell imaging shows viable cancer cells exhibiting red fluorescence (derived from Capan-2-mCherry cells) cocultured with CAR-NK cells at a 1:1 ratio for 12 h in the absence or presence of CAFs.

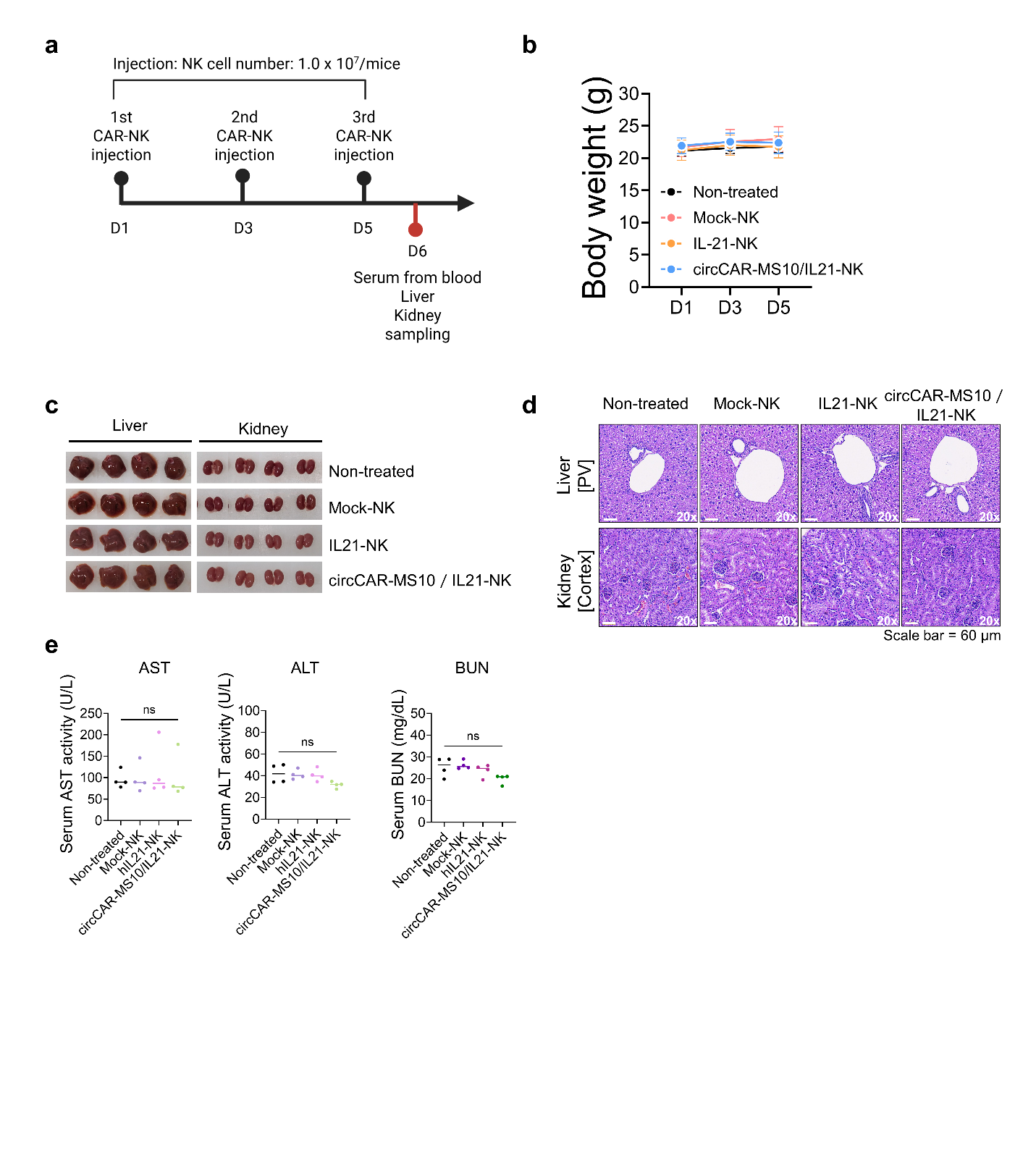

**Supplementary Figure 13. Systemic toxicity assessment following repeated circCAR-NK administration**

**(a)** Schematic of the toxicity-evaluation schedule. **(b)** Body weight measurements of mice treated with each NK cell group. **(c)** Gross morphology of liver and kidney tissues collected on day 6 after NK injection. **(d)** Histological analysis of sectioned tissues. **(e)** Serum biochemical indicators of hepatic and renal function, including aspartate aminotransferase (AST), alanine aminotransferase (ALT), and blood urea nitrogen (BUN). Data were analyzed using one-way ANOVA in **(e)**; data are presented as mean ± s.d.; ns, nonsignificant

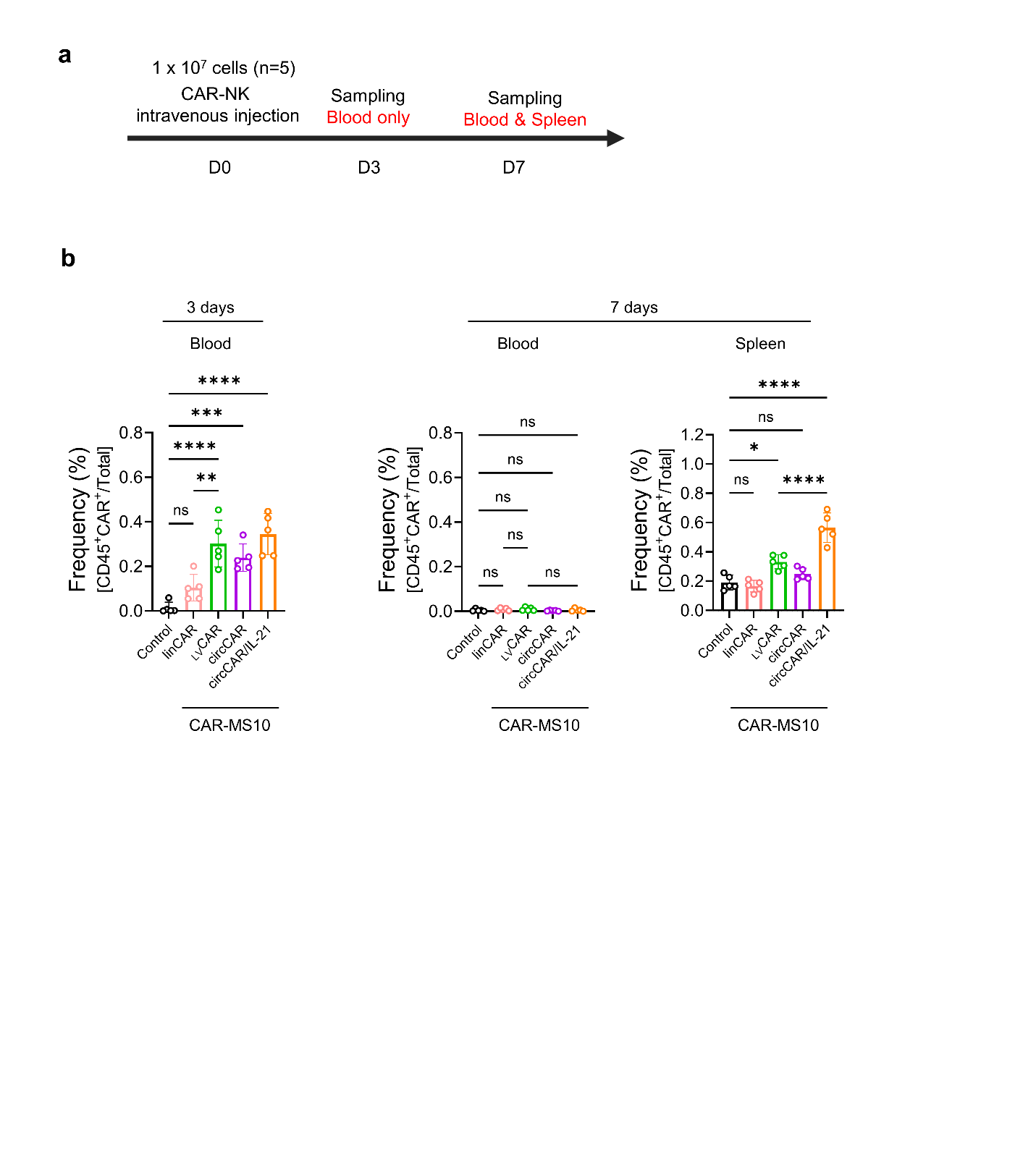
**Supplementary Figure 14. *In vivo* persistence of CAR-NK cells following intravenous infusion**

**(a)** Schematic of the experiment timeline. **(b)** Quantification of CD45^+^CAR^+^ NK cell frequencies in peripheral blood and spleen at the indicated time points. Data were analyzed using one-way ANOVA in **(b)**; data are presented as mean ± s.d.; **p* < 0.05, ***p* < 0.01, ****p* < 0.001, and *****p* < 0.0001.

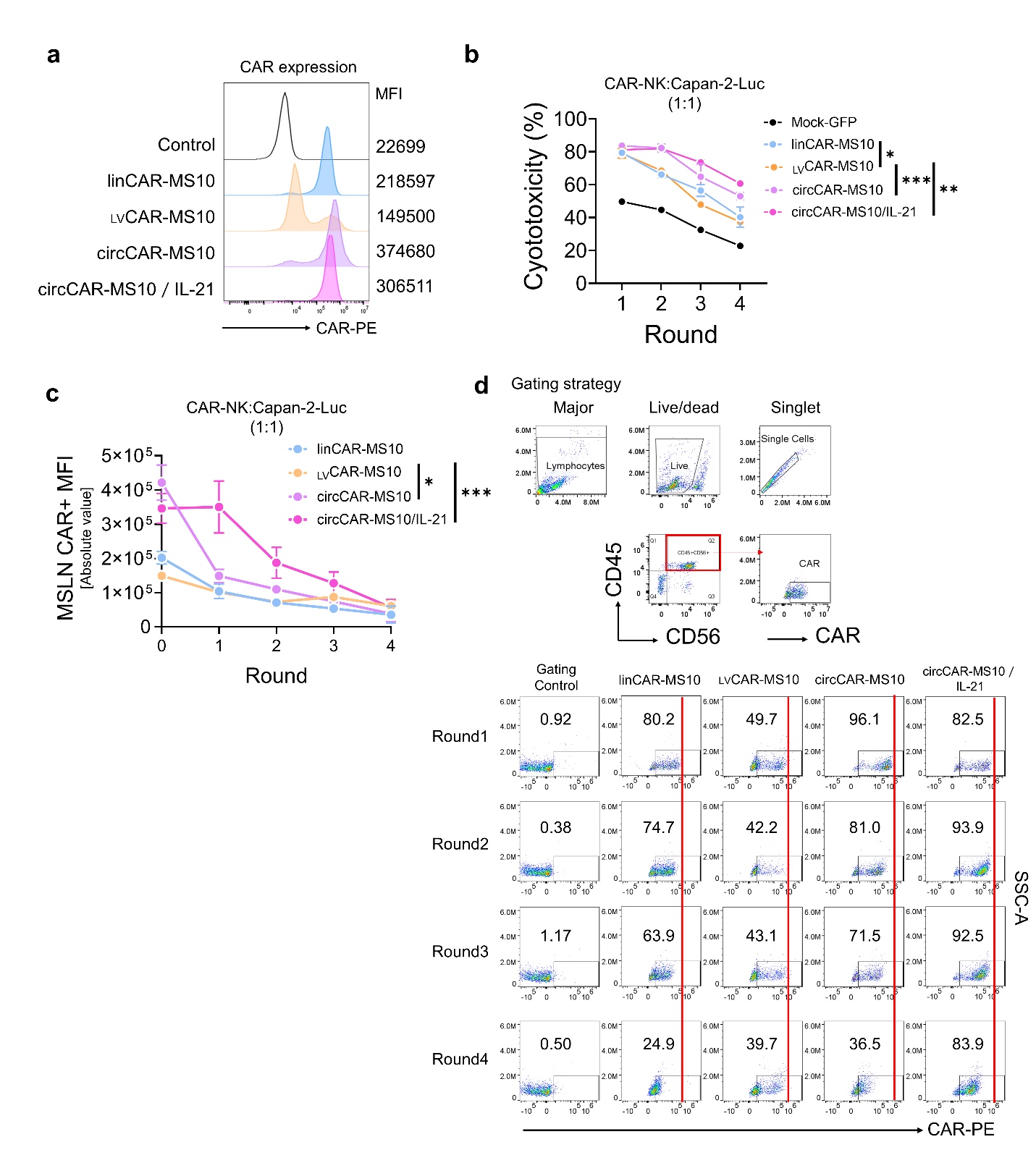

**Supplementary Figure 15. CAR-stability assay under repeated antigen encounters**

**(a)** Surface CAR expression in CAR-NK cells. **(b)** Serial killing assay of CAR-NK cells sequentially exposed to fresh Capan-2 cancer cells at 12 h intervals. **(c)** Quantification of CAR downregulation across each CAR-NK platform following sequential rounds of coculture with Capan-2-Luc target cells (Rounds 1–4). For each round, NK cells were cocultured with tumor cells for 12 h at a 1:1 E:T ratio and subsequently analyzed by FACS to assess surface CAR expression within CD45^+^CD56^+^ NK cells. **(d)** Gating strategy and dot plots showing surface CAR expression, presented as percentage of CAR population. Data were analyzed using two-way ANOVA in **(b)** and **(c)**; data are presented as mean ± s.d.; **p* < 0.05, ***p* < 0.01, ****p* < 0.001, and *****p* < 0.0001.

**Supplementary Tables**

**Supplementary Table 1.** Nucleotide and amino acid sequences of oligonucleotide primers, scFvs, and CAR constructs

| Oligonucleotide primers | V_H_ Forward | Oligonucleotide primer_sequence [5’ → 3’] |
| --- | --- | --- |
|  | Mouse VH for-1 | 5’-GAG GCC GCT AGG GCC GAG GTT CDS CTG CAA CAG TY-3’ |
|  | Mouse VH for-2 | 5’-GAG GCC GCT AGG GCC CAG GTG CAA MTG MAG SAG TC-3’ |
|  | Mouse VH for-3 | 5’-GAG GCC GCT AGG GCC GAV GTG MWG CTG GTG GAG TC-3’ |
|  | Mouse VH for-4 | 5’-GAG GCC GCT AGG GCC CAG GTT AYT CTG AAA GAG TC-3’ |
|  | Mouse VH for-5 | 5’-GAG GCC GCT AGG GCC GAK GTG CAG CTT CAG SAG TC-3’ |
|  | Mouse VH for-6 | 5’-GAG GCC GCT AGG GCC CAG ATC CAG TTS GYG CAG TC-3’ |
|  | Mouse VH for-7 | 5’-GAG GCC GCT AGG GCC CAG RTC CAA CTG CAG CAG YC-3’ |
|  | Mouse VH for-8 | 5’-GAG GCC GCT AGG GCC GAG GTG MAG CTA STT GAG WC-3’ |
|  | Mouse VH for-9 | 5’-GAG GCC GCT AGG GCC GAA GTG AAG MTT GAG GAG TC-3’ |
|  | Mouse VH for-10 | 5’-GAG GCC GCT AGG GCC GAT GTG AAC CTG GAA GTG TC-3’ |
|  | Mouse VH for-11 | 5’-GAG GCC GCT AGG GCC CAG ATK CAG CTT MAG GAG TC-3’ |
|  | Mouse VH for-12 | 5’-GAG GCC GCT AGG GCC CAG GCT TAT CTG CAG CAG TC-3’ |
|  | Mouse VH for-13 | 5’-GAG GCC GCT AGG GCC CAG GTT CAC CTA CAA CAG TC-3’ |
|  | Mouse VH for-14 | 5’-GAG GCC GCT AGG GCC CAG GTG CAG CTT GTA GAG AC-3’ |
|  | Mouse VH for-15 | 5’-GAG GCC GCT AGG GCC GAR GTG MAG CTG KTG GAG AC-3’ |
|  | V_H_ reverse |  |
|  | Mouse VH rev-1 | 5’-CGA GGA GAC GGT GAC MGT GG-3’ |
|  | Mouse VH rev-2 | 5’-CGC AGA GAC AGT GAC CAG AG-3’ |
|  | Mouse VH rev-3 | 5’-CGA GGA GAC TGT GAG AST GG-3’ |
|  | Vκ Forward |  |
|  | Mouse Vκ for-1 | 5’-GAT AAA AGA GAG GCC GCT AGG GCC GAC AWT GTT CTC ACC CAG TC-3’ |
|  | Mouse Vκ for-2 | 5’-GAT AAA AGA GAG GCC GCT AGG GCC GAC ATC CAG ATG ACA CAG WC-3’ |
|  | Mouse Vκ for-3 | 5’-GAT AAA AGA GAG GCC GCT AGG GCC GAT RTT GTG ATG ACC CAG WC-3’ |
|  | Mouse Vκ for-4 | 5’-GAT AAA AGA GAG GCC GCT AGG GCC GAC ATT STG MTG ACC CAG TC-3’ |
|  | Mouse Vκ for-5 | 5’-GAT AAA AGA GAG GCC GCT AGG GCC GAT GTT GTG VTG ACC CAA AC-3’ |
|  | Mouse Vκ for-6 | 5’-GAT AAA AGA GAG GCC GCT AGG GCC GAC ACA ACT GTG ACC CAG TC-3’ |
|  | Mouse Vκ for-7 | 5’-GAT AAA AGA GAG GCC GCT AGG GCC GAY ATT KTG CTC ACT CAG TC-3’ |
|  | Mouse Vκ for-8 | 5’-GAT AAA AGA GAG GCC GCT AGG GCC GAT ATT GTG ATR ACC CAG GM-3’ |
|  | Mouse Vκ for-9 | 5’-GAT AAA AGA GAG GCC GCT AGG GCC GAC ATT GTA ATG ACC CAA TC-3’ |
|  | Mouse Vκ for-10 | 5’-GAT AAA AGA GAG GCC GCT AGG GCC GAC ATT GTG ATG WCA CAG TC-3’ |
|  | Mouse Vκ for-11 | 5’-GAT AAA AGA GAG GCC GCT AGG GCC GAT RTC CAG ATG AMC CAG TC-3’ |
|  | Mouse Vκ for-12 | 5’-GAT AAA AGA GAG GCC GCT AGG GCC GAT GGA GAA ACA ACA CAG GC-3’ |
|  | Vκ Reverse |  |
|  | Mouse Vκ rev-1 | 5’- ACA GAT GGT GCA GCC ACC GTG CGT TTB ATT TCC AGC TTG G -3’ |
|  | Mouse Vκ rev-2 | 5’- ACA GAT GGT GCA GCC ACC GTG CGT TTT ATT TCC AAT TTT G -3’ |
|  | CH forward |  |
|  | CH for-1 | 5’-CCA CKG TCA CCG TCT CCT CGG CCT CCA CCA AGG GCC CAT CG-3’ |
|  | CH for-2 | 5’-CTC TGG TCA CTG TCT CTG CGG CCT CCA CCA AGG GCC CAT CG-3’ |
|  | CH for-3 | 5’-CCA STC TCA CAG TCT CCT CGG CCT CCA CCA AGG GCC CAT CG-3’ |
|  | CH reverse | 5’-CGA ATT CAG AAC CTC TTG GAA CTA GCA AGT-3’ |
|  | CL forward | 5’-CGT ACG GTG GCT GCA CCA TCT GT-3’ |
|  | CL reverse | 5’-AAT TAC ATG ACT CGA GCT ACT TGT CAT CGT C-3’ |
| scFv | **Clone number** | **Amino acid sequence** |
|  | CLMS10 | DIVMTQSPAIMSASPGEKVTMTCSASSSVSYMHWYQQKSGTSPKRWIYDTSRLASGVPTRFSGSGSGTSYSLTISSMEAEDAATYYCQQWSSYPLTFGAGTKLEIKRGGGGSGGGGSGGGGSQVQLQQPGAELVKPGASVKLSCKASGYTFTSYWMHWVKQRPGQGLEWIGMIHPNSGSTNYNEKFKSKATLTVDKSSSTAYMQLSSLTSEDSAVYYCARRHYYGGRYEYFDVWGTGTTVTVSS |
|  | CLMS20 | RLQMTQSPAIMSASPGEKVTMTCSASSSVSYMHWYQQKSGTSPKRWIYDTSKLASGVPARFSGSGSGTSYSLTISSMEAEDAATYYCQQWSSYPLTFGAGTKLEIKRGGGGSGGGGSGGGGSQVQLQQSGAELVKPGASVKLSCKASGYTFTSYWMHWVKQRPGQGLEWIGMIHPNSGSTNYNEKFKSKATLTVDKSSSTAYMQLSSLTSEDSAVYFCASAPIWYFDIWGTGTTVTVSS |
|  | CLMS32 | DIVMTQSHKFMSTSVGDRVSITCKASQDVGTAVAWYQQKPGQSPKALIYSASYRYSGVPDRFTGSGSGTDFTLTINNVQSEDLADYFCQQYSTYPLTFGAGTKLEIKR GGGGSGGGGSGGGGSQAYLQQSGAELVKPGASVKLSCKASGYTFTSYWMHWVKQRPGQGLEWIGMIHPNSGSSHYNEKFKSKATLTVDTSSSTAYMQLSSLTSEDSAVYYCARFFLYDYDAWFAYWGQGTLVTVSA |
|  | CLMS36 | DILLTQSPAIMSASPGEKVTMTCSASSSVSYMHWYQQKSGTSPKRWIYDTSKLASGVPARFSGSGSGTSYSLTISSMEAEDAATYYCQQWSSYPLTFGAGTKLEIKRGGGGSGGGGSGGGGSQVQLQQPGAELVKPGASVKLSCKASGYTFTSYWMHWVKQRPGHGLEWIGMIHPNSGSTHYNEKFKSKATLTVDKSSSTAYMQLSSLTSEDSAVYYCARKVWDYDWFAYWGQGTLVTVSS |
|  | CLMS76 | DIVLTQSPAIMSASPGEKVTMTCSASSSVSYMHWYQQKPGTSPKPWIYLTSNLASGVPARFSGSGSGTSYSLTISSMEAEDAATYYCQQWSSYPLTFGAGTKLEIKRGGGGSGGGGSGGGGSQAYLQQSGAELVKPGASVKLSCKASGYTFTSYWMHWVKQRPGQGLEWIGMIHPNSGSTNYNENFKSKATLTADKSSSTAYIQLSSLTSEDSAVYYCAIGWYWYFDVWGTGTTVTVSS |
|  | CLMS88 | DIVMTQSPAIMSASPGEKVTITCSASSSVSYMHWYQQKSGTSPKRWIYDTSKLASGVPARFSGSGSGTSYSLTISRMEAEDAATYYCQQWSSYPLTFGAGTKLEMKRGGGGSGGGGSGGGGSQIQLQQPGAELVKPGASVKMSCKASGYTFTSYWMHWVKQRPGQGLEWIGMIHPNSGSTIYNEKFKSKATLTVDKSSSTAYMQLSSLTSEDSAVYYCARDGHTTARYWYFDVWGTGTTVTVSA |
|  | CLMS95 | DIVMTQSPAIMSASLGERVTMTCSASSSVSYMYWYQQKSGTSPKRWIYDTSKLASGVPARFSGSGSGTSYSLTISSMEAEDAATYYCQQWSSYPLTFGAGTKLEIKRGGGGSGGGGSGGGGSQVQLQQSGAELVKPGASVKLSCKASGYTFTSYWITWVKQRPGQGLEWIGMIHPNSGSTNYNEKFKNKATLTVDKSSSTAYMQLSSLTSEDSAVYYCTILYERGYFDVWGTGTTVTVSS |
|  | SS1 | DIELTQSPAIMSASPGEKVTMTCSASSSVSYMHWYQQKSGTSPKRWIYDTSKLASGVPGRFSGSGSGNSYSLTISSVEAEDDATYYCQQWSGYPLTFGAGTKLEIRGGGGSGGGGSGGGGSQVQLQQSGPELEKPGASVKISCKASGYSFTGYTMNWVKQSHGKSLEWIGLITPYNGASSYNQKFRGKATLTVDKSSSTAYMDLLSLTSEDSAVYFCARGGYDGRGFDYWGQGTTVTVSS |
|  | 15B6 | QAVVTQESALTTSPGETVTLTCRSSTGAVTTGNYPNWVQEKPDHLFTGLIAGTNNRAPGVPARFSGSLIGDKAALTITGAQTEDEAIYFCALWFSSHWVFGGGTKLTVLGGGGSGGGGSGGGGSEVQLQQSGPVLVKPGASVKISCKASGYSFTGYYMHWVRQSLVKRLEWIGRINPYTGVPSYKHNFKDKASLTVDKSSSTAYMELHSLTSEDSAVYYCARELGGYWGQGTTLTVSS |
| scFv | **Clone number** | **Nucleotide sequence** |
|  | CLMS10 | 5’-GAT ATT GTG ATG ACC CAG TCT CCA GCA ATC ATG TCT GCA TCT CCA GGG GAG AAG GTC ACC ATG ACC TGC AGT GCC AGC TCA AGT GTA AGT TAC ATG CAC TGG TAC CAG CAG AAG TCA GGC ACC TCC CCC AAA AGA TGG ATT TAT GAC ACA TCC AGA CTG GCT TCT GGA GTC CCT ACT CGC TTC AGT GGC AGT GGG TCT GGG ACC TCT TAC TCT CTC ACA ATC AGC AGC ATG GAG GCT GAA GAT GCT GCC ACT TAT TAC TGC CAG CAG TGG AGT AGT TAC CCG CTC ACG TTC GGT GCT GGG ACC AAG CTG GAA ATC AAA CGC GGA GGT GGG GGC AGT GGA GGC GGT GGT AGT GGA GGC GGA GGA AGT CAG GTC CAA CTG CAG CAG CCT GGG GCT GAG CTG GTA AAG CCT GGG GCT TCA GTG AAG TTG TCC TGC AAG GCT TCT GGC TAC ACT TTC ACC AGC TAC TGG ATG CAC TGG GTG AAG CAG AGG CCT GGA CAA GGC CTT GAG TGG ATT GGA ATG ATT CAT CCT AAT AGT GGT AGT ACT AAC TAC AAT GAG AAG TTC AAG AGC AAG GCC ACA CTG ACT GTA GAC AAA TCC TCC AGC ACA GCC TAC ATG CAA CTC AGC AGC CTG ACA TCT GAG GAC TCT GCG GTC TAT TAC TGT GCA AGA AGG CAT TAC TAC GGT GGT AGG TAC GAG TAC TTC GAT GTC TGG GGC ACA GGG ACC ACT GTC ACC GTC TCC TCG-3’ |
|  | CLMS20 | 5’-CGA CTC CAG ATG ACA CAG TCT CCA GCA ATC ATG TCT GCA TCT CCA GGG GAG AAG GTC ACC ATG ACC TGC AGT GCC AGC TCA AGT GTA AGT TAC ATG CAC TGG TAC CAG CAG AAG TCA GGC ACC TCC CCC AAG AGA TGG ATT TAT GAC ACA TCC AAA CTG GCT TCT GGA GTC CCT GCT CGC TTC AGT GGC AGT GGG TCT GGG ACC TCT TAC TCT CTC ACA ATC AGC AGC ATG GAG GCT GAA GAT GCT GCC ACT TAT TAC TGC CAG CAG TGG AGT AGT TAC CCG CTC ACG TTC GGT GCT GGG ACC AAG CTG GAA ATA AAA CGC GGA GGT GGG GGC AGT GGA GGC GGT GGT AGT GGA GGC GGA GGA AGT CAG GTC CAA CTG CAG CAG TCT GGG GCT GAA TTG GTA AAG CCT GGG GCT TCA GTG AAG TTG TCC TGC AAG GCT TCT GGC TAC ACT TTC ACC AGC TAC TGG ATG CAC TGG GTG AAG CAG AGG CCT GGA CAA GGC CTT GAG TGG ATT GGA ATG ATT CAT CCT AAT AGT GGT AGT ACT AAC TAC AAT GAG AAG TTC AAG AGC AAG GCC ACA CTG ACT GTA GAC AAA TCC TCC AGC ACA GCC TAC ATG CAG CTC AGC AGC CTG ACA TCT GAG GAC TCT GCG GTC TAT TTC TGT GCA AGT GCC CCA ATC TGG TAC TTC GAT ATC TGG GGC ACA GGG ACC ACG GTC ACC GTC TCC TCG-3’ |
|  | CLMS32 | 5’-GAT ATT GTG ATG ACC CAG TCT CAC AAA TTC ATG TCC ACA TCA GTA GGA GAC AGG GTC AGC ATC ACC TGC AAG GCC AGT CAG GAT GTG GGT ACT GCT GTA GCC TGG TAT CAA CAG AAA CCA GGG CAA TCT CCT AAA GCA CTG ATT TAC TCG GCA TCC TAC CGG TAC AGT GGA GTC CCT GAT CGC TTC ACA GGC AGT GGA TCT GGG ACA GAT TTC ACT CTC ACC ATT AAC AAT GTG CAG TCT GAA GAC TTG GCA GAT TAT TTC TGT CAG CAA TAT AGC ACC TAT CCT CTC ACG TTC GGT GCT GGG ACC AAG CTG GAA ATA AAA CGC GGA GGT GGG GGC AGT GGA GGC GGT GGT AGT GGA GGC GGA GGA AGT CAG GCT TAT CTG CAG CAG TCT GGG GCT GAG CTG GTA AAG CCT GGG GCT TCA GTG AAG TTG TCC TGC AAG GCT TCT GGC TAC ACT TTC ACC AGC TAC TGG ATG CAC TGG GTG AAG CAG AGG CCT GGA CAA GGC CTT GAG TGG ATT GGA ATG ATT CAT CCT AAT AGT GGT AGT TCT CAC TAC AAT GAG AAG TTC AAG AGC AAG GCC ACA CTG ACT GTA GAC ACA TCC TCC AGC ACA GCC TAC ATG CAG CTC AGC AGC CTG ACA TCT GAG GAC TCT GCG GTC TAT TAC TGT GCA AGA TTT TTC CTC TAT GAT TAC GAC GCC TGG TTT GCT TAC TGG GGC CAA GGG ACT CTG GTC ACT GTC TCT GCG-3’ |
|  | CLMS36 | 5’-GAC ATT CTG CTG ACC CAG TCT CCA GCA ATC ATG TCT GCA TCT CCA GGG GAG AAG GTC ACC ATG ACC TGC AGT GCC AGC TCA AGT GTA AGT TAC ATG CAC TGG TAC CAG CAG AAG TCA GGC ACC TCC CCC AAA AGA TGG ATT TAT GAC ACA TCC AAA CTG GCT TCT GGA GTC CCT GCT CGC TTC AGT GGC AGT GGG TCT GGG ACC TCT TAC TCT CTC ACA ATC AGC AGC ATG GAG GCT GAA GAT GCT GCC ACT TAT TAC TGC CAG CAG TGG AGT AGT TAC CCG CTC ACG TTC GGT GCT GGG ACC AAG CTG GAA ATA AAA CGC GGA GGT GGG GGC AGT GGA GGC GGT GGT AGT GGA GGC GGA GGA AGT CAG GTC CAA CTG CAG CAG CCT GGG GCT GAG CTG GTA AAG CCT GGG GCT TCA GTG AAG TTG TCC TGC AAG GCT TCT GGC TAC ACT TTC ACC AGC TAC TGG ATG CAC TGG GTG AAG CAA AGG CCT GGA CAC GGC CTT GAG TGG ATT GGA ATG ATT CAT CCT AAT AGT GGT AGT ACT CAC TAC AAT GAG AAG TTC AAG AGC AAG GCC ACA CTG ACT GTA GAC AAA TCC TCC AGC ACA GCC TAC ATG CAA CTC AGC AGC CTG ACA TCT GAG GAC TCT GCG GTC TAT TAC TGT GCA AGA AAA GTT TGG GAT TAC GAC TGG TTT GCT TAC TGG GGC CAA GGG ACT CTG GTC ACA GTC TCC TCG-3’ |
|  | CLMS76 | 5’-GAC ATT GTG CTC ACT CAG TCT CCA GCA ATC ATG TCT GCA TCT CCA GGG GAG AAG GTC ACC ATG ACC TGC AGT GCC AGC TCA AGT GTA AGT TAC ATG CAC TGG TAC CAG CAG AAG CCA GGC ACC TCC CCC AAA CCC TGG ATT TAT CTC ACA TCC AAC CTG GCT TCT GGA GTC CCT GCT CGC TTC AGT GGC AGT GGG TCT GGG ACC TCT TAC TCT CTC ACA ATC AGC AGC ATG GAG GCT GAA GAT GCT GCC ACT TAT TAC TGC CAG CAG TGG AGT AGT TAC CCA CTC ACG TTC GGT GCT GGG ACC AAG CTG GAA ATC AAA CGC GGA GGT GGG GGC AGT GGA GGC GGT GGT AGT GGA GGC GGA GGA AGT CAG GCT TAT CTG CAG CAG TCT GGG GCT GAG CTG GTA AAG CCT GGG GCT TCA GTG AAG TTG TCC TGC AAG GCT TCT GGC TAC ACT TTC ACC AGC TAC TGG ATG CAC TGG GTG AAG CAG AGG CCT GGA CAA GGC CTT GAG TGG ATT GGA ATG ATT CAT CCT AAT AGT GGT AGT ACT AAC TAC AAT GAG AAC TTC AAG AGC AAG GCC ACA CTG ACT GCA GAC AAA TCC TCC AGC ACT GCC TAC ATT CAG CTC AGC AGT CTG ACA TCT GAG GAC TCT GCG GTC TAT TAC TGT GCA ATA GGA TGG TAC TGG TAC TTC GAT GTC TGG GGC ACA GGG ACC ACG GTC ACA GTC TCC TCG-3’ |
|  | CLMS88 | 5’-GAC ATT GTG ATG ACC CAG TCT CCA GCA ATC ATG TCT GCA TCT CCA GGG GAG AAG GTC ACC ATA ACC TGC AGT GCC AGC TCA AGT GTA AGT TAC ATG CAC TGG TAC CAG CAG AAG TCA GGC ACC TCC CCC AAA AGA TGG ATT TAT GAC ACA TCC AAA CTG GCT TCT GGA GTC CCT GCT CGC TTC AGT GGC AGT GGG TCT GGG ACC TCT TAC TCT CTC ACA ATC AGC CGA ATG GAG GCT GAA GAT GCT GCC ACT TAT TAC TGC CAG CAG TGG AGT AGT TAC CCG CTC ACG TTC GGT GCT GGG ACC AAG CTG GAA ATG AAA CGC GGA GGT GGG GGC AGT GGA GGC GGT GGT AGT GGA GGC GGA GGA AGT CAG ATC CAA CTG CAG CAG CCT GGG GCT GAG CTT GTG AAG CCT GGG GCT TCA GTG AAG ATG TCC TGC AAG GCT TCT GGC TAC ACC TTC ACC AGC TAC TGG ATG CAC TGG GTG AAG CAG AGG CCT GGA CAA GGC CTT GAG TGG ATT GGA ATG ATT CAT CCT AAT AGT GGT AGT ACT ATC TAC AAT GAG AAG TTC AAG AGC AAG GCC ACA CTG ACT GTA GAC AAA TCC TCC AGC ACA GCC TAC ATG CAG CTC AGC AGC CTG ACA TCT GAG GAC TCT GCG GTC TAT TAT TGT GCA AGA GAC GGG CAT ACT ACG GCC CGC TAC TGG TAC TTC GAT GTC TGG GGC ACA GGG ACC ACT GTC ACC GTC TCT GCG-3’ |
|  | CLMS95 | 5’-GAT ATT GTG ATG ACC CAG TCT CCA GCA ATC ATG TCT GCA TCT CTA GGG GAA CGG GTC ACC ATG ACC TGC AGT GCC AGC TCA AGT GTA AGT TAC ATG TAC TGG TAC CAG CAG AAG TCA GGC ACC TCC CCC AAA AGA TGG ATT TAT GAC ACA TCC AAA CTG GCT TCT GGA GTC CCT GCT CGC TTC AGT GGC AGT GGG TCT GGG ACC TCT TAC TCT CTC ACA ATC AGC AGC ATG GAG GCT GAA GAT GCT GCC ACT TAT TAC TGC CAG CAG TGG AGT AGT TAC CCG CTC ACG TTC GGT GCT GGG ACC AAG CTG GAA ATA AAA CGT GGA GGT GGG GGC AGT GGA GGC GGT GGT AGT GGA GGC GGA GGA AGT CAG GTC CAA CTG CAG CAG TCT GGG GCT GAG CTG GTA AAG CCT GGG GCT TCA GTG AAG TTG TCC TGC AAG GCT TCT GGC TAC ACC TTC ACC AGC TAC TGG ATA ACC TGG GTG AAG CAG AGG CCT GGA CAA GGC CTT GAG TGG ATT GGA ATG ATT CAT CCT AAT AGT GGT AGT ACT AAC TAC AAT GAG AAG TTC AAG AAC AAG GCC ACA CTG ACT GTA GAC AAA TCC TCC AGC ACA GCC TAC ATG CAA CTC AGC AGC CTG ACA TCT GAG GAC TCT GCA GTC TAT TAC TGT ACA ATA CTC TAC GAG AGA GGG TAC TTC GAT GTC TGG GGC ACA GGG ACC ACT GTC ACC GTC TCC TCG-3’ |
|  | SS1 | 5’-GAT ATC GAA CTC ACT CAG TCC CCA GCA ATC ATG TCC GCT TCA CCG GGA GAA AAG GTG ACC ATG ACT TGC TCG GCC TCC TCG TCC GTG TCA TAC ATG CAC TGG TAC CAA CAA AAA TCG GGG ACC TCC CCT AAG AGA TGG ATC TAC GAT ACC AGC AAA CTG GCT TCA GGC GTG CCG GGA CGC TTC TCG GGT TCG GGG AGC GGA AAT TCG TAT TCG TTG ACC ATT TCG TCC GTG GAA GCC GAG GAC GAC GCA ACT TAT TAC TGC CAA CAG TGG TCA GGC TAC CCG CTC ACT TTC GGA GCC GGC ACT AAG CTG GAG ATC AGG GGA GGC GGA GGG AGC GGA GGA GGA GGC AGC GGA GGT GGA GGG TCG CAA GTC CAG CTC CAG CAG TCG GGC CCA GAG TTG GAG AAG CCT GGG GCG AGC GTG AAG ATC TCA TGC AAA GCC TCA GGC TAC TCC TTT ACT GGA TAC ACG ATG AAT TGG GTG AAA CAG TCG CAT GGA AAG TCA CTG GAA TGG ATC GGT CTG ATT ACG CCC TAC AAC GGC GCC TCC AGC TAC AAC CAG AAG TTC AGG GGA AAG GCG ACC CTT ACT GTC GAC AAG TCG TCA AGC ACC GCC TAC ATG GAC CTC CTG TCC CTG ACC TCC GAA GAT AGC GCG GTC TAC TTT TGT GCA CGC GGA GGT TAC GAT GGA CGG GGA TTC GAC TAC TGG GGC CAG GGA ACC ACT GTC ACC GTG TCG AGC-3’ |
|  | 15B6 | 5’-CAA GCC GTA GTA ACA CAA GAG TCA GCA CTT ACA ACC AGT CCC GGG GAA ACG GTC ACC TTG ACT TGC CGG TCA AGC ACG GGC GCT GTC ACC ACC GGT AAC TAT CCC AAT TGG GTG CAA GAG AAG CCT GAC CAC CTG TTC ACG GGA CTC ATC GCG GGG ACA AAC AAT AGG GCA CCT GGG GTC CCG GCG AGG TTT AGT GGG AGC CTG ATC GGC GAC AAA GCA GCC CTC ACC ATT ACC GGG GCG CAA ACT GAA GAC GAG GCT ATT TAC TTC TGT GCG CTT TGG TTC TCC TCT CAC TGG GTC TTC GGC GGT GGA ACA AAA CTG ACG GTC CTT GGA GGT GGA GGG AGT GGA GGC GGT GGT TCT GGA GGC GGC GGG AGC GAG GTT CAG CTC CAA CAA AGT GGT CCA GTG TTG GTG AAG CCA GGT GCC TCT GTA AAG ATC AGC TGT AAG GCT AGT GGC TAT TCC TTC ACG GGA TAC TAC ATG CAC TGG GTT CGC CAA AGC CTC GTA AAA CGG CTC GAG TGG ATC GGT AGG ATA AAC CCG TAT ACG GGG GTT CCC TCT TAT AAG CAT AAT TTT AAA GAC AAG GCC AGC CTG ACT GTA GAC AAG TCA AGC TCA ACT GCC TAT ATG GAG CTC CAT AGC CTG ACA TCC GAG GAT TCC GCA GTG TAC TAC TGT GCG CGA GAA CTT GGC GGT TAC TGG GGG CAG GGT ACT ACC CTT ACG GTA AGT AGT-3’ |
|  | **Name** | **Amino acid sequence** |
| CAR construct | Mock-GFP | MAPAMEIECRITGTLNGVEFELVGGGEGTPKQGRMTNKMKSTKGALTFSPYLLSHVMGYGFYHFGTYPSGYENPFLHAINNGGYTNTRIEKYEDGGVLHVSFSYRYEAGRVIGDFKVVGTGFPEDSVIFTDKIIRSNATVEHLHPMGDNVLVGSFARTFSLRDGGYYSFVVDSHMHFKSAIHPSILQNGGPMFAFRRVEELHSNTELGIVEYQHAFKTPIAFA |
|  | CD8 leader | MALPVTALLLPLALLLHAARP |
|  | CD8 hinge | TTTPAPRPPTPAPTIASQPLSLRPEACRPAAGGAVHTRGLDFACD |
|  | CD8 transmembrane | IYIWAPLAGTCGVLLLSLVITLYC |
|  | NKG2D transmembrane | PFFFCCFIAVAMGIRFIIMVT |
|  | 4-1BB | KRGRKKLLYIFKQPFMRPVQTTQEEDGCSCRFPEEEEGGCEL |
|  | CD3ζ | RVKFSRSADAPAYKQGQNQLYNELNLGRREEYDVLDKRRGRDPEMGGKPRRKNPQEGLYNELQKDKMAEAYSEIGMKGERRRGKGHDGLYQGLSTATKDTYDALHMQALPPR |
|  | OX40 | ALYLLRRDQRLPPDAHKPPGGGSFRTPIQEEQADAHSTLAKI |
|  | 2B4 | WRRKRKEKQSETSPKEFLTIYEDVKDLKTRRNHEQEQTFPGGGSTIYSMIQSQSSAPTSQEPAYTLYSLIQPSRKSGSRKRNHSPSFNSTIYEVIGKSQPKAQNPARLSRKELENFDVYS |
|  | DAP10 | LCARPRRSPAQEDGKVYINMPGRG |
|  | DAP12 | YFLGRLVPRGRGAAEAATRKQRITETESPYQELQGQRSDVYSDLNTQRPYYK |
|  | **Name** | **Nucleotide sequence** |
| CAR construct | Mock-GFP | 5’-ATG GCC CCC GCC ATG GAG ATC GAG TGC CGC ATC ACC GGC ACC CTG AAC GGC GTG GAG TTC GAG CTG GTG GGC GGC GGA GAG GGC ACC CCC AAG CAG GGC CGC ATG ACC AAC AAG ATG AAG AGC ACC AAA GGC GCC CTG ACC TTC AGC CCC TAC CTG CTG AGC CAC GTG ATG GGC TAC GGC TTC TAC CAC TTC GGC ACC TAC CCC AGC GGC TAC GAG AAC CCC TTC CTG CAC GCC ATC AAC AAC GGC GGC TAC ACC AAC ACC CGC ATC GAG AAG TAC GAG GAC GGC GGC GTG CTG CAC GTG AGC TTC AGC TAC CGC TAC GAG GCC GGC CGC GTG ATC GGC GAC TTC AAG GTG GTG GGC ACC GGC TTC CCC GAG GAC AGC GTG ATC TTC ACC GAC AAG ATC ATC CGC AGC AAC GCC ACC GTG GAG CAC CTG CAC CCC ATG GGC GAT AAC GTG CTG GTG GGC AGC TTC GCC CGC ACC TTC AGC CTG CGC GAC GGC GGC TAC TAC AGC TTC GTG GTG GAC AGC CAC ATG CAC TTC AAG AGC GCC ATC CAC CCC AGC ATC CTG CAG AAC GGG GGC CCC ATG TTC GCC TTC CGC CGC GTG GAG GAG CTG CAC AGC AAC ACC GAG CTG GGC ATC GTG GAG TAC CAG CAC GCC TTC AAG ACC CCC ATT GCC TTC GCC-3’ |
|  | CD8 leader | 5’-ATG GCC CTC CCC GTG ACC GCC CTC CTA CTG CCA CTG GCT TTG CTG CTC CAT GCC GCT AGA CCC-3’ |
|  | CD8 hinge | 5’-ACT ACG ACC CCC GCG CCA CGC CCA CCC ACG CCG GCT CCC ACC ATC GCA TCG CAA CCA CTG AGC CTC AGA CCC GAA GCA TGC CGG CCC GCT GCG GGA GGA GCG GTG CAC ACA AGA GGC CTG GAC TTC GCC TGC GAC-3’ |
|  | CD8 transmembrane | 5’-ATC TAC ATC TGG GCC CCC CTG GCC GGC ACC TGC GGC GTG CTG CTG CTT AGT TTA GTC ATC ACC CTG TAC TGC-3’ |
|  | NKG2D transmembrane | 5’-CCC TTC TTC TTT TGT TGT TTT ATA GCC GTC GCC ATG GGG ATA AGA TTT ATC ATC ATG GTG ACG-3’ |
|  | 4-1BB | 5’-AAG AGA GGC AGA AAG AAG CTG CTG TAC ATC TTC AAG CAG CCC TTC ATG AGA CCC GTG CAG ACC ACC CAA GAG GAG GAC GGC TGC AGC TGC AGA TTC CCC GAG GAG GAG GAG GGC GGC TGC GAG CTG-3’ |
|  | CD3ζ | 5’-AGA GTG AAG TTC AGC AGA AGC GCC GAC GCC CCC GCC TAC AAG CAA GGG CAG AAT CAG CTG TAT AAC GAA CTC AAC CTG GGC AGA AGA GAG GAG TAC GAC GTG CTG GAC AAG AGA AGA GGC AGA GAC CCC GAG ATG GGC GGC AAG CCT AGA AGA AAG AAC CCC CAA GAG GGC CTG TAC AAC GAG CTG CAA AAG GAC AAG ATG GCC GAG GCC TAC AGC GAG ATC GGC ATG AAG GGC GAG AGA AGA AGA GGC AAG GGC CAC GAC GGC CTG TAC CAA GGC CTG AGC ACC GCC ACC AAG GAC ACC TAC GAC GCC CTG CAC ATG CAA GCC CTG CCC CCT AGA-3’ |
|  | OX40 | 5’-GCA CTC TAT TTA CTG AGA CGG GAT CAA CGC CTT CCC CCT GAC GCA CAT AAA CCA CCC GGC GGC GGT AGC TTT AGA ACA CCA ATT CAG GAA GAA CAA GCA GAT GCT CAT TCT ACG CTT GCA AAG ATT-3’ |
|  | 2B4 | 5’-TGG AGA CGA AAA CGC AAA GAG AAA CAA TCT GAA ACA TCC CCA AAA GAG TTC CTC ACT ATA TAT GAG GAC GTG AAA GAC TTA AAG ACG CGC CGA AAC CAT GAA CAA GAA CAA ACA TTC CCG GGC GGC GGA TCC ACG ATA TAT AGC ATG ATT CAA AGC CAA TCC AGC GCC CCT ACC AGC CAG GAG CCA GCC TAC ACC CTG TAC TCT CTT ATC CAA CCC TCT CGC AAA AGT GGT AGC CGA AAA CGC AAT CAT TCT CCC AGC TTT AAC TCC ACA ATA TAC GAG GTC ATC GGT AAA TCC CAG CCA AAG GCG CAA AAT CCC GCC CGG CTT TCC AGG AAG GAA CTC GAA AAT TTC GAC GTC TAC TCT-3’ |
|  | DAP10 | 5’-CTC TGT GCC AGA CCT CGT AGG AGT CCG GCG CAA GAG GAC GGG AAG GTT TAT ATT AAT ATG CCG GGT AGA GGG-3’ |
|  | DAP12 | 5’-TAT TTT CTC GGT CGC CTC GTG CCA AGG GGT CGC GGC GCC GCT GAA GCT GCT ACT CGT AAG CAA AGG ATT ACC GAA ACG GAA TCT CCG TAC CAA GAA TTA CAA GGA CAA CGG AGT GAC GTG TAT AGT GAT TTG AAT ACC CAA CGC CCA TAC TAT AAG-3’ |

**Supplementary Table 2.** EC₅₀ values of anti-MSLN antibodies determined by ELISA

|  | EC_50_ (nM) | | |
| --- | --- | --- | --- |
| Clone ID | matMSLN | solMSLN | matMSLN vs solMSLN |
| CLMS10 | 18.6 $\pm$ 1.1 | > 1000 | > 53.7 |
| CLMS20 | 116.2 $\pm$ 33.3 | n.d. | selective |
| CLMS32 | 1.7 $\pm$ 0.1 | 12.4 $\pm$ 1.4 | 7.29 |
| CLMS36 | 4.7 $\pm$ 2.3 | 3.0 $\pm$ 0.2 | 0.63 |
| CLMS76 | 2.9 $\pm$ 0.06 | 26.0 $\pm$ 9.9* | 8.97 |
| CLMS88 | 151.1 $\pm$ 33.3 | n.d. | selective |
| CLMS95 | 26.1 $\pm$ 5.3 | n.d. | selective |
| SS1 | 3.1 $\pm$ 0.11 | 2.7 $\pm$ 0.2 | 0.86 |
| 15B6 | 180.9 $\pm$ 83.6 | n.d. | selective |

* relatively low saturation signal; n.d, binding not detected

**Supplementary Table 3.** EC₅₀ values of anti-MSLN antibodies determined by cell binding assay

| Clone ID | EC_50_ (nM) |
| --- | --- |
| CLMS10 | 7.1 $\pm$ 0.1 |
| CLMS20 | > 1000 |
| CLMS32 | 28.2 $\pm$ 1.9 |
| CLMS36 | 7.8 $\pm$ 0.1 |
| CLMS76 | 146.2 $\pm$ 12.1 |
| CLMS88 | > 1000 |
| CLMS95 | 21.1 $\pm$ 3.6 |
| SS1 | 2.8 $\pm$ 0.6 |
| 15B6 | 161.8 $\pm$ 30.3 |

**Uncropped western blots and agarose gel image**

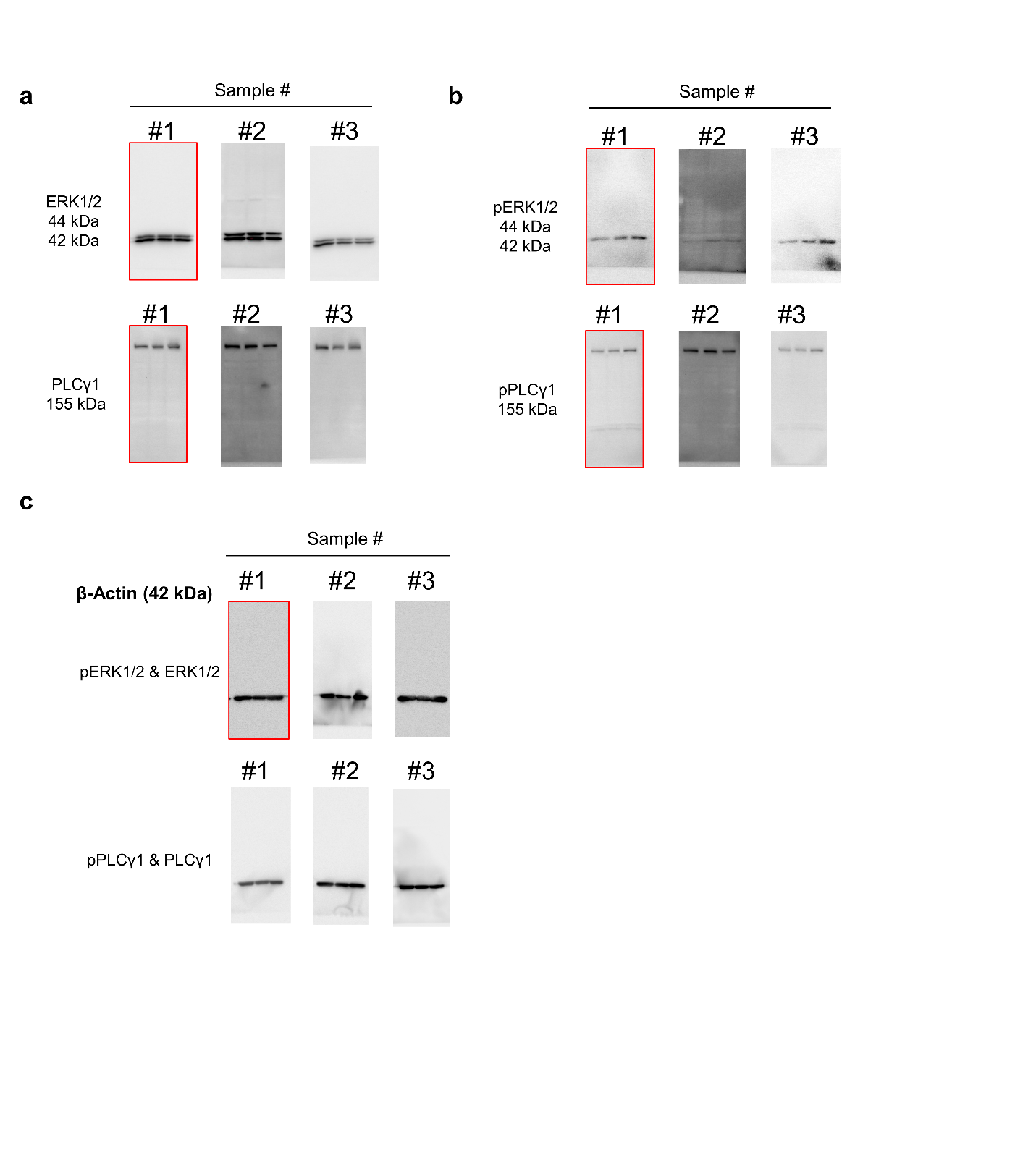

**Full western blots related to Fig. 4g.** Intracellular signaling pathway activation induced by the lead construct (linCAR-MS10-OX40ζ), quantified by normalizing phosphorylated PLCγ1 and ERK1/2 levels to their respective total protein levels (n = 3). (a) Blots of total PLCγ1 and ERK1/2. (b) Blots of phosphorylated PLCγ1 and ERK1/2. (c) Blots of β-actin. The blots outlined in red were selected as representative images for the figure.

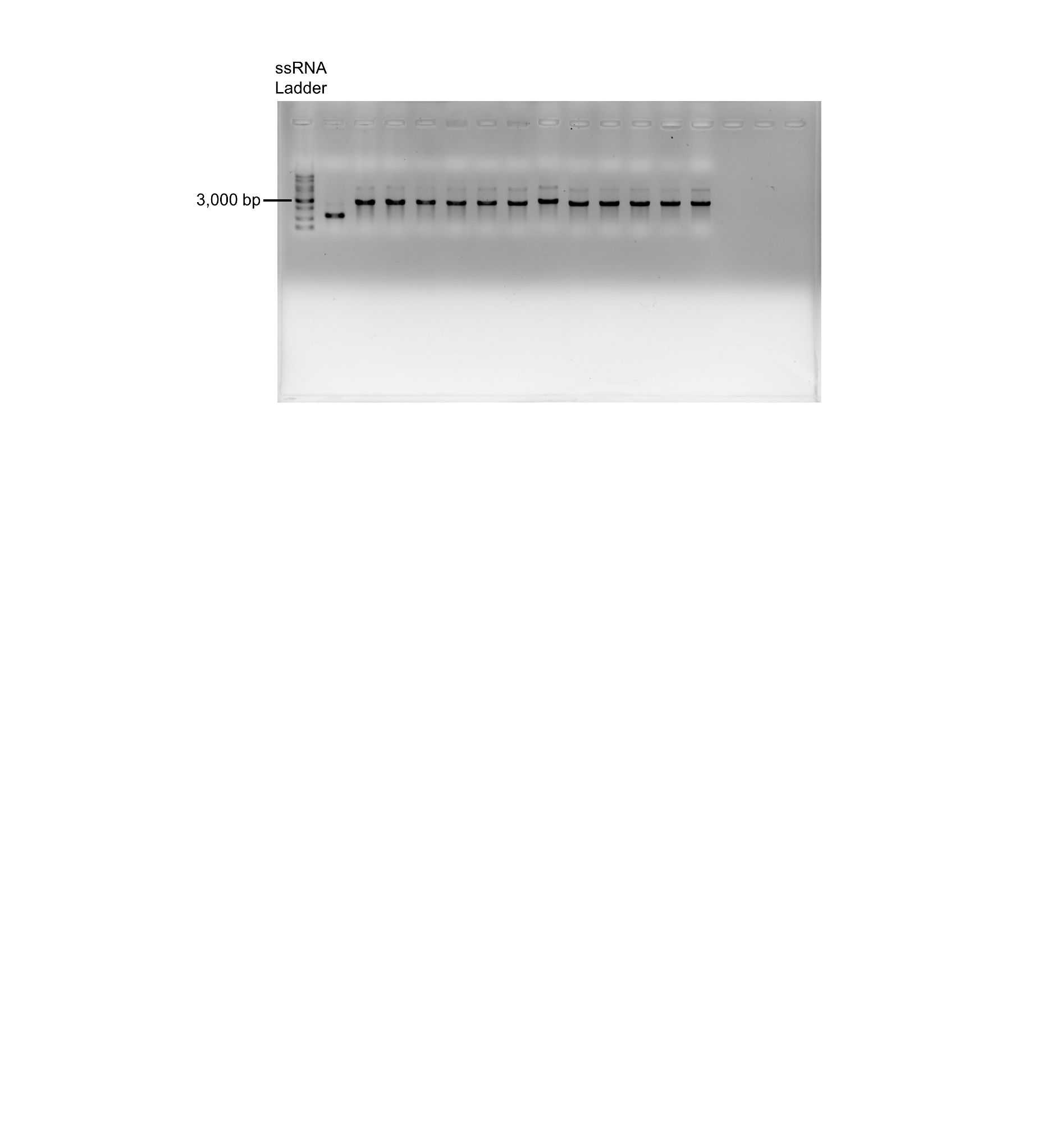

**Agarose gel image corresponding to Supplementary Figure 6a.** Agarose gel electrophoresis confirming the integrity and purity of *in vitro* transcribed linRNAs encoding various CAR constructs.

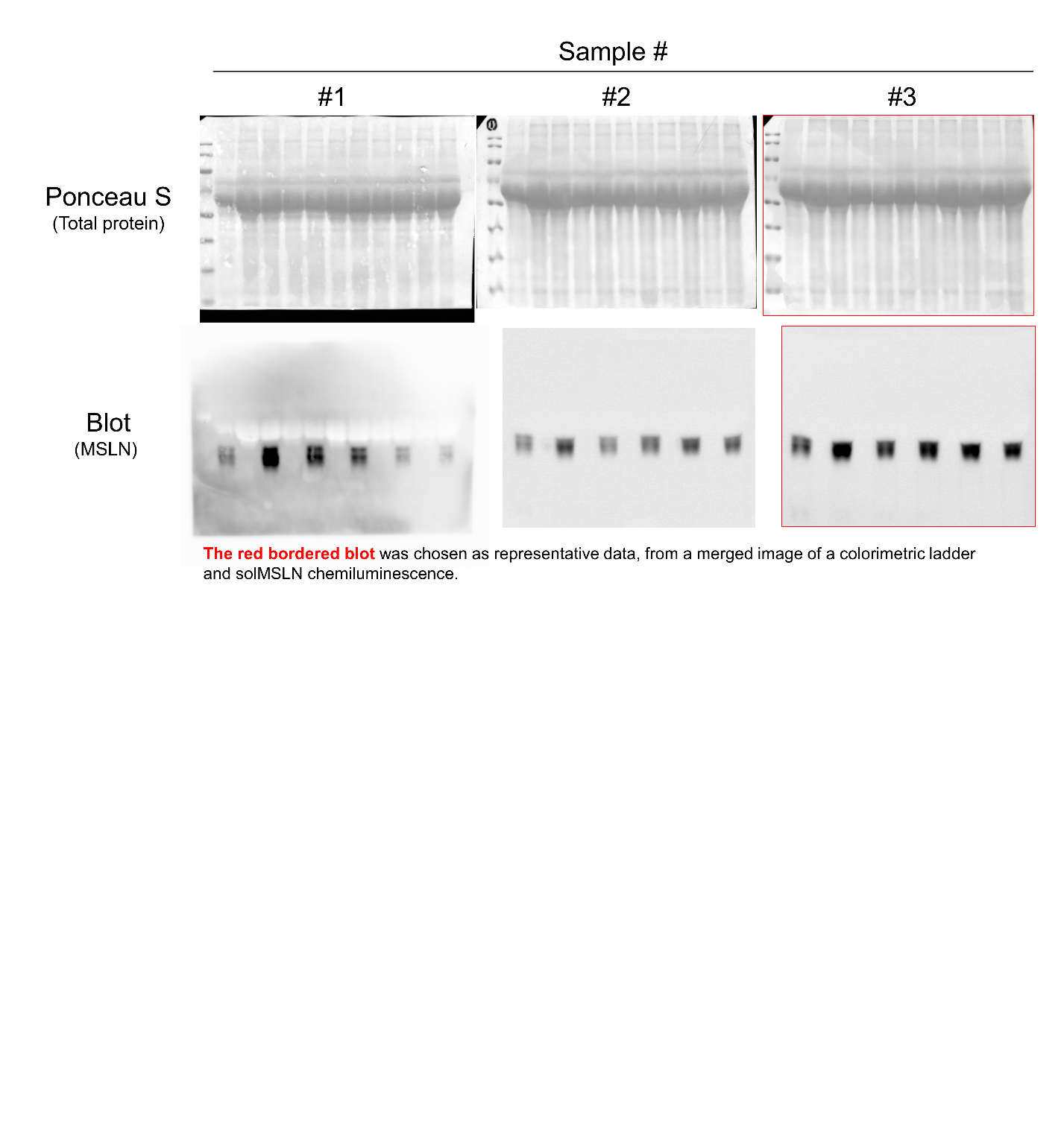

**Full western blots related to Supplementary Figure 12b.** Western blot analysis of solMSLN levels in conditioned media collected from Capan-2 cells alone, CAFs alone, or cocultures of Capan-2 cells with pancreatic CAFs or HPPFs. The blots **outlined in red were selected as representative images.**
